# Leveraging the BAF chromatin remodeling complex for targeted transcriptional rewiring in cancer

**DOI:** 10.64898/2026.03.30.715217

**Authors:** Dmitri Segal, Lisa Kainacher, Timo Muffel, Thomas M. Geiger, Severin Lechner, Danai Mavridi, Dimitrios-Ilias Balourdas, Maurice Leon Nelles, Caroline Schaetz, Stefan Knapp, Andreas C. Joerger, Radosław P. Nowak, Christian Steinebach, Georg E. Winter

**Affiliations:** CeMM Research Center for Molecular Medicine of the Austrian Academy of Sciences, Vienna, Austria; AITHYRA Research Institute for Biomedical Artificial Intelligence of the Austrian Academy of Sciences, Vienna, Austria; University of Greifswald, Institute of Pharmacy, Greifswald, Germany; Institute of Structural Biology, University Hospital Bonn, University of Bonn, Bonn, Germany; Goethe University, Institute of Pharmaceutical Chemistry, Frankfurt am Main, Germany; Structural Genomics Consortium (SGC), Buchmann Institute for Molecular Life Sciences, Frankfurt am Main, Germany

## Abstract

Chromatin accessibility is essential for maintaining the fidelity of gene regulation and is dynamically regulated by epigenetic enzymes that are often dysregulated in cancer. The most commonly mutated regulator is the modular, multi-subunit Brahma-associated factor (BAF) chromatin remodeling complex. Previous attempts to exploit BAF complex mutations as potential tumor vulnerabilities using loss-of-function approaches have shown limited clinical success. Here, we instead propose a gain-of-function (GOF) strategy and establish Transcriptional/Remodeling chemical Inducers of Proximity (TRIPs), a class of neomorphic molecules that recruit active BAF complexes to rewire an oncogenic repressor, B-cell lymphoma 6 (BCL6). TRIPs potently induce transcriptional de-repression and apoptosis in Diffuse Large B-cell Lymphoma (DLBCL), enabled by ternary complex formation between BCL6 and BAF. CRISPR knockout screening identifies the PBAF complex as an essential contributor to cellular TRIP efficacy. Finally, we demonstrate that TRIP induces chromatin enrichment of BAF at BCL6-bound sites, resulting in ATPase-dependent eviction of BCL6, and de-repression of pro-apoptotic BCL6 target genes. We establish BAF recruitment for targeted chromatin remodeling as a viable GOF pharmacological strategy for tackling diseases driven by aberrant gene repression.

## Introduction

Proper regulation of gene expression is essential for organismal development and faithful maintenance of cellular identity. This is a dynamic process that is highly regulated by altering physical access to DNA. Epigenetic regulators, including histone modifying enzymes and ATP-dependent chromatin remodelers, work in concert to modulate chromatin accessibility and maintain the fidelity of gene regulation^1–4^. Failure to faithfully maintain chromatin accessibility can occur due to mutations or imbalances in several master regulators of chromatin accessibility, leading to cancer development and progression^5^. One such master regulator is the SWItch/Sucrose Non-Fermentable (SWI/SNF) or Brahma-associated factor (BAF) complex^6^. BAF complexes are multi-subunit assemblies that drive ATP-dependent chromatin remodeling. They are composed of 29 modular subunits^7–9^ that give rise to three different subtypes: canonical BAF (cBAF), non-canonical BAF (ncBAF), and Polybromo-associated BAF (PBAF) complexes^10,11^. Across all cancer types, BAF complexes show the highest mutational burden amongst remodeling complexes, with more than 20% of all human tumors harboring a mutation in at least one of these subunits^12,13^. For this reason, the BAF complex has emerged as a pivotal therapeutic target in cancers with BAF alterations^14–19^, predominantly pursued via synthetic lethality strategies^8^. These include targeting paralog-deficiencies, such as ARID1B for ARID1A-mutant^20^ or SMARCA2 for SMARCA4-mutant cancers^21,22^, and subcomplex dependencies, such as ncBAF for SSX-SS18 fusion-positive synovial sarcoma^7,23^ and SMARCB1-null malignant rhabdoid tumor^7,24^. Intensive synthetic efforts to develop cancer therapeutics have yielded several drug candidates^25–28^. However, therapeutic disruption of BAF function has thus far not achieved clinical success^29–32^. Collectively, the prevalence of BAF mutations in cancer highlights the complex’s pathophysiological relevance and the importance and potential for therapeutic exploration.

Targeted tumor therapy with small molecules has traditionally focused on loss-of-function (LOF) approaches, including inhibition and degradation of cancer drivers^33–35^. However, chemical inducers of proximity (CIPs) have emerged as a conceptual framework for pharmacological modalities that can extend beyond LOF, including modulation of post-translational modifications and targeted re-localization^36,37^. In this regard, recent seminal work has established TCIPs (Transcriptional/epigenetic Chemical Inducers of Proximity)^38^ as a fundamentally different approach to targeted therapy. TCIPs are heterobifunctional molecules that rewire oncogenic transcription by inducing proximity between a tumor-associated transcription factor and a transcriptional regulator. In diffuse large B cell lymphoma (DLBCL), cells commonly overexpress BCL6, a transcriptional repressor that inhibits expression of pro-apoptotic genes^39–41^. Recruitment of potent transcriptional activators, including BRD4 and CDK9^38,42^, to BCL6 induces a gain-of-function (GOF) phenotype characterized by activating pro-apoptotic gene expression, resulting in rapid onset of cell death with efficacy/potency that far exceeds BCL6 LOF approaches. While an abundance of mutations has validated BAF as a disease relevant target, co-opting the BAF complex for GOF pharmacology remains largely unexplored.

Here, we introduce TRIPs (<u>T</u>ranscriptional/<u>R</u>emodeling chemical Inducers of <u>P</u>roximity), which chemically induce proximity and ternary complex formation between BCL6 and the catalytic BAF subunits SMARCA2/4. The resulting complex leads to rapid induction of apoptosis and potent cell viability effects specifically in BCL6-expressing DLBCL cell lines, despite sub-stoichiometric target engagement of the bifunctional molecule. Through CRISPR/Cas9 screening, we identify subunit and BAF subtype dependencies that are required for transcriptional de-repression and cell killing. Importantly, we demonstrate that targeting the ATP-dependent chromatin remodeling function of BAF to BCL6 loci leads to eviction and genome-wide redistribution of BCL6, thereby reactivating pro-apoptotic BCL6 target genes, including CDKN1A and ARID3A. In contrast to conventional BAF inactivating strategies, which rely on strong tumor-specific dependencies and near-complete target elimination, TRIPs harness the chromatin remodeling function of the BAF complex in a GOF approach to directly de-repress pro-apoptotic BCL6 target genes.

## Results

### TRIP1 rapidly and selectively kills DLBCL cells

To utilize the BAF complex in a GOF proximity induction mechanism, we focused on rewiring the transcriptional circuitry of BCL6 in DLBCL cells. We envisioned a strategy based on a bifunctional molecule, which binds to both BCL6 and the SMARCA2/4 bromodomain (BD), thereby recruiting the BAF complex as an effector to BCL6-bound loci, leading to an increase in chromatin accessibility, eviction of BCL6, and rapid activation of pro-apoptotic genes (**Figure 1a**).

**Figure 1.**
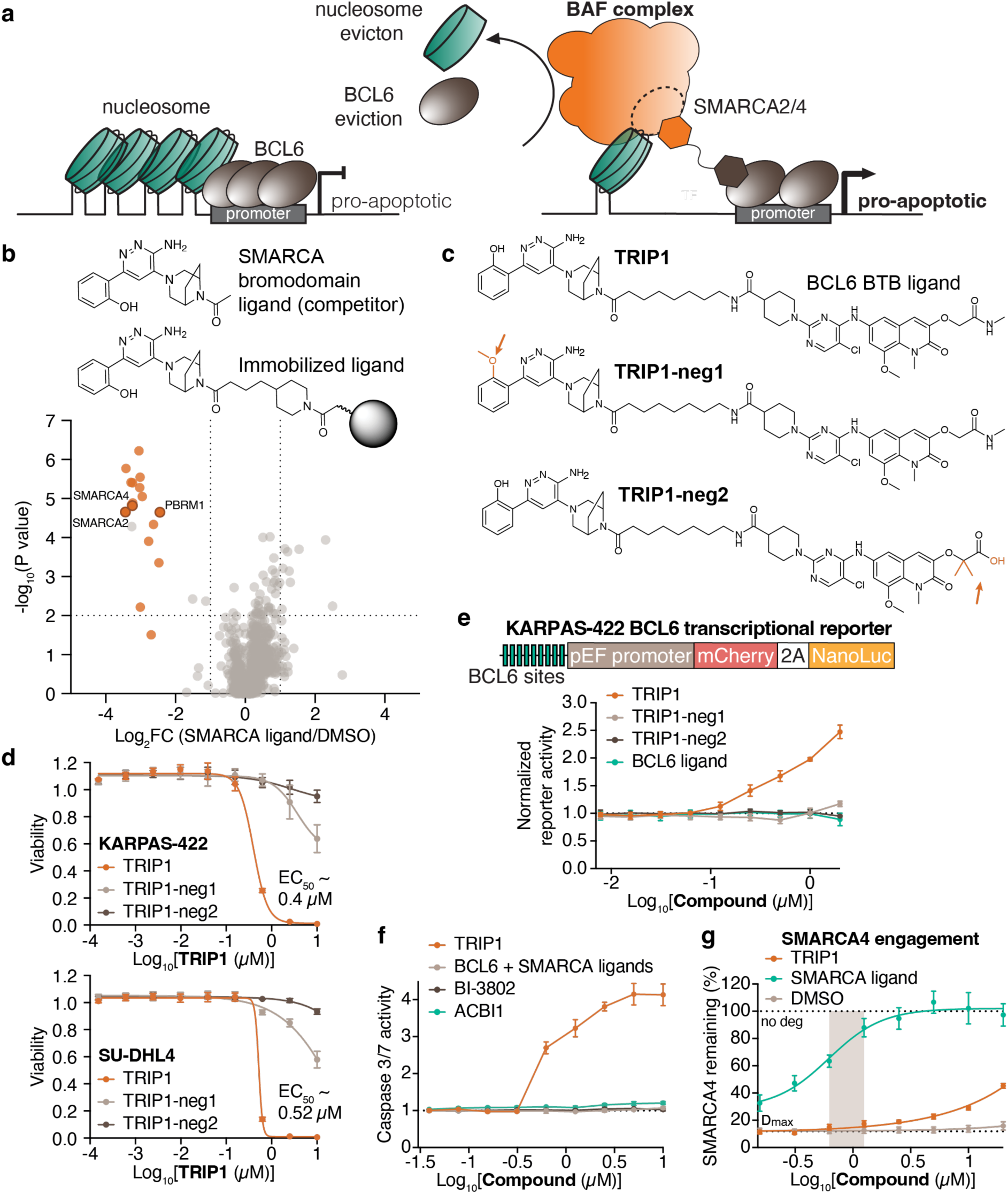
TRIP1 potently kills DLBCL cell lines with sub-stoichiometric engagement of SMARCA2/4. **a**, Schematic of chemically induced strategy for recruitment of the BAF complex (via SMARCA2/4) to BCL6. The BAF complex at BCL6 loci leads to eviction of nucleosomes and BCL6, increased chromatin accessibility, and de-repression of pro-apoptotic BCL6 target genes. **b**, Overview of chemoproteomic competition assay for SMARCA ligand characterization. SMARCA ligand with linker was immobilized on NHS-activated Sepharose beads and outcompeted with the SMARCA ligand (top). Each data point in the volcano plot (bottom) represents a dose-response score for a protein. The p-value indicates the significance of a dose-response according to the CurveCurator statistics pipeline, and the log2 fold change indicates the effect size of the dose-dependent depletion. Binding curves for highlighted BAF complex members in orange are shown in **Extended Data** Figure 1a. Protein with missing/imputed values in their dose-response curves were filtered out. Hits: Log_2_FC ≥ |1| and -log_10_P value ≥ 2 (dotted lines). **c**, Chemical structures of BCL6-SMARCA bifunctional compounds, TRIP1, TRIP1-neg1 (SMARCA non-binding), and TRIP1-neg2 (BCL6 non-binding). **d**, CellTiter-Glo cell viability assays in diffuse large B cell lymphoma (DLBCL) cell lines. KARPAS-422 (top) and SU-DHL4 (bottom) cells were treated with a dilution series of TRIP1 or negative controls for 72 hours. Viability with compound treatment is normalized to DMSO vehicle control; data represent mean ± SD, n = 3 independent replicates. **e**, BCL6 transcriptional reporter de-repression. KARPAS-422 cells expressing a BCL6 transcriptional reporter (top) were treated with a dilution series of TRIP1 or negative controls for 24 hours. Reporter activity is normalized to DMSO vehicle control; data represent mean ± SD, n = 3 independent replicates. **f**, CaspaseGlo 3/7 apoptosis detection. KARPAS-422 cells were treated with a dilution series of compounds for 16 hours. Caspase 3/7 activity is normalized to DMSO vehicle control; data represent mean ± SD, n = 3 independent replicates. **g**, SMARCA4 target engagement. Monoclonal HEK293T cells expressing endogenously tagged SMARCA4-HiBiT were pre-treated with a dilution series of TRIP1 or the SMARCA ligand for 2 hours, followed by co-treatment with 0.1 µM ACBI1 for 8 hours. SMARCA4-HiBiT levels are shown as a remaining fraction relative to DMSO vehicle control (no ACBI1 treatment). D_max_ represents the fraction of SMARCA4 remaining when ACBI1 is co-treated with DMSO. Gray bar indicates early cytotoxic/pro-apoptotic concentrations of TRIP1. Data represent mean ± SD, n = 6 independent replicates.

First, we aimed to determine whether a SMARCA ligand can be leveraged to recruit the entire BAF molecular machinery to BCL6 for targeted chromatin remodeling. Here, we used a functionally silent ligand with a terminal diazabicyclooctane handle that binds to the BDs of SMARCA2/4 and BD5 of PBRM1, while maintaining the enzymatic activity of the complex^43–47^. Using a competition-based chemoproteomics pulldown assay, we determined that the BD ligand can bind the intended targets, SMARCA2/4 and PBRM1, and more importantly, pull down most components of the BAF complex^11,48^, confirming that our strategy is capable of recruiting fully formed BAF complexes (**Figure 1b, Extended Data Figure 1a**, **Supplementary Table 1**). We also confirmed that this ligand binds to SMARCA4 BD and PBRM1 BD5 in a dose-dependent manner using an *in vitro* thermal shift assay (**Extended Data Figure 1b**). A central concern in using the SMARCA BD ligand was the potential for non-selective binding to other BD containing rewiring factors, such as BRD4^38^. Importantly, we found that the employed SMARCA BD ligand is selective and does not result in a thermal shift for BRD4 BD1 or bind BRD4 in the chemoproteomics experiment (**Figure 1b, Extended Data Figure 1b, Supplementary Table 1**), thereby establishing it as a bona-fide ligand for selective BAF complex recruitment.

To develop TRIPs, we connected this binder to an established BCL6 BTB domain binding ligand (BI-3812), which has previously been leveraged for TCIP synthesis^38,42,49^. We chose a bifunctional molecule design featuring a flexible carbon linker, hereafter referred to as TRIP1 (**Figure 1c**). We also synthesized corresponding negative control molecules with chemical alterations that mitigate SMARCA binding (TRIP1-neg1) and BCL6 binding (TRIP1-neg2). TRIP1 was able to robustly kill DLBCL cell lines, KARPAS-422 (EC_50_ ∼ 0.4 µM) and SU-DHL4 (EC_50_ ∼ 0.52 µM), after just 72 hours, unlike the negative control compounds or the combination of both ligands (**Figure 1d**, **Extended Data Figure 1c**). Furthermore, we observed that the cell viability phenotype is confined to BCL6 expressing cell lines (**Extended Data Figure 1d**), suggesting that non-BCL6 expressing cells would be spared from the effects of TRIP1, thereby confirming a functional dependence on BCL6.

To further characterize TRIP1-induced cellular phenotypes, we next tested whether TRIP1 can de-repress BCL6-regulated gene expression using a BCL6-repressed transcriptional reporter. Here, endogenous BCL6 in KARPAS-422 cells is constitutively repressing a highly active pEF promoter upstream from a reporter gene cassette. Only TRIP1, but not the negative control compounds or BCL6 ligand, was able to effectively de-repress this reporter (**Figure 1e**). In line with reporter activation, TRIP1 also potently induced caspase 3/7 activity after just 16 hours, indicative of apoptosis induction (**Figure 1f**), as expected from the anti-apoptotic circuitry of BCL6^38,42^. In contrast, degradation-based LOF strategies are either unable to reduce cell viability (BCL6 degrader, BI-3802^50,51^) or only trigger cytostatic phenotypes (SMARCA2/4 degrader, ACBI1^43^) in the DLBCL background (**Extended Data Figure 1e**). This is further evidenced by the inability of these degraders to induce apoptosis (**Figure 1f**). Importantly, TRIP1 does not induce degradation of SMARCA2/4 in endogenous HiBiT assays, confirming that the observed effects are not driven by the loss of BAF complex components (**Extended Data Figure 1f**).

LOF approaches generally require substantial target disruption, including high occupancy for inhibitors, or near-complete degradation to elicit phenotypic effects. We therefore sought to determine how much BAF recruitment is required to drive the gain-of-function phenotype. To approximate BAF engagement, we designed a competitive BAF degradation assay based on the principle that ligand binding to the SMARCA2/4 BD would rescue SMARCA2/4 degradation via a degrader that engages the same domain. Strikingly, we observed that, at concentrations which trigger reporter activity and apoptosis, TRIP1 only mildly rescued degradation of SMARCA2/4 by ACBI1. In contrast, the SMARCA ligand fully rescues degradation, suggesting that BAF-mediated rewiring drives phenotypes at sub-stoichiometric occupancy. Thus, these data indicate that TRIP1 only requires a small portion of available BAF complexes for efficacy, leaving most BAF complexes unperturbed (**Figure 1g**, **Extended Data Figure 1g**). Collectively, the ability of TRIP1 to de-repress gene expression, reduce cell viability, and induce apoptosis with sub-optimal target engagement strongly implies an event-driven, rather than occupancy-driven pharmacology.

### Functionality of TRIP1 depends on ternary complex formation

To determine whether TRIP1 induces ternary complex formation between BCL6 and SMARCA2/4, we developed a time-resolved fluorescence resonance energy transfer (TR-FRET) assay that measures proximity between the FITC-labeled BTB domain of BCL6 and the biotinylated BDs of SMARCA2/4 using streptavidin-terbium cryptate (SA-Tb) as the donor. We observed a clear TRIP1-induced dose-dependent increase of complex formation *in vitro*, which saturated at approximately 1 μM of compound (**Figure 2a**, **Extended Data Figure 2a**). Complex formation was only minimally induced with the TRIP1 negative control compounds. We also observed that proximity between SMARCA2/4 and BCL6 *in vitro* can be abrogated by increasing concentrations of the SMARCA or BCL6 ligands to compete for the binding pockets, again supporting TRIP1-induced ternary complex formation (**Figure 2b**).

**Figure 2.**
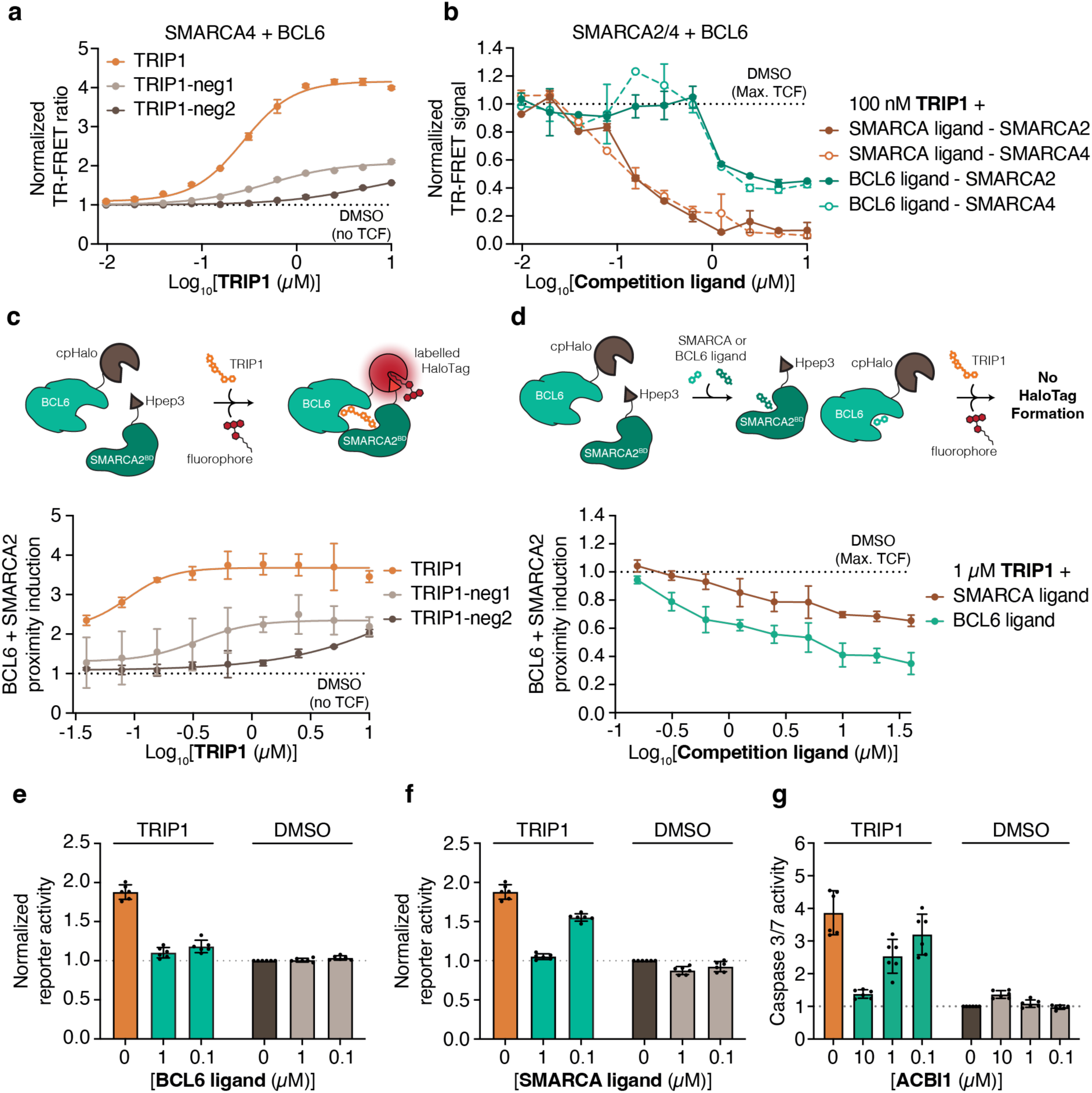
TRIP1 induces ternary complex formation between BCL6 and SMARCA2/4. **a**, *In vitro* TR-FRET ternary complex formation assay. The interaction between biotinylated SMARCA4 bromodomain (100 nM) and FITC-labeled BCL6 BTB domain (500 nM) was measured with increasing concentrations of TRIP1. The biotinylated SMARCA4 bromodomain was captured with streptavidin-terbium cryptate donor (SA-Tb, 2 nM), and TR-FRET signal was monitored upon complex formation. The resulting TR-FRET ratio was normalized to the DMSO vehicle control; data represent mean ± SD, n = 2 independent replicates. **b**, TR-FRET competition assay. The interaction between biotinylated SMARCA2/4 bromodomains (100 nM) and FITC-labeled BCL6 BTB domain (500 nM) was measured in the presence of 100 nM TRIP1 and increasing concentrations of BCL6 or SMARCA ligand. The resulting TR-FRET ratio was background subtracted and normalized to the DMSO vehicle control; data represent mean ± SD, n = 2 independent replicates. **c**, SplitHalo assay for BCL6-SMARCA2 interaction. Schematic of the in-cell splitHalo assay to probe induced protein-protein interactions (top). A HaloTag enzyme is split into two complementing parts, cpHalo and Hpep3 peptide, which only assemble into a functional self-labeling HaloTag enzyme when actively brought into proximity and supplied with TAMRA dye. HEK293T cells co-expressing BCL6-cpHalo and SMARCA2(BD)-Hpep3 were treated with a dilution series of TRIP1 or negative controls and incubated with compound and the covalent HaloTag dye TAMRA for 3 hours (bottom). Data is normalized to DMSO vehicle control, which corresponds to the baseline signal upon TAMRA addition; data represent mean ± SD, n = 3 independent replicates. **d**, SplitHalo competition assay for TRIP1-induced BCL6-SMARCA2 interaction. Schematic of the in-cell splitHalo competition assay to probe the inhibition of induced protein-protein interactions (top). Cells are pre-incubated with excess amounts of protein ligands to saturate binding pockets and prevent or reduce ternary complex formation. HEK293T cells co-expressing BCL6-cpHalo and SMARCA2(BD)-Hpep3 were pre-treated with a dilution series of SMARCA or BCL6 ligand for 30 minutes before adding 1 µM TRIP1 and TAMRA dye for 3 hours (bottom). Data is normalized to TRIP1 with DMSO vehicle control without ligand addition, corresponding to maximum ternary complex formation; data represent mean ± SD, n = 3 independent replicates. **e**, **f**, BCL6 transcriptional reporter competition. KARPAS-422 cells expressing a BCL6 transcriptional reporter were pre-treated with DMSO, the BCL6 ligand (**e**), or SMARCA ligand (**f**) for 8 hours, followed by co-treatment with 0.5 µM of TRIP1 for 24 hours. Reporter activity is normalized to DMSO vehicle control without TRIP1 co-treatment; data represent mean ± SD, n = 6 independent replicates. **g**, CaspaseGlo 3/7 apoptosis pre-degradation. KARPAS-422 cells were pre-treated with DMSO or SMARCA degrader (ACBI1) for 8 hours, followed by co-treatment with 1 µM of TRIP1 for 16 hours. Caspase 3/7 activity is normalized to DMSO vehicle control without TRIP1 co-treatment; data represent mean ± SD, n = 6 independent replicates.

We next asked whether TRIP1 induces ternary complex formation in cells by employing the SplitHalo interaction assay^52^. In brief, SMARCA2 BD was tagged with the small Hpep3 peptide, which has low intrinsic affinity to the larger cpHalo fragment fused to BCL6. Upon proximity between SMARCA2 and BCL6, the peptide Hpep3 complements cpHalo, and the fully functional self-labeling HaloTag enzyme is assembled, which undergoes covalent labeling in the presence of the TAMRA dye. Consistent with the TR-FRET assay, TRIP1 induced a dose-dependent increase in proximity between BCL6 and the SMARCA2 BD (**Figure 2c**). In cells, proximity was induced with concentrations as low as 40 nM and saturated at approximately 625 nM. In line with the *in vitro* TR-FRET data, TRIP1 induced cellular ternary complex formation was much more pronounced compared to negative control compounds with mitigated binding to either SMARCA2/4 (TRIP1-neg1) or BCL6 (TRIP1-neg2). Finally, we observed that proximity between BCL6 and SMARCA can be outcompeted in a dose-dependent manner with BCL6 or SMARCA ligands (**Figure 2d**), supporting TR-FRET data and reinforcing that ternary complex formation is induced by TRIP1 in cells.

Having established that TRIP1 induces proximity between BCL6 and SMARCA2/4, we next asked if ternary complex formation is functionally required. To this end, we assessed if individual ligands can outcompete TRIP1-induced cellular phenotypes. Indeed, we found that both the BCL6 and SMARCA ligands reduced or completely abolished BCL6 transcriptional reporter activation (**Figures 2e-f**). We next assessed apoptosis via caspase 3/7 activity, a downstream effect of TRIP1. Competition with a high concentration (10 µM) of the BCL6 ligand was able to reduce TRIP1-induced apoptosis, whereas reversible binding to SMARCA2/4 via the ligand was insufficient to maintain competition (**Extended Data Figures 2b-c**). This can be rationalized by the fractional occupancy required for TRIP1 to elicit its phenotypic effects. However, elimination of SMARCA2/4, via ACBI1-mediated pre-degradation, potently reduced caspase 3/7 activity in KARPAS-422 cells treated with TRIP1 (**Figure 2g**). Taken together, our data suggest that both BCL6 and SMARCA2/4 are functionally required for TRIP1-induced transcriptional de-repression and cell killing.

### CRISPR screen identifies PBAF complex as essential for TRIP1-induced cytotoxicity

To broadly interrogate the cellular and transcriptional machinery required for TRIP1-induced cytotoxicity, we aimed to identify the genetic determinants of TRIP1 function. To this goal, we performed a genome-wide viability-based CRISPR/Cas9 knockout screen in KARPAS-422 cells to identify genes whose loss confers resistance to TRIP1, indicating their relevance for TRIP1 efficacy. This led to the identification of all PBAF-specific subunits, BRD7, PBRM1, PHF10, and ARID2^11,48^, as critical dependencies for TRIP1-mediated cytotoxicity (**Figure 3a, Supplementary Table 2**). In contrast, sgRNAs targeting other BAF complex subunits were not significantly enriched in cells surviving TRIP1 treatment. The results of this screen can be partially explained by essentiality in KARPAS-422 cells and genetic redundancy (**Extended Data Figure 3a**). Unlike PBAF complex components, which are each necessary for full and functional complex assembly, loss of other BAF complex members, such as the core SMARCC1 subunit, can be compensated by redundant paralogous proteins^8,11^, and hence would not be identified in a CRISPR/Cas9 screen that only generates a single knockout in each cell. On the other hand, cells with knockouts of highly essential subunits, such as SMARCB1 and SMARCE1^53,54^, would drop out during the selection process independently of TRIP1 treatment. BRD9 is one subunit that was not enriched in the CRISPR screen, despite not being essential or redundant, possibly suggesting that the ncBAF complex is not involved in TRIP1-mediated toxicity in KARPAS-422 cells (**Figure 3a**, **Extended Data Figure 3a**). Finally, we identified RBM38 as a highly enriched hit in the CRISPR screen. RBM38, an RNA-binding protein, was shown to be functionally important for maintaining/stabilizing the CDKN1A (p21) transcript^55,56^, thereby implicating the cyclin-dependent kinase inhibitor as an important factor for TRIP1 function.

**Figure 3.**
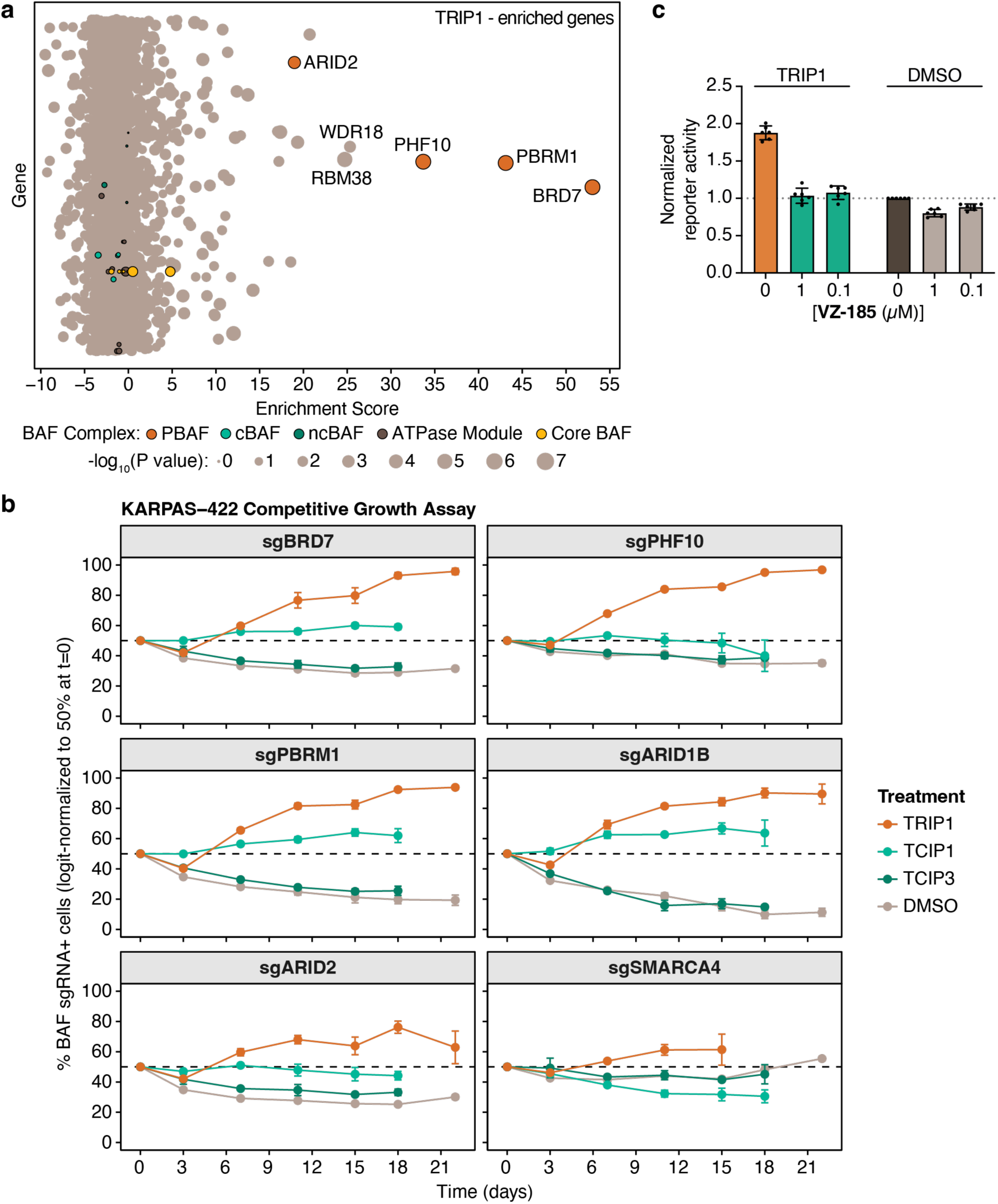
TRIP1 function is dependent on PBAF complex components. **a**, TRIP1 resistance CRISPR viability screen. KARPAS-422 iCas9 cells were mutagenized with a genome-wide sgRNA library and cultivated in the presence of DMSO or TRIP1 (400 nM) for 33 days. Gene-level enrichment score was calculated as -log10 (P value) × log2 (fold-change). BAF complex members are colored based on involvement in different sub-complexes or modules. n = 2 independent replicates. **b**, Competitive growth assay in KARPAS-422 cells. Control KARPAS-422 iCas9 cells expressing AAVS1-targeting sgRNA and BFP were mixed with KARPAS-422 iCas9 cells expressing BAF subunit-targeting sgRNAs and mCherry. Cell mixtures were treated with DMSO, TRIP1 (400 nM), BRD4 recruiting TCIP1 (4 nM), or p300/CBP recruiting TCIP3 (4 nM) and evaluated in regular intervals every 3 or 4 days via flow cytometry. Data were logit-transformed and normalized to 50% at day 0 to enable comparison across conditions. Data represent mean ± SD, n = 3 independent replicates. **c**, BCL6 transcriptional reporter pre-degradation. KARPAS-422 cells expressing a BCL6 transcriptional reporter were pre-treated with DMSO or BRD7/9 degrader (VZ-185) for 8 hours, followed by co-treatment with 0.5 µM of TRIP1 for 24 hours. Reporter activity is normalized to DMSO vehicle control without TRIP1 co-treatment; data represent mean ± SD, n = 6 independent replicates.

To validate the results of the viability-based CRISPR screen, we turned to a competitive growth assay where two populations of cells, one harboring negative control sgRNA targeting AAVS1 and one harboring sgRNA for BAF complex subunits, were mixed and allowed to grow for three weeks in the presence of DMSO or various TCIPs, including TRIP1. Here, we observed that in KARPAS-422 cells, loss of PBAF complex subunits, BRD7, PBRM1, PHF10, or ARID2, provided a clear survival advantage in the presence of TRIP1 (**Figure 3b**). Importantly, these survival advantages largely did not manifest in cells treated with BRD4 and CBP/p300 recruiting TCIPs, TCIP1, and TCIP3, respectively^38,49^, suggesting that strong PBAF dependency is specific to the SMARCA2/4 recruiting TRIP1. We observed that loss of the PBAF subunits, BRD7, PHF10, and PBRM1, also provided a survival advantage in SU-DHL4 cells, albeit weaker than what was observed with KARPAS-422 cells (**Extended Data Figure 3b**). Finally, we wondered whether pre-degradation of the PBAF complex member BRD7 might prevent TRIP1-induced phenotypes. Indeed, chemical ablation of BRD7/9 with the BRD7/9 PROTAC VZ-185^57^ completely prevented activation of the BCL6 transcriptional reporter (**Figure 3c**), supporting the mechanistic role of the PBAF complex in TRIP1 activity.

Competitive growth assays can also resolve functional dependencies even if the gene knockout itself reduces cellular fitness. Leveraging this, we monitored the potential survival advantage of ARID1B knockout in KARPAS-422 cells, which harbor a deletion mutation in ARID1A and are selectively dependent on its paralog ARID1B for survival^20,54,58^ (**Extended Data Figure 3a**). In support of this, cells expressing sgRNAs targeting ARID1B dropped out compared to cells expressing control sgRNAs targeting AAVS1 when cells were control-treated (DMSO). In contrast, cells displayed a survival benefit under treatment challenge with TRIP1 (**Figure 3b**). This proved specific for the ARID1A mutant background in KARPAS-422, as SU-DHL4 cells, which possess wild-type ARID1A, were neither affected by ARID1B knockout, nor did they receive any survival advantage under TRIP1 treatment (**Extended Data Figure 3b**). Collectively, these observations imply that the canonical BAF complex is also involved in TRIP1-mediated toxicity.

### TRIP1 upregulates BCL6 target genes involved in apoptosis and cell-cycle arrest

To investigate how TRIP1 globally influences gene expression, we employed RNA-sequencing (3’ fingerprinting) in KARPAS-422 cells (**Extended Data Figures 4a-b**, **Supplementary Table 3**). Here, we observed dose- and time-dependent transcriptional changes in the presence of TRIP1, which were not observed with the negative control compounds (**Figures 4a-b**, **Extended Data Figure 4c**). Consistent with the hypothesis that TRIP1 specifically targets BCL6-regulated genes, a Gene Set Enrichment Analysis (GSEA) demonstrated strong enrichment of BCL6 targets genes^38,42^ (**Figure 4c**, **Extended Data Figure 4d, Supplementary Table 3**). This enrichment within the TRIP1 upregulated genes was clearly more robust with the active compound relative to negative controls. Moreover, a Landscape In Silico deletion Analysis (LISA)^59^ predicted that BCL6 is the most likely transcriptional regulator of the differential expressed genes following TRIP1 treatment (**Figure 4d, Supplementary Table 3**), suggesting that TRIP1 indeed functions by rewiring BCL6 and regulating the expression of BCL6 target genes.

**Figure 4.**
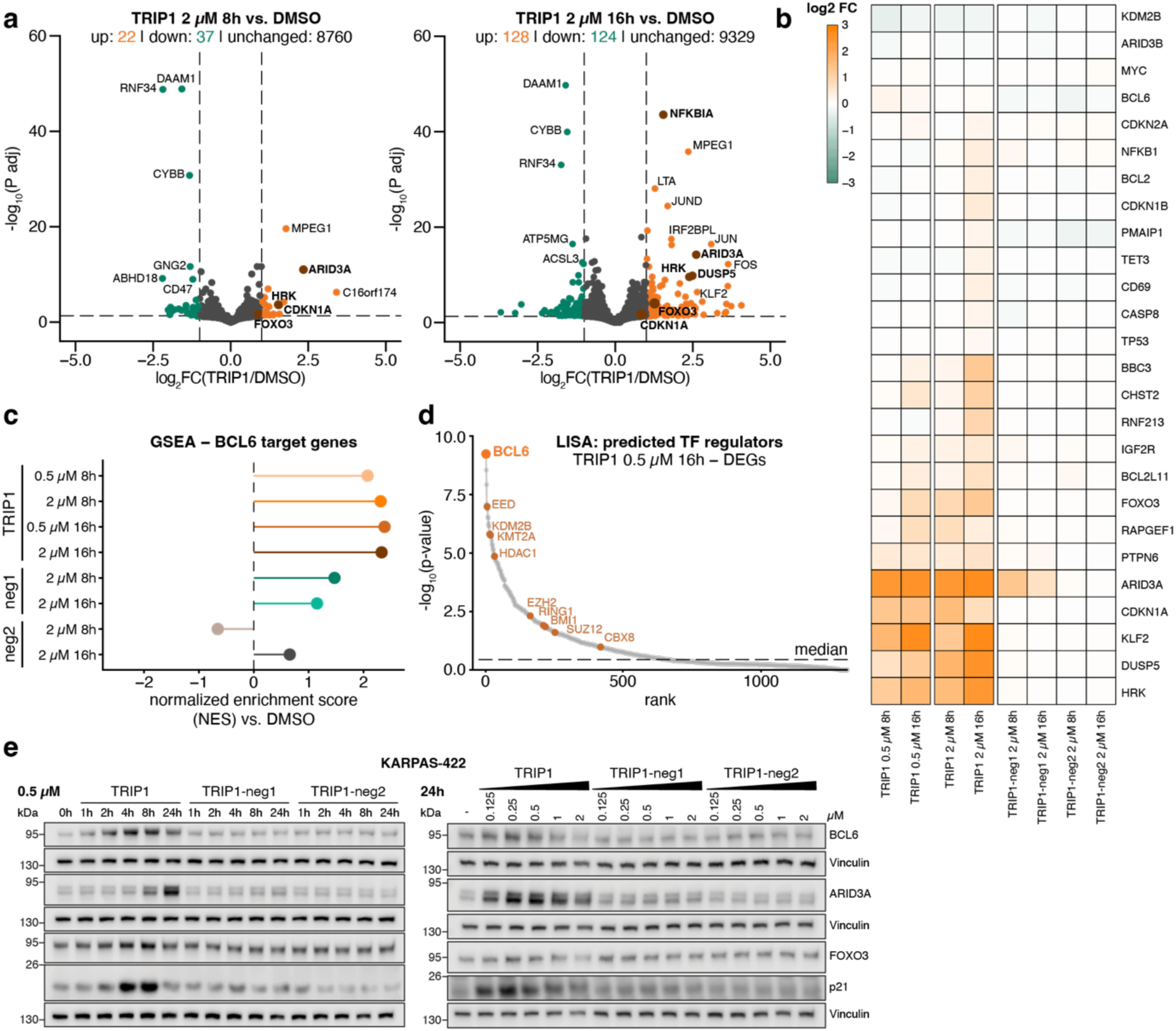
TRIP1 de-represses transcription of BCL6 target genes. **a**, Genes up- and down-regulated after TRIP1 treatment. 3’ RNA fingerprinting in KARPAS-422 cells after 8h (left) or 16h (right) treatment with 2 µM TRIP1. BCL6 target genes involved in apoptosis, B-cell differentiation, and cell-cycle are highlighted in brown/bold. Hits: Log_2_FC ≥ |1| and P adj < 0.05 (dotted lines). **b**, Gene expression changes (3’ RNA fingerprinting) in KARPAS-422 cells of selected BCL6 target genes after treatment with TRIP1 or negative control compounds at two different time points and concentrations. **c**, Lollipop plot summarizing gene set enrichment analysis (GSEA) of BCL6 target genes and genes shown to be regulated by BCL6 targeting TCIPs in **Extended Data** Figure 4d. Normalized enrichment scores are plotted for each condition compared to DMSO. **d**, LISA analysis of differentially expressed genes (DEGs) after TRIP1 treatment (16h, 0.5 µM). BCL6 (orange) is the top predicted transcriptional regulator of DEGs after TRIP1 treatment. Other transcriptional repressors with a high score are highlighted in brown. For **a-d**, differential gene expression analysis was performed using DESeq2. Log₂ fold changes were shrunk using the apeglm method. P-values were obtained from the Wald test and adjusted for multiple testing using the Benjamini-Hochberg procedure. Genes with an adjusted p-value (padj) < 0.05 and |log₂FC| > 1 are highlighted as differentially expressed; n= 3 independent replicates. **e**, Validation of TRIP1-induced expression changes at the protein level. Time-resolved (left) and dose-resolved (right) expression changes of BCL6, ARID3A, FOXO3 and p21 (CDKN1A) in KARPAS-422 cells. Loading controls were performed per gel. Western blots are representative of two independent replicates.

To understand how TRIP1 might be exerting phenotypic changes that result in reduced cell viability and death of DLBCL cells, we focused on a small subset of proteins involved in cell-cycle arrest and apoptosis at the mRNA and protein levels. Known BCL6 target genes, including the proapoptotic BH3-only protein BIM (BCL2L11), the cell-cycle arrest protein p21, and the pro-apoptotic transcriptional regulator FOXO3 were specifically upregulated by TRIP1^39,60–62^, independent of p53^63^, which is inactivated/lost in KARPAS-422 cells^64^ (**Figure 4b**). Other relevant upregulated anti-proliferative factors include a pro-apoptotic protein, Harakiri (HRK), a phosphatase and direct negative regulator of ERK1/2 signaling, DUSP5, and a B-cell differentiation/pro-apoptotic factor, ARID3A^65–68^. At the protein level, we observed dose- and time-dependent increases in p21, FOXO3, and ARID3A levels in KARPAS-422 and SU-DHL4 cells, which were not observed with the negative control compounds (**Figure 4e**, **Extended data Figure 5a**). Additionally, BCL6 levels increased in a dose- and time-dependent manner with TRIP1 treatment, either through a previously described auto-regulatory feedforward loop or increased protein stability^38^. Interestingly, the response to TRIP1 can be separated into different phases, with p21 and FOXO3 showing upregulation as early as 2-4 hours, highlighting their role as critical early response genes (**Figure 4e**, **Extended data Figure 5a**). In contrast, ARID3A upregulation occurs at 8-24 hours, and pro-apoptotic caspase 3 cleavage also follows a late response pattern at 24 hours (**Extended data Figures 5b-c**), although activity can be measured as early as 16 hours (**Figure 1f**). In general, TRIP1 induces expression of a specific set of BCL6 target genes with critical anti-proliferative and pro-apoptotic functions, thereby triggering cell death of DLBCL cell lines.

### ATPase-dependent chromatin remodeling by the BAF complex evicts BCL6 upon TRIP1 treatment

To determine whether the specific upregulation of BCL6 target genes coincides with BAF recruitment to BCL6-bound loci, we performed CUT&RUN chromatin profiling. We treated KARPAS-422 cells with TRIP1 at multiple time points, profiled genome-wide BCL6 and SMARCA4 binding, and assessed occupancy changes after treatment. Here, we found that BCL6 was lost from bound sites after 4 hours and 8 hours despite increased BCL6 protein abundance following TRIP1 treatment (**Figure 4e**, **Figure 5a**). Concurrently, we observed a stable increase of SMARCA4 at BCL6 peaks, indicating that TRIP1 can evict BCL6 from chromatin while increasing BAF loading at these loci. The increase of SMARCA4 was accompanied by an increase of histone 3 lysine 27 acetylation (H3K27ac), a marker for active chromatin regions^69^, at BCL6 loci (**Extended Data Figure 6a**). Importantly, TRIP1 did not lead to a loss of SMARCA4 at SMARCA4 peaks (**Extended Data Figure 6b**). To better understand the genome-wide binding patterns within and around coding regions and their change upon TRIP1 treatment, we checked BCL6 and SMARCA4 binding, as well as H3K27ac deposition across all annotated human genes. K-means clustering identified three gene groups with high, intermediate, or low BCL6 occupancy. The smallest cluster, BCL6_high_ genes, showed a pronounced loss of BCL6 following TRIP1 treatment, accompanied by increased SMARCA4 binding and H3K27ac deposition. In contrast, BCL6_mid_ genes exhibited more modest changes, while BCL6_low_ genes remained unchanged (**Extended Data Figure 6c**). Together, these results indicate that the extent of TRIP1-induced chromatin remodeling scales with the initial level of BCL6 occupancy.

**Figure 5.**
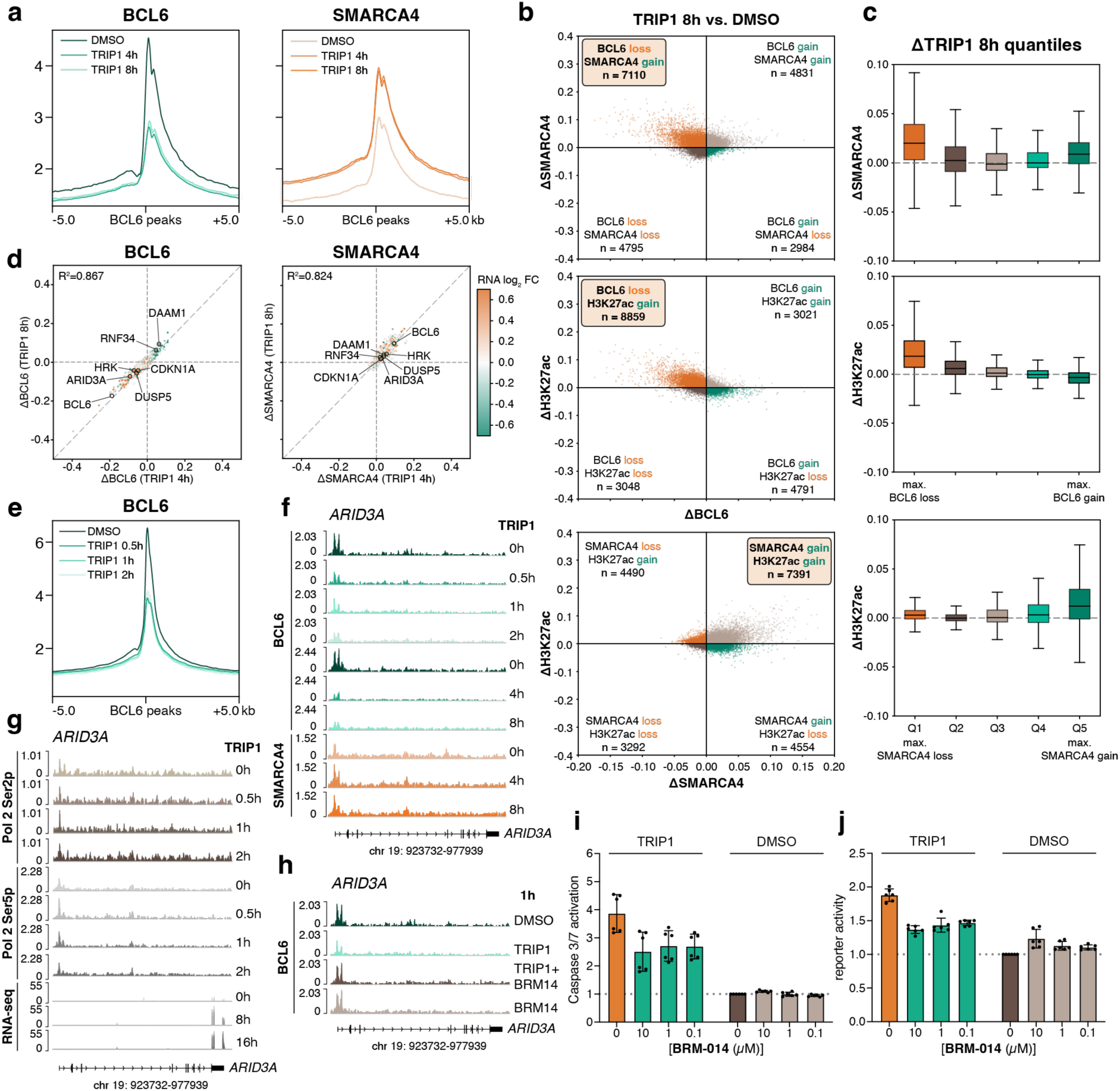
TRIP1 evicts BCL6 from chromatin via ATPase-dependent BAF activity. **a,** CUT&RUN binding profiles at BCL6 peaks. Binding profiles of BCL6 (left) and SMARCA4 (right) at BCL6 peaks for DMSO/TRIP1 (2 µM) treated cells after 4 and 8 hours. BCL6 peaks were called from DMSO-treated KARPAS-422 samples. Profiles were centered on peaks and extended 5 kb upstream and downstream of the peak location. **b**, Correlation between BCL6 and SMARCA4 (top), BCL6 and H3K27ac (middle), or SMARCA4 and H3K27ac (bottom) signal changes on gene bodies upon TRIP1 (8h, 2 µM) treatment compared to DMSO. Single dots represent hg38 genes. **c,** Relationship between TRIP1-induced BCL6/SMARCA4 binding change and H3K27 acetylation or SMARCA4 binding. Genes were ranked based on their differential binding of BCL6 (top) and SMARCA4 (bottom) after TRIP1 treatment (8h, 2 µM) and segmented into respective 20% quantiles. SMARCA4 (top) and H3K27ac (middle) changes at genes per BCL6 quintile. H3K27ac change (bottom) at genes segmented by SMARCA4 quintiles. After overall association was assessed by Kruskal-Wallis was successfully, pairwise comparisons were made by Dunn’s post-hoc test with Benjamini-Hochberg FDR correction. **d**, Relationship of TRIP1-induced gene expression changes with BCL6/SMARCA4 binding changes after 4 and 8 hours of TRIP1 treatment. Pearson correlation coefficient and R^2^ were calculated. Dots represent single genes colored by their gene expression change upon TRIP1 treatment (2 µM after 16 hours vs. DMSO). Selected differentially expressed genes and BCL6 are highlighted. For **b-d**, Normalized CUT&RUN signal on gene bodies ±3 kb up and downstream to include regulatory regions was calculated for TRIP1 and DMSO. The scores were subtracted to calculate differential binding. **e**, CUT&RUN binding profiles at BCL6 peaks. Binding profiles of BCL6 at BCL6 peaks after short, low-dose TRIP1 (1 µM) treatment up to 2h. BCL6 peaks were called from DMSO-treated KARPAS-422 CUT&RUN samples. Profiles were centered on peaks and extended 5 kb upstream and downstream of the peak location. **f-h**, Genome tracks of the ARID3A gene locus. Time-resolved (**f**) BCL6 and SMARCA4 or (**g**) RNA Pol II serine 2/5 phosphorylation signal is computed along the gene locus to infer transcriptional dynamics. For **g**, Below, RNA-seq reads are mapped to the gene locus. **h**, BAF ATPase-dependent BCL6 eviction after 1 hour of DMSO or TRIP1 (1 µM) co-treatment with BRM-014 (1 µM). BCL6signal is computed along the ARID3A gene locus. Exon position and genome location are indicated below the genome tracks. All CUT&RUN data is from two merged independent replicates. **i**, CaspaseGlo 3/7 apoptosis ATPase pre-inhibition. KARPAS-422 cells were pre-treated with DMSO or SMARCA2/4 ATPase inhibitor (BRM-014) for 8 hours, followed by co-treatment with 1 µM of TRIP1 for 16 hours. Caspase 3/7 activity is normalized to DMSO control without TRIP1 co-treatment; data represent mean ± SD, n = 6 independent replicates. **j**, BCL6 transcriptional reporter ATPase pre-inhibition. KARPAS-422 cells expressing a BCL6 transcriptional reporter were pre-treated with DMSO or SMARCA2/4 ATPase inhibitor (BRM-014) for 8 hours, followed by co-treatment with 0.5 µM of TRIP1 for 24 hours. Reporter activity is normalized to DMSO vehicle control without TRIP1 co-treatment; data represent mean ± SD, n = 6 independent replicates.

To confirm that TRIP1-induced global chromatin remodeling also occurs at individual genes, we correlated changes in gene body-bound BCL6 with SMARCA4 recruitment and H3K27ac deposition at single gene loci. As expected, we observed that genes that lost BCL6 preferentially gained SMARCA4 and H3K27ac signal after 8 hours. Similarly, genes with increased BAF occupancy upon TRIP1 treatment showed increased deposition of H3K27ac (**Figure 5b, Supplementary Table 4**). Next, we ranked genes by their decrease in BCL6 signal upon TRIP1 treatment. Genes with the greatest BCL6 loss upon TRIP1 treatment (quantile Q1) showed the greatest increase in SMARCA4 and H3K27ac signal. Accordingly, genes with increased BAF occupancy showed a simultaneous increase in H3K27ac after 8 hours (**Figure 5c**). To determine whether the changes at 4 and 8 hours reflect stable or dynamic changes in BCL6 and BAF binding, we correlated BCL6 and SMARCA4 changes across individual genes at both time points. The strong correlation (BCL6: R^2^=0.867, SMARCA4: R^2^=0.824) confirmed that BCL6 loss and SMARCA4 gain on single genes persisted for at least 8 hours. Accordingly, genes exhibiting BCL6 loss and SMARCA4 gain were largely upregulated (**Figure 5d**). In general, these gene-level changes confirmed that global shifts reflected e8ective remodeling occurring at single gene loci after TRIP1 treatment.

To gain further insight into the early chromatin changes upon TRIP1 treatment, we next profiled KARPAS-422 cells after short TRIP1 incubations. Remarkably, the loss of BCL6 from BCL6 sites occurred after only 30 minutes (**Figure 5e**), while overall BAF occupancy remained stable (**Extended data Figure 7a)**. In agreement, BCL6 was also rapidly lost from the locus encoding the ARID3A protein (**Figure 5f**), which was found to be strongly upregulated by TRIP1 (**Figures 4a**-**b**). This was accompanied by increased levels of RNA Pol II serine 5 phosphorylation within 30 minutes (**Figure 5g**). The rapid BCL6 loss was also evident at the BCL6 locus (**Extended data Figure 7b**). Clustering of BCL6-bound loci by binding strength and pattern, revealed a similar loss of BCL6 across all motifs within 30 minutes (**Extended data Figures 7c**-**d)**, consistent with previous reports highlighting that BAF-mediated eviction of PRC1/2 and repressive histone marks occurs within just 5-20 minutes^70^.

Given the core function of BAF as a chromatin remodeling complex, we wondered if its ATPase activity, which catalytically drives nucleosome remodeling^71^, is responsible for the observed chromatin changes. To this end, we co-treated KARPAS-422 cells with the SMARCA2/4 ATPase inhibitor BRM-014^72^ and TRIP1, and profiled chromatin changes compared to single compound treatments. We first profiled the ARID3A locus, which is strongly regulated by TRIP1 (**Figure 4**, **Figures 5f-g**), and observed that TRIP1-induced BCL6 eviction was indeed fully abrogated by BAF ATPase co-inhibition (**Figure 5h**), while SMARCA4 enrichment at the locus remained unaffected (**Extended data Figure 8a).** Next, we clustered all BCL6 binding sites according to their binding pattern (**Extended data Figure 8b**). Clusters 1 and 2 represented sites with strong BCL6 binding spanning the entire 10 kb regions. At these binding sites, BRM-014 co-treatment highly attenuated TRIP1-induced BCL6 loss. Sites less densely occupied by BCL6, corresponding to clusters 3-5, showed no change upon ATPase co-inhibition (**Extended data Figure 8c**). These data suggest that BCL6 eviction is mediated by the ATP-dependent chromatin remodeling function of BAF.

According to this working model, ATPase function drives the cytotoxic phenotype observed after TRIP1 treatment. Indeed, we found that both apoptosis induction (**Figure 5i**) and transcriptional activation of the BCL6 reporter (**Figure 5j**) were partially reduced, but not fully abrogated, by BRM-014 pre-treatment. This indicates that ATPase-dependent chromatin remodeling via TRIP1-recruited BAF complexes can account for some, but not all, observed TRIP1 effects in DLBCL cells and that other transcriptional changes might be independent of the catalytic turnover, but potentially explained by steric interference or co-activator recruitment.

## Discussion

Due to its high mutational burden across several malignancies, several subunits of the BAF complex have emerged as attractive cancer targets. Until now, pre-clinical and clinical BAF-targeting strategies were designed to inhibit or degrade BAF subunits to elicit an anti-tumor effect^43,46,57,72–75^. Here, we demonstrated a reverse strategy that leverages the chromatin remodeling function of BAF to rewire the chromatin environment in tumor cells in a targeted and programmable manner. Using the chemical inducer of proximity TRIP1 in BCL6 expressing DLBCL cell lines, we found that BAF complex recruitment could effectively remodel BCL6-bound chromatin, leading to de-repression of pro-apoptotic genes and activation of cell death.

Collectively, our data argues that the potency and kinetics of these molecules arise from direct transcriptional rewiring that is achieved by event-driven GOF pharmacology that relies on proximity induction rather than maintainance of BCL6 occupancy. These observations differentiate TRIP1 from RIPTACs (Regulated Induced Proximity Targeting Chimeras), another heterobifunctional modality with promising clinical impact^76^. RIPTACs represent a form of occupancy-driven LOF pharmacology and link an overexpressed tumor-specific protein, such as BCL6, to an essential protein, such as BRD4, thereby sequestering and inactivating the latter^77^. However, several critical points argue against occupancy-driven BAF disruption via TRIP1. Unlike SMARCA2/4 or BCL6 degradation, which also represents LOF pharmacology, TRIP1 can rapidly induce toxicity and apoptosis. Moreover, TRIP1 elicits potent cytotoxic/pro-apoptotic responses at extremely low levels of SMARCA2/4 engagement. This defies the proposed mechanism of RIPTACs which rely on saturated and stable ternary complex formation to disable an essential protein. Finally, TRIP1 induces BAF enrichment on chromatin, which coincides with increased active chromatin marks and ATPase-dependent BCL6 eviction, a process that is incompatible with occupancy-driven LOF.

Using unbiased genetic screens, we here provide the first in-depth look at genetic determinants of CIPs that rewire gene control circuits. Genome-wide CRISPR/Cas9 screening identified a specific dependency for all subunits of the PBAF complex in TRIP1 function. Although the observed dependencies may reflect subunit redundancy or essentiality-related dropout in KARPAS-422 cells, rescue of TRIP1-induced reporter activation by a BRD7/9 degrader suggests a prominent role for PBAF in BCL6 rewiring. Importantly, TRIP1 can potently induce apoptosis in KARPAS-422 cells, despite harboring a heterozygous deletion in ARID1A^53,54,58^. This exemplifies the fidelity of our approach in BAF mutant genetic backgrounds. Our GOF strategy bypasses the need to exploit synthetic lethalities, which arise from paralog deficiencies, such as ARID1B in ARID1A-mutant cells^20^. Ultimately, these findings demonstrate that TRIP1 function depends on the BAF complex subtypes available in a given genetic context, with BAF mutational status and expression levels shaping the chromatin rewiring response across cell lines.

Using genomic profiling, we demonstrate robust enrichment of SMARCA4 at BCL6 target genes. Importantly, this is accompanied by increased active histone marks at key BCL6 targets, resulting in elevated gene expression. We provide the first insight into CIP-induced BCL6 chromatin changes, which revealed prominent BCL6 eviction with TRIP1, pointing to a unique mechanism of action, mediated by direct chromatin remodelling, that is distinct from TCIPs^38,42,49^. Moreover, we show that TRIP1-triggered eviction requires ATPase-dependent chromatin remodeling. Partial rescue of TRIP1-induced apoptosis and reporter de-repression upon ATPase inhibition suggests that additional mechanisms beyond nucleosome eviction likely contribute to its activity. Indeed, partial nucleosome unwrapping occurs despite ATPase inhibition^71^. Additionally, interaction with other transcriptional co-activators could mediate nucleosome remodeling-independent BCL6 de-repression^78^.

The successful rewiring of BCL6 circuits in DLBCL via BAF complex recruitment and targeted chromatin remodeling demonstrates the power of leveraging BAF functionality, as opposed to its depletion. Importantly, previous studies showed that synthetic recruitment of the BAF complex was sufficient for de-repression of an engineered PRC2-repressed reporter locus^70^. This highlights the versatility of BAF complex recruitment to achieve localized chromatin remodeling and potent eviction of chromatin-bound repressors beyond BCL6, such as PRC2. Future studies will illuminate the reach of co-opting the BAF complex, as well as other chromatin remodeling factors, to activate genetic loci that are aberrantly repressed or otherwise reduced in expression to ameliorate or correct disease states. While the focus of this study is on cancer, other diseases, including haploinsu8iciencies, metabolic enzyme deficiencies, or neurodegenerative diseases, could be actionable by chemical rewiring of chromatin remodeling complexes.

## Supporting information

Proteomics

Cut&Run

RNASeq

CRISPR

Flow Cytometry Gating

Chemical Synthesis

## Acknowledgements

The Winter lab was and is supported by funding from the European Research Council (ERC) under the European Union’s Horizon 2020 research and innovation program (grant agreement 851478 and 101170771), as well as by funding from the Austrian Science Fund (FWF, projects P7909, P36746 and P5918723) and the Vienna Science and Technology Fund (WWTF, project LS21-015). D.S. is supported by MSCA Postdoctoral Fellowship 2023 (HORIZON-MSCA-2023-PF-01, proposal 101153103). L.K is supported by Boehringer Ingelheim Fonds PhD fellowship (2025). S.L is supported by an EMBO Postdoctoral Fellowship (ALTF 236-2024). Work in the Steinebach lab is supported by funding from the German Research Foundation (DFG, 552374678). T.M. is supported by a scholarship from the Studienstiftung des deutschen Volkes. Work in the Nowak lab is supported by funding from the German Research Foundation (DFG, 552374678). R.P.N. is a member of the excellence cluster ImmunoSensation3 funded by the German Research Foundation (DFG) under Germany’s Excellence Strategy (EXC2151, 390873048). Work in the Joerger and Knapp labs is supported by funding from the German Cancer Aid grant TACTIC (project 70115201), which is part of the preclinical cancer drug development network (preCDD). D.M. is supported by an Onassis Foundation fellowship (F ZU 049-1/ 2024-2025). D.M., D-I.B., S.K., and A.C.J. are grateful for support from the Structural Genomics Consortium (SGC), a registered charity (No:1097737) that received funds from Bayer AG, Boehringer Ingelheim, Bristol Myers Squibb, Genentech, Genome Canada through Ontario Genomics Institute, [OGI-196], EU/EFPIA/OICR/McGill/KTH/Diamond Innovative Medicines Initiative 2 Joint Undertaking [EUbOPEN grant 875510], Janssen, Merck KGaA, Pfizer, and Takeda.

We thank the CeMM Biomedical Sequencing Facility for NGS sample processing, sequencing, and data curation. We thank the Proteomics team of the Molecular Discovery Platform at CeMM for mass-spectrometric data acquisition. We thank Michael Gütschow from the University of Bonn for support.

## Author contributions

D.S., L.K., Ch.S., and G.E.W. conceived and planned this project. D.S. and L.K. designed and conducted experiments. D.S., L.K., and G.E.W. analysed and interpreted original data. All Chemistry was performed by T.M. and M.L.N. under the supervision of Ch.S. L.K. and S.L. performed and analysed chemoproteomics pulldowns, with data processing being done by S.L. *In vitro* TR-FRET assays were performed by T.M.G. under supervision of R.P.N. *In vitro* thermal shift assays were performed by D.M. under supervision of S.K. and A.C.J. D.S performed and analysed viability-based CRISPR/Cas9 screens, with data processing being done by Ca.S. L.K and D.S. performed and analysed RNA-sequencing and CUT&RUN transcriptional profiling, with data processing being done by L.K. D.S., L.K., T.M., T.M.G., S.L., D.M., D-I.B., M.L.N. established critical reagents and methodology. D.S., L.K., and G.E.W. co-wrote the manuscript with input from all co-authors.

## Data availability

Processed dose-response chemoproteomics data for the SMARCA2/4 ligand (Figure 1b, Extended Data Figure 1a) are provided in Supplementary Table 1. The LC-MS/MS chemoproteomics data, including the used protein reference database, has been deposited in the MassIVE proteomics database with the dataset identifier MSV000100926. Raw read counts, sgRNAs, and sgRNA sequences for the viability-based CRISPR/Cas9 screen (Figure 3a) are provided in Supplementary Table 2. Processed RNA-sequencing tables with gene counts, a list of BCL6 target genes used for gene set enrichment analysis and LISA analysis results (Figure 4, Extended Data Figure 4) are provided in Supplementary Table 3. Differential binding (delta values) for CUT&RUN samples (Figure 5) is provided in Supplementary Table 4. Sequencing data for RNAseq (3’ RNA fingerprinting) and CUT&RUN have been deposited to the Gene Expression Omnibus (GEO). All biological materials are available upon reasonable requests under material transfer agreements (MTA) with the AITHYRA Research Institute for Biomedical Artificial Intelligence of the Austrian Academy of Sciences.

## Methods

### Chemical synthesis

A detailed description of compound synthesis is provided in the supplementary Information file.

### Cell culture

KARPAS-422 and SU-DHL4 iCas9 cells were a gift from Johannes Zuber (Research Institute of Molecular Pathology, Vienna). HEK293T and Lenti-X 293T lentiviral packaging cells (Clontech) were maintained in high glucose Dulbecco’s Modified Eagle’s Medium (DMEM, Gibco/Sigma-Aldrich). KARPAS-422, SU-DHL4, MV4-11, K562, JURKAT, and EOL1 cells were maintained in Roswell Park Memorial Institute (RPMI)-1640 Medium (Sigma-Aldrich). Both DMEM and RPMI were supplemented with 10% Fetal Bovine Serum (FBS) and 1% (v/v) penicillin/streptomycin (all supplied by Thermo Fisher Scientific/Sigma-Aldrich). Cell lines were grown in a humidified incubator at 37 °C and 5% CO_2_, routinely tested for mycoplasma contamination, and authenticated by short tandem repeat profiling.

### Plasmids and oligonucleotides

Generation of the human genome-wide sgRNA library used for the CRISPR/Cas9 screen and viral vectors used for the engineering of inducible Cas9 cell lines have been previously described^79^. The genome-wide sgRNA library used for the viability-based CRISPR–Cas9 screen is shown in Supplementary Table 2. For competitive growth assays, sgRNA for AAVS1 was Golden Gate cloned into a lentiviral sgRNA expression cassette with BFP (hU6-promoter-sgRNA-TracrRNA-mousePGK-eBFP2), and sgRNAs for BAF subunits were Golden Gate cloned into a lentiviral sgRNA expression cassette with mCherry-Neomycin (hU6-promoter-sgRNA-TracrRNA-mousePGK-mCherry-P2A-Neomycin). Guide sequences used for the competitive growth assays are listed below, in the method section pertaining to the competitive growth assays. To generate the BCL6 transcriptional reporter, a fragment containing 9xBCL6 sites upstream from an EF1α promoter controlling the expression of mCherry-T2A-NanoLuc (9xBCL6-EF1α-mCherry-T2A-NanoLuc) was restriction digest cloned into the pLEX305 lentiviral cassette using PspXI and MluI restriction enzymes.

For splitHalo proximity induction assays^52^, the SMARCA2-Hpep3 vector was generated by subcloning the bromodomain (residues 1360-1534) of SMARCA2 from SMARCA2 pDONR223 (a gift from Mikko Taipale) and inserting it into a pRRL lentiviral vector, fused to the Hpep3 tag at the N-terminus with a GGGS linker, under the control of an SFFV promoter. Downstream expression of mTagBFP was driven by the EF1a promoter to confirm transduction e8iciency and allow for gating of SMARCA2-Hpep3positive cells. The BCL6-cpHalo vector was generated by inserting the BCL6 CDS from BCL6 pDONR223 (a gift from Mikko Taipale) into a pRRL lentiviral vector, fused to monomeric eGFP-GSGGSG-cpHalo at the N-terminus separated by a 9xGSG linker under a SFFV promoter. EGFP expression was used to assess transduction efficiency and allow for gating of BCL6-cpHalopositive cells.

### Lentivirus production and transduction

Semiconfluent Lenti-X cells in 10 cm dishes were co-transfected with 1 µg of the envelope plasmid pMD2.G (Addgene #12259), 2 µg of packaging plasmid psPAX2 (Addgene #12260), and 4 µg of the lentiviral plasmid using polyethylenimine (PEI MAX MW 40,000, Polysciences). 3 days after transfection, supernatant containing virus was collected and filtered through a 0.45 mm filter. Target cells were infected in the presence of 8 μg mL^−1^ polybrene (szabo scandic, SACSC-134220).

### Chemoproteomic pulldown

SMARCA bromodomain ligand (1 µmol) with linker was incubated with DMSO-washed NHS-activated sepharose beads (1 mL; ∼20 µmol NHS groups per mL beads) and triethylamine (20 µL) in DMSO (2 mL) on an end-over-end shaker overnight at room temperature in the dark. Aminoethanol (50 µL) was then added to quench remaining NHS-activated carboxylic acid groups. After 16 hours, the beads were washed with 10 mL DMSO followed by 30 mL ethanol to yield the SMARCA affinity matrix for pulldown experiments.

MV4-11 cells were grown in RPMI-1640 medium supplemented with 10% FBS and 1% PenStrep. Cells were harvested, washed with 1xPBS, and then lysed in buffer containing 0.8% Igepal, 50 mM Tris-HCl pH 7.5, 5% glycerol, 1.5 mM MgCl₂, 150 mM NaCl, 1 mM Na₃VO₄, 25 mM NaF and 1 mM dithiothreitol (DTT), supplemented with protease inhibitors. After one freeze-thaw cycle lysates were cleared by centrifugation at 21,000g for 30 min at 4 °C. Protein concentration was determined by BCA assay and adjusted to 5 mg mL⁻¹ and 0.4% Igepal by dilution with detergent-free lysis buffer and buffer containing 0.4% Igepal, respectively.

Aliquots of lysate (0.5 mL) were pre-incubated with eight concentrations of free SMARCA ligand (10, 30, 100, 300, 1,000, 3,000, 10,000 and 30,000 nM) or DMSO vehicle control for 45 min at 4 °C with gentle shaking. Samples were then transferred to 17 µL of affinity matrix equilibrated in lysis buffer in a filter plate (Porvair Sciences, #240002) and incubated for 30 min at 4 °C with gentle shaking. Beads were washed once with 1 mL lysis buffer containing 0.4% Igepal and twice with 2 mL lysis buffer containing 0.2% Igepal. Filter plates were centrifuged at 300 g for 2 min to remove residual detergent-containing buffer and washed three additional times with detergent-free lysis buffer. Pulled down proteins were denatured on-bead in 40 µL 8 M urea in Tris-HCl containing 10 mM dithiothreitol (DTT) for 30 min at 37 °C with shaking at 700 rpm. Iodoacetamide (4 µL of 550 mM) was added for alkylation at 37 °C with shaking at 700 rpm. Samples were diluted to 1 M urea with 250 µL 40 mM Tris-HCl and digested overnight at 37 °C with trypsin (final concentration 1 ng mL⁻¹). Peptides were recovered, acidified with 10 µL 10% formic acid, desalted using C18 StageTips, vacuum-dried and stored at −20 °C until LC–MS/MS analysis.

For proteomic data acquisition, a nanoflow LC−ESI-MS/MS setup comprised of a Dionex Ultimate 3000 RSLCnano system coupled to a Fusion Lumos mass spectrometer (both ThermoFisher Scientific Inc.) was used in positive ionization mode. MS data acquisition was performed in data-dependent acquisition (DDA) mode. For proteome analyses, half of the competition pulldown peptides were delivered to a trap column (Acclaim™ PepMap™ 100 C18, 3 μm, 5 × 0.3 mm, Thermo Fisher Scientific) at a flowrate of 5 μL/min in HPLC grade water with 0.1% (v/v) TFA. After 10 min of loading, peptides were transferred to an analytical column (ReproSil Pur C18-AQ, 3 μm, Dr. Maisch, 500 mm × 75 μm, self-packed) and separated using a stepped gradient from minute 11 at 4% solvent B (0.4% (v/v) FA in 90% ACN) to minute 61 at 24% solvent B and minute 81 at 36% solvent B at 300 nL/min flow rate. The nano-LC solvent A was 0.4% (v/v) FA HPLC-grade water. MS1 spectra were recorded at a resolution of 60,000 using an automatic gain control target value of 4 × 105 and a maximum injection time of 50 ms. The cycle time was set to 2 seconds. Only precursors with charge state 2 to 6 which fall in a mass range between 360 to 1300 Da were selected and dynamic exclusion of 30 s was enabled. Peptide fragmentation was performed using higher energy collision dissociation (HCD) and a normalized collision energy of 30%. The precursor isolation window width was set to 1.3 m/z. MS2 spectra were acquired at a resolution of 30,000 with an automatic gain control target value of 5 × 104 and a maximum injection time of 54 ms.

Protein identification and quantification was performed using MaxQuant (v2.4.9.0)^80^ by searching the LC–MS/MS data against all canonical protein sequences as annotated in the Uniprot reference database (uniprotkb_reviewed_true_canon_human, 20,434 entries, downloaded 29 April 2024) using the embedded search engine Andromeda. Carbamidomethylated cysteine was set as fixed modification and oxidation of methionine and amino-terminal protein acetylation as variable modifications. Trypsin/P was specified as the proteolytic enzyme, and up to two missed cleavage sites were allowed. The minimum length of amino acids was set to seven, and all data were adjusted to 1% peptide spectrum matches and 1% protein false discovery rate. Label-free quantification and match between runs was enabled^80^.

The CurveCurator^81^ pipeline was employed to calculate the dose-dependent residual binding of each protein to the affinity matrix relative to the DMSO control and to estimate effect potency (EC_50_), effect size, and the statistical significance of the observed dose response. MaxQuant LFQ protein intensity values were used for relative protein abundance calculations. Protein groups with missing values and CurveCurator imputations were filtered out. Data processed through the CurveCurator pipeline is shown in Supplementary Table 1.

### CellTiter-Glo® cell viability assay

KARPAS-422 and SU-DHL4 iCas9 cells were seeded at a density of 20,000 – 30,000 cells per well and MV4-11, K562, EOL1, and JURKAT cells were seeded at a density of 10,000 – 15,000 cells per well in opaque white 96-well plates. Cells in 96-well plates were treated with compounds indicated in figures immediately after seeding. 3 days (72 hours) after treatment, the levels of ATP present in each well were measured by mixing cells with CellTiter-Glo® reagent (Promega, G7570) according to the manufacturer’s instructions and reading luminescence levels using a Perkin Elmer Victor X3 (PerkinElmer 2030 software v4.0) or PromegaGloMax Discover (software v4.2.0, firmware v4.92.0) microplate reader . Treated wells were normalized to a DMSO-only control and analysed/plotted using GraphPad Prism (v10.6.1) via fitting of non-linear regression curves for extraction of EC_50_ values.

### BCL6 transcriptional reporter

KARPAS-422 iCas9 cells were transduced with a lentiviral cassette containing a BCL6 transcriptional reporter (9xBCL6-EF1α-mCherry-T2A-NanoLuc) and blasticidin resistance marker. Following several rounds of Blasticidin selection, cells were single cell sorted to generate monoclonal reporter clones. Clones were tested for transcriptional de-repression using the BRD4 recruiting TCIP1^38^. One selected clone was further transduced with a lentiviral cassette for expressing Firefly luciferase and cells were selected using hygromycin.

For reporter assays, transcriptional reporter cells were seeded at a density of 20,000 – 30,000 cells per well in opaque white 96-well plates and immediately treated with compounds indicated in figures. 24 hours after compound treatment the levels of firefly and NanoLuc luciferase in each well was measured using the Nano-Glo® Dual-Luciferase® Reporter Assay System (Promega, N1610) according to the manufacturer’s instructions and a Perkin Elmer Victor X3 (PerkinElmer 2030 software v4.0) or Promega GloMax Discover (software v4.2.0, firmware v4.92.0) microplate reader. To obtain normalized values for transcriptional de-repression, NanoLuc luminescence levels in each well, corresponding directly to the BCL6 reporter, were divided by firefly luminescence levels, corresponding to a non-specific luminescence control. The resulting values were subsequently normalized to a DMSO-only control and plotted using GraphPad Prism (v10.6.1).

### Caspase-Glo® 3/7 apoptosis detection

KARPAS-422 iCas9 cells were seeded at a density of 20,000 – 30,000 cells per well in opaque white 96-well plates and immediately treated with compounds indicated in figures. 16 hours after compound treatment, caspase-3/7 activity in each well was measured by mixing cells with Caspase-Glo® 3/7 Assay System reagent (Promega, G8091) according to the manufacturer’s instructions and reading luminescence levels using a Perkin Elmer Victor X3 (PerkinElmer 2030 software v4.0) or Promega GloMax Discover (software v4.2.0, firmware v4.92.0) microplate reader. Treated wells were normalized to a DMSO-only control and analysed/plotted using GraphPad Prism (v10.6.1)

### HiBiT degradation

HEK293T cells expressing endogenously tagged SMARCA2- or SMARCA4-HiBiT were seeded at a density of 30,000 cells per well in clear-bottom white 96-well plates. The following day, cells were treated with compounds indicated in figures. After 8 or 24 hours treatment, the levels of SMARCA2 or SMARCA4 in each well were measured by mixing cells with LgBit and Nano-Glo® HiBiT Lytic Detection System reagent (Promega, N3030) according to the manufacturer’s instructions and reading luminescence levels using a Perkin Elmer Victor X3 (PerkinElmer 2030 software v4.0) or Promega GloMax Discover (software v4.2.0, firmware v4.92.0) microplate reader. Treated wells were normalized to a DMSO-only control and analysed/plotted using GraphPad Prism (v10.6.1).

### Protein expression and purification for differential scanning fluorimetry

Human bromodomains were expressed and purified following published protocols^45,82,83^. Briefly, proteins were expressed at 18 °C in E. coli BL21(DE3)-R3-pRARE2 cells using pNIC28-Bsa4-based plasmids encoding N-terminally His6-tagged bromodomains. The following domain boundaries were used, with the UniProt IDs given in parentheses: BRD4 BD1, N44-E168 (O60885); SMARCA4, L1451-D1569 (P51532); and PBRM1 BD5, S645- D766 (Q86U86). After protein expression, cells were harvested by centrifugation and re-suspended in lysis buffer (25-50 mM HEPES pH 7.5 at 4 °C, 500 mM NaCl, 10-30 mM imidazole, 0.5-1 mM TCEP, 5% v/v glycerol). Cells were then lysed on ice by sonication, after which polyethylenimine was added to a final concentration of 0.15%, and the lysate was clarified by centrifugation. Immobilized metal affinity chromatography (IMAC) purification of the supernatant was performed by binding to Ni^2+^ beads, followed by elution of the His-tagged protein using an imidazole gradient (50-300 mM imidazole in lysis buffer). Tobacco etch virus (TEV) protease was added to the His-tagged protein, which was then dialysed overnight against dialysis buffer (150 mM NaCl, 20 mM HEPES, pH 7.5 at 4 °C, 0.5 mM TCEP, 5% v/v glycerol). The cleaved protein was separated from the protease and the affinity tag by a second Ni^2+^-rebinding IMAC. As a final purification step, size exclusion chromatography (SEC) was performed using buffer containing 10-20 mM HEPES, pH 7.5 at 4 °C, 150-500 mM NaCl, 0.5 mM TCEP, and 5% v/v glycerol, on a Superdex 75 pg 16/600 HighLoad gel filtration column (GE/Amersham Biosciences) operated on an ÄktaPrime Plus system (GE/Amersham Biosciences). Purified proteins were concentrated, flash frozen in liquid nitrogen, and stored at -80 °C.

### Differential scanning fluorimetry (DSF)

Melting temperatures, *T_m_* values, of bromodomains were determined by DSF using an Agilent MX3005P real-time qPCR instrument (excitation/emission filters = 492/610 nm). Measurements were performed in a 96-well plate with an assay buffer consisting of 5 μM protein in 25 mM HEPES, pH 7.5, 150 mM NaCl, 0.5 mM TCEP, 2.5% (v/v) DMSO, and varying concentrations (25, 100, and 250 μM) of the SMARCA ligand. The fluorescent dye SYPRO Orange (5000×, Invitrogen) was added at a dilution of 1:1000. The fluorescence signal was monitored upon temperature increase from 25 to 95°C, at a heating rate of 3 °C/min, and *T_m_* values were calculated after fitting the fluorescence curves to the Boltzmann function.

### Western blotting

KARPAS-422 and SU-DHL4 iCas9 cells were seeded at a density of 0.5-1 x 10^6^ cells per mL in 12-well plates and immediately treated with compounds indicated in the corresponding figures. After specified time-points indicated in figure legends, cells were harvest into 1.5 mL Eppendorf tubes and washed once with PBS before lysis on ice for 20 minutes with RIPA buffer (150 mM NaCl, 1% Triton X-100, 0.5% sodium deoxycholate, 0.1% SDS, 50 mM Tris-HCl pH 8) supplemented with benzonase (1:1000, Sigma-Aldrich, 70746-3), HALT EDTA-free protease inhibitor cocktail (1:100, Thermo Fisher Scientific, 78437), and 20 mM DTT. Lysates were prepared with 4 x Bolt LDS sample bu8er (Thermo Fisher, B0008), heated at 95 °C for 5 minutes, and then run on SurePAGE, 4–12% Bis-Tris gels (GenScript) with Tris-MES-SDS running buffer (GenScript). Proteins were transferred to nitrocellulose blotting membranes (Amersham Protran, 0.45 µm NC), blocked for 1 hour with 5% milk or 3% BSA in TBS-T at room temperature, before incubation with primary antibodies at 4 °C, overnight. The following primary antibodies were used: GAPDH (1:1000, Cell Signalling Technology, 2118), SMARCA4 (1:1000, Cell Signalling Technology, 49360), caspase-3 (1:1000, Cell Signalling Technology, 9662), BCL6 (1:1000, Cell Signalling Technology, 5650), vinculin (1:1000, Cell Signalling Technology, 13901), p21 Waf1/Cip1 (1:1000, Cell Signalling Technology, 2947), FOXO3 (1:1000, Cell Signalling Technology, 2497), ARID3A (1:1000, Cell Signalling Technology, 43033). Following overnight treatment, membranes were washed in TBS-T and incubated with horseradish peroxidase (HRP)-conjugated secondary antibodies for 1-2 hours at room temperature. The following secondary antibody was used: HRP anti-rabbit IgG (1:5000, Cell Signaling Technology, 7074). Lastly, membranes were washed again with TBS-T and then imaged on LI-COR ODYSSEY XF operated on Image Studio software (v6.1.0.79).

### Protein expression and purification for TR-FRET

For protein expression and purification, codon-optimized coding sequences for N-terminal Strep-Avi-TEV–tagged human BCL6 (residues 5–360), SMARCA2 (residues 1373–1399,1418–1511), and SMARCA4 (residues 1448–1568) in pET28a(+) expression vectors were obtained from Twist Bioscience and transformed into *E. Coli* Rosetta (BCL6) or Rosetta-pCDF-BirA^84^ cells used for in-vivo biotinylation (SMARCA2, SMARCA4). A single colony was used to inoculate a starter culture in LB medium supplemented with 50 μg/mL kanamycin and 25 μg/mL chloramphenicol. For SMARCA2/4 samples, an additional 50 μg/mL of spectinomycin was included and cultures were incubated overnight at 37 °C. Large scale expression was performed in TB medium supplemented with 50 μg/mL kanamycin and 25 μg/mL chloramphenicol and for SMARCA2/4 samples, an additional 50 μg/mL spectinomycin. Cells were incubated at 37°C, shaking at 140 rpm until an OD_600_ of 1.2 was reached. Protein expression was induced by addition of 400 µM IPTG and the cultures were further incubated at 18°C overnight. In the case of SMARCA2/4 biotin was added at a final concentration of 100 mM for *in cellulo* biotinylation of the Avi-tag during this incubation period^84^. After overnight incubation, the cells were harvested by centrifugation (4,000 x g, 20 min, 4 °C) and the cell pellet was solubilized in buffer (50 mM Hepes-HCl pH 8.0, 200 mM NaCl, 1 mM TCEP) supplemented with 1 mM PMSF and lysed by sonication. Following ultracentrifugation (20000 × g, 40 minutes), the soluble fractions were filtered through 0.45 µm membranes and incubated with Streptactin XT High-capacity beads for 1 h, rolling at 4°C. Following incubation, the beads were washed with wash buffer (50 mM Hepes-HCl pH 8.0, 200 mM NaCl, 1 mM TCEP) and the proteins were eluted with elution buffer (50 mM Hepes-HCl pH 8.0, 200 mM NaCl, 1 mM TCEP, 50 mM biotin). The protein containing fractions were concentrated using Amicon Ultra centrifugal filters (Millipore). For BCL6, the protein was FITC labeled by overnight incubation with a 1.5-fold molar excess of NHS-FITC (ThermoFisher #46409) at 4°C. BCL6-FITC was passed over PD10 columns (Cytiva) and concentrated again. Finally, all proteins were further purified on a Superdex 75 16/600 GL size exclusion column (Cytiva) in buffer containing 50 mM Hepes pH 7.5, 200 mM NaCl, and 1 mM TCEP. The purified proteins were concentrated using an Amicon Ultra centrifugal filter (Millipore), flash-frozen in liquid nitrogen, and stored at -80°C.

### Time-resolved fluorescence resonance energy transfer (TR-FRET)

For TR-FRET ternary complex formation assays, the assay mix of 100 nM SMARCA2-Biotin or 100 nM SMARCA4-Biotin, 500 nM BCL6-FITC, 2 nM Streptavidin-Tb (Invitrogen #PV3965) prepared in assay buffer (50 mM Hepes pH 7.5, 200 mM NaCl 0.1% Pluronic F-68 solution (Sigma), 1 % BSA, 1 mM TCEP) was added at 12.5 μL/well to a black 384 well plate (Corning # 4514). For dose-response measurements, compounds were dispensed using a D300 Dispenser (Tecan), the DMSO content was normalized to 0.5% (v/v), and the plate was sealed with aluminum foil. After 15 minutes of incubation at room temperature, the TR-FRET signals at 490 nm (terbium) and 520 nm (FITC) were recorded with a 60 μs delay over 400 μs and over 5 cycles for each data point using a BMG PHERAstar FSX microplate reader (PHERAstar Software v5.70 R6, Firmware v1.33). The TR-FRET signals were extracted by calculating the 520/490 nm ratio and averaged over the 5 cycles for each data point. Data from two independent replicates were plotted using GraphPad Prism (v10.6.1) and curve fitting was performed using a four-parameter variable slope equation.

### SplitHalo proximity induction

The splitHalo interaction assay was designed based on previous reports describing a split HaloTag enzyme to record protein-protein interactions^52^. The splitHalo BCL6-SMARCA ternary complex formation reporter cell line was generated by transducing two split constructs for SMARCA2-Hpep3 and BCL6-cpHalo, respectively. Lentivirus particles were produced for both constructs and HEK293T cells were transduced consecutively with BCL6-cpHalo first and SMARCA2-Hpep3 second.

50,000 HEK293T BCL6-SMARCA2 splitHalo reporter cells were seeded into transparent 96-well plates the night before for overnight attachment. For competition experiments, cells were pre-treated with the indicated ligand concentration for 30 minutes. For treatment with test compounds, drug dilutions were prepared in DMEM with a final concentration of 100 nM TAMRA dye (MedChemExpress, HY-D2270). After 3 hours of simultaneous compound incubation and HaloTag labelling, supernatant was removed and cells were washed with 1xPBS once to remove excess TAMRA dye and phenolred. Cells were trypsinized, resuspended in cold FACS buffer (1x PBS, 5% FBS, 1 mM EDTA), transferred to transparent U-bottom 96-well plates and kept on ice until measurement. Cell fluorescence was measured on either the Attune™ CytPix™ (Attune Cytometric software v6.0.1) or the BD LSRFortessa™ X-20 (BD FACSDiva software v9.0.1) flow cytometer in the high-throughput plate mode. For data analysis, TAMRA background subtraction was performed based on the signal obtained from GFP^-^BFP^-^ (double-negative/untransduced) cells and TAMRA signal was normalized to background-subtracted GFP signal to account for varying transduction efficiency. The resulting raw TAMRA signal was then normalized to the average of DMSO TAMRA signal to assess ternary complex formation relative to DMSO vehicle-treated cells. Data were plotted using GraphPad Prism (v10.6.1) and curve fitting performed using a four-parameter variable slope equation. Gating strategy is shown in Supplementary Figure 1a.

### Viability-based CRISPR/Cas9 screen

Doxycycline-inducible Cas9 (iCas9) KARPAS-422 cells were transduced in the presence of 8μg mL^-1^ polybrene with a genome-wide sgRNA library targeting 18184 human genes (6 sgRNAs per gene, 1000 non-targeting controls,109082 guides total) at a multiplicity of infection of 0.07 and approximately 1000x library representation (Supplementary Table 2). Cells expressing sgRNAs were selected for 17 days with G418 (1 mg mL^−1^, Sigma-Aldrich, A1720). Following selection, Cas9 expression was induced with doxycycline (0.25 μg mL^−1^, PanReac AppliChem, A2951). Four days after Cas9 induction, cells were treated with DMSO or TRIP1 (400 nM), in two biological replicates each. Every 3-4 days cells were split with fresh media containing DMSO or TRIP1 (400 nM). After 33 days of compound treatment, surviving cells were isolated via lymphocyte separation medium gradient centrifugation and harvested.

For library preparation, genomic DNA from surviving cells was isolated by cell lysis (10 mM Tris-HCl, 150 mM NaCl, 10 mM EDTA, 0.1% SDS), proteinase K treatment (New England Biolabs, P8107) and DNAse-free RNAse digestion (Thermo Fisher Scientific). DNA was then purified through two rounds of phenol extraction (Sigma Aldrich, P4557) and isopropanol precipitation (Sigma-Aldrich, I9516). Isolated genomic DNA was subjected to a two-step PCR amplification of the sgRNA cassette using AmpliTaq Gold polymerase (Thermo Fisher Scientific, 4311818), with sample-specific barcodes introduced during the first PCR and standard Illumina adapters added in the second PCR. The resulting PCR products were purified using Mag-Bind TotalPure NGS beads (Omega Bio-tek, M1378-00) and the final Illumina libraries were pooled and sequenced on an Illumina NovaSeq 6000 platform.

Downstream processing of the screen data utilized the crispr-process-nf Nextflow workflow (https://github.com/ZuberLab/crispr-process-nf/). In brief, adapter, barcode, and spacer sequences were removed from raw FASTQ reads using cutadapt (v4.4), followed by demultiplexing according to sample-specific barcodes. Processed reads were aligned to the genome-wide sgRNA library^79^ with Bowtie2 (v2.4.5), and sgRNA counts were obtained using featureCounts (v2.0.1). Resulting count matrices were further analyzed with the crispr-mageck-nf pipeline (https://github.com/ZuberLab/crispr-mageck-nf/) for statistical evaluation. Gene-level enrichment scores were computed using MAGeCK (v0.5.9)^85^ by comparing TRIP1-treated samples to DMSO controls, applying median normalization and estimating variance across biological replicates. The large language model (LLM) Claude Opus (versions 4.1/4.6, Anthropic) was used for AI (artificial intelligence)-aided writing of code for data visualization. All AI-generated code was manually proofread before usage.

### CRISPR knockout competitive growth assay

KARPAS-422 and SU-DHL4 iCas9 cells were transduced with lentiviral sgRNA expression cassettes containing either AAVS1 targeting sgRNA and BFP or BAF targeting sgRNA and mCherry-P2A-Neomycin. Cells containing sgAAVS1 were selected by sorting for BFP positive cells. Cells containing BAF targeting sgRNAs were selected via neomycin and then confirmed via flow cytometry for mCherry expression. Following selection, 15,000 sgAAVS1 containing cells were mixed with 15,000 sgBAF containing cells in clear 96-well plates and then induced with 0.25 ug/mL doxycycline. 4 days after induction, the BFP and mCherry ratios for each sgRNA mixture in each iCas9 cell line was measured using flow cytometry for timepoint 0 and then cells were treated with DMSO, TRIP1 (400 nM), BRD4 targeting TCIP1 (4 nM), or p300/CBP targeting TCIP3 (4 nM). SU-DHL4 iCas9 sgRNA cell mixtures were only treated with DMSO or TRIP1 (400 nM). BFP and mCherry ratios for each sgRNA mixture were measured every 3-4 days using the BD LSRFortessa™ X-20(BD FACSDiva software v9.0.1) flow cytometer in the high-throughput plate mode and compounds were replenished.

For data normalization, first the percentage of mCherry+ (sgBAF) cells was calculated by subtracting the percentage of BFP+ (sgAAVS1) cells from 100. A logit transformation approach was used to account for the variation in initial mixing ratios across replicates and conditions. The percentage of sgBAF containing cells was converted to a proportion and the logit function [log (p/ (1 − p))] was applied to transform the proportion to a log-odds scale. The logit function was applied to all timepoints, with timepoint 0 serving as the baseline reference for each replicate, gene knockout, and treatment. To center all conditions to 50% at timepoint 0, the logit value at timepoint 0 was subtracted from the logit of each timepoint’s proportion, and the logit of 0.5 was added. The inverse logit function [exp(x) / (1 + exp(x)) × 100] was then used to convert all normalized logit values to a percentage scale. Certain knockout and treatment conditions, at days 18 and 22, involving competitive growth assays in KARPAS-422 iCas9 cells were excluded from the analysis due to a lack of detected live cells. Gating strategy is shown in Supplementary Figure 1b. The large language model (LLM) Claude Opus (versions 4.1/4.6, Anthropic) was used for AI (artificial intelligence)-aided writing of code for data normalization and visualization. All AI-generated code was manually proofread before usage.

The following sgRNAs were used for competitive growth assays: sgBRD7 (GGCAAGTCTAATCTCACAG), sgARID1B (GTCTGCGTCAAAGAGATCGG), sgPBRM1 (GAGTTGTCGGAATAACCAA), sgPHF10 (GGTTATCCAGGTACCTCAA), sgARID2 (GTGGTAGGAGTAAAACGGA), sgSMARCA4 (GCAGCAGACAGACGAGTACG), sgAAVS1 (GCTGTGCCCCGATGCACAC).

### CUT&RUN

The CUT&RUN experiment was conducted following the CUT&RUN protocol described by EpiCyper (EpiCypher CUTANA™ CUT&RUN Protocol v2.0 2022) with minor modifications. Briefly, 10 million KARPAS422 cells per treatment were seeded into T25 flasks and treated with the indicated compounds for 30 minutes, 1 hour and 2 hours (1 µM, run 1) or 4 and 8 hours (2 µM, run 2). Meanwhile, Concanavalin A beads were processed in bulk as described, and activated beads were stored at 4°C until use. At the end of the treatment incubation time, cells were split into two independent replicates and batch-processed with 10% excess. After cells were washed with PBS, the cell pellet was resuspended in nuclear extraction buffer and incubated on ice for 10 minutes. Samples were centrifuged again, and the nuclei pellet was resuspended in nuclear extraction buffer. Nuclei were mixed with activated beads and allowed to adsorb at RT for 10 minutes. In the meantime, antibody solutions were prepared by mixing 50 µL antibody buffer with 1 µL antibody per reaction. The following antibodies were used: IgG rabbit isotype control (Cell Signaling Technology, CST3900), BCL6 (Cell Signaling Technology, CST49360), SMARCA4 (Cell Signaling Technology, CST49360), H3K27ac (Cell Signaling Technology, CST8173), RNA Pol II Ser2 phospho (Cell Signaling Technology, CST13499), RNA Pol II Ser5 phospho (Cell Signaling Technology, CST13523). After 10 minutes, nuclei-bead mixture was aliquoted into PCR tubes and placed on a magnet. Following separation, supernatant was discarded, antibody solution was added, and samples were gently vortexed to resuspend. Tubes were put to nutate overnight at 4 °C.

On the next day, nuclei-beads were washed twice with digitonin buffer before adding 50 µL pAG-MNase at a final conc. 700 ng/mL in digitonin buffer. Nuclei were incubated with pAG-MNase at RT, while a 2 mM CaCl_2_ digitonin buffer was prepared to enable the catalytic reaction. After 30 minutes, nuclei were washed with digitonin buffer twice before adding 50 µL CaCl_2_ solution and incubating for 90 minutes at 4°C on a nutator. Stop buffer was freshly prepared to inhibit the MNase activity by chelating Ca^2+^ ions and 33 µL were added per sample. The reaction was incubated at 37°C for 10 minutes to release cleaved chromatin into the supernatant. Supernatant was transferred into fresh tubes and DNA was purified using the Monarch® Spin PCR & DNA Cleanup Kit (New England Biolabs, T1130L) according to the manufacturer’s instructions. DNA was eluted in 12 µL elution buffer pre-heated to 64°C. DNA concentration was quantified using the Qubit^TM^ fluorometer (Invitrogen) using the Qubit™ dsDNA HS Assay Kit (Invitrogen, Q32851). DNA was stored at -20°C and processed the next day.

Sample library was prepared as previously described with modifications^86^. Briefly, from 5 ng input DNA or all available DNA for the IgG isotype control, if necessary, using the NEBNext® Ultra™ II DNA Library Prep Kit for Illumina® (New England Biolabs, E7645L) and multiplexing samples using dual indexing primers NEBNext® Multiplex Oligos for Illumina® (New England Biolabs, E7600S). DNA was diluted with water and incubated with end prep mastermix. Diluted adapters were added to all samples and ligated to DNA fragments. Next, USER enzyme was added and incubated at 37°C for 15 minutes. Meanwhile, Mag-Bind® TotalPure NGS magnetic beads (Omega Bio-tek, M1378-02) were warmed up to RT. 80 µL beads were added and pipetted to mix. After incubation, the samples were separated on a magnetic rack, supernatant was removed and beads were washed with 80% EtOH twice. After complete evaporation, DNA was eluted using 15 µL 0.1x TE buffer (IDTE (1X TE Solution), IDT, 11-05-01-05) and stored at -20°C overnight. The day after, combinatorial dual indexing primers and 2x Q5 master mix were added for PCR. PCR settings were denaturation at 98°C for 30 sec, 14 annealing/elongation cycles of 98°C 10 sec, 57°C 10 sec, followed by a final elongation step at 65°C for 5 minutes. Mag-Bind® TotalPure NGS magnetic beads were warmed up to RT and added to DNA at a 1:1 ratio. DNA was washed twice with 80% EtOH. Once dry, DNA was eluted using 15 µL 0.1x TE buffer. After magnetic separation, DNA-containing supernatant was transferred to fresh tubes and quantified using the Quibit^TM^ fluorometer (see above). Samples were pooled using 50 ng each and sent to Genewiz/Azenta for fragment analysis with a Bioanalyzer. Pooled samples were sequenced paired-end on a NovaSeq X+ platform with a read length of 150 bp.

### CUT&RUN analysis

Data were de-multiplexed and raw .fastq files were processed using the *Nextflow*^87^ (Nextflow/23.04.1)-based *nf-core/cutandrun*^88^ pipeline (https://nf-co.re/cutandrun/3.0/) with the following settings:

> nextflow run nf-core/cutandrun -r 3.2.2 -profile

> singularity --peakcaller ’macs2, seacr’ --genome “GRCh38”

> --dt_calc_all_matrix true --normalisation_mode “CPM”

to compute CUT&RUN signals of single replicates in bam format with specification of blacklist regions. These packages and indicated versions thereof were used in the following analysis: SAMtools/1.18-GCC-12.3.0, deepTools/3.5.1 and BEDTools/2.30.0. After inspecting quality control data for all samples to confirm reproducibility of single replicates and IgG antibody isotype controls, marked up bam files from the nextflow pipeline were merged using *samtools merge*. Files were sorted by read name using *samtools sort* and mate information was fixed with *samtools fixmate*. Files were then sorted by genomic position using *samtools sort*. Sorted bam files were filtered for annotated ENCODE hg38 blacklist regions^89^ using *bedtools intersect* and indexed using *samtools index*. Quality of blacklist-removed files was controlled using *multiBamSummary*. Blacklist-removed, indexed files were used to (1) generate BigWig coverage tracks using *bamCoverage* with the settings:

> --normalizeUsing CPM \--smoothLength 60 \--binSize 10

and (2) call peaks using MACS2 peakcalling^90^ employing *macs2 callpeak* (MACS2/2.2.7.1) with the settings:

> -f BAM -g hs -n --broad -q 0.05

Overlapping peaks were calculated using *bedtools intersect* and selected peak sets were compared to publicly available datasets from the ChIP atlas (v3.0)^91^.

To calculate antibody-specific signals on specified genome regions, a binding matrix was created using the *computeMatrix* command with the settings:

> --skipZeros and --missingDataAsZero

Heatmaps and profiles were generated from the matrix using *plotHeatmap* and *plotProfile*. For clustered samples, k means clustering was applied by specifying --kmeans. To calculate signal around BCL6 or SMARCA4 peaks, MACS2 broad peaks of the respective DMSO samples were used as input files and regions centered using mode “reference-point” and --referencePoint center -b 5000 -a 5000. To compute signals on single genes, hg38-annotated gene lists were used^92^. Length-scaled signal across gene bodies and unscaled signal 3000 bp up-and downstream was calculated using the mode “scale-regions” and --regionBodyLength 5000 -b 3000 -a 3000.

Correlation metrics were calculated as follows. Coordinates of single genes were extended by 3000 bp up- and downstream to cover proximal regulatory regions. Size-normalized signals were calculated per gene using *multiBigwigSummary* and exported for each condition. Signal difference between TRIP1- and DMSO-treated samples was calculated and used for plotting boxplot and correlation graphs. Genome tracks were visualized as bigWig files in IGV (Integrated Genome Viewer, v2.17.4)^93^. CUT&RUN correlation analysis and data visualization code was created using the large language model (LLM) Claude Opus (versions 4.1/4.6, Anthropic). All AI-generated code was manually proofread before usage.

### RNA-seq (3’ RNA fingerprinting)

500,000 KARPAS-422 cells were seeded in 24-well plates and treated with compounds of interest for the indicated times and concentrations in triplicates. At harvest, cells were transferred to round-bottom 96-well plates and washed with PBS. RNA was extracted from washed cells using the NucleoSpin 96 RNA Core Kit (Macherey-Nagel, 740466.4) according to the manufacturer’s instructions. Eluted RNA was frozen and submitted to the Biomedical Sequencing Facility (BSF) at the Research Center for Molecular Medicine of the Austrian Academy of Sciences. NGS libraries were prepared from total RNA samples with the QuantSeq 3’ mRNA-Seq V2 Library Prep Kit with Unique Dual Indices (UDI) (Cat. No. 191, 193 – 196, Lexogen GmbH, Vienna, Austria) and the UMI Second Strand Synthesis Module for QuantSeq FWD (Illumina, Read 1, Cat. No. 081.96). Resulting library concentrations were quantified with the Qubit® 1X dsDNA HS Assay Kit (Q32856, Thermo Fisher Scientific, Waltham, MA, USA) and the fragment size distribution was assessed using the High Sensitivity DNA Kit (5067-4626, Agilent, Santa Clara, CA, USA) on a 2100 Bioanalyzer High-Resolution Automated Electrophoresis instrument (G2939A/B, Agilent). Before sequencing, sample-specific NGS libraries were diluted and pooled in equimolar amounts. Libraries were single-end sequenced with 100bp fragments on the NovaSeq 6000 platform.

### RNA-seq analysis

Raw sequencing files were supplied as bam files. Bam files were converted into fastq files using *samtools fastq* (SAMtools/1.18-GCC-12.3.0) and quality was controlled using FastQC (FastQC/0.11.9-Java-17). Sequences were trimmed from adapters, NextSeq artefacts and polyA sequences using *cutadapt* (cutadapt/4.4-GCCcore-11.3.0) with parameters:

> -m 20 -O 20 -a “polyA=A{20}” -a “QUALITY=G{20}” -n 2

> ”$file” |

> -m 20 -O 3 -e 0.100000 --nextseq-trim=10 -a

> ”A{18}AGATCGGAAGAGCACACGTCTGAACTCCAGTCAC” |

> -m 20 -O 20 -g “AGATCGGAAGAGCACACGTCTGAACTCCAGTCAC”.

Afterwards, FastQC was performed again to confirm correct trimming. Trimmed reads were aligned and mapped to hg38 using STAR alignment^94^ (STAR-2.7.11b) with the parameters:

> --runThreadN 8 --outSAMtype BAM SortedByCoordinate --

> outFilterType BySJout --outFilterMultimapNmax 20 --

> alignSJoverhangMin 8 --alignSJDBoverhangMin 1 --

> outFilterMismatchNmax 999 --outFilterMismatchNoverLmax

> 0.6 --alignIntronMin 20 --alignIntronMax 1000000 --

> alignMatesGapMax 1000000 --outSAMattributes NH HI NM MD.

MultiQC^95^ (MultiQC/1.14-foss-2022a) was run to confirm integrity of aligned reads. Read count matrices were computed using *htseq-count*^96^ (HTSeq/0.11.2-foss-2020b) with settings

> -m intersection-nonempty -s yes -f bam.

To annotate genome coverage for visualisation in IGV (Integrated Genome Viewer, v2.17.4), signal was calculated using *bamCoverage* (deepTools/3.5.1-foss-2022a) with default settings.

Differential expression analysis was performed using DESeq2^97^ (v 1.42.1), which employs the Wald statistical test for pairwise comparison and Benjamini-Hochberg multiple-testing correction^98^. Log_2_ fold (log2FC) changes were shrunk using the apeglm method^99^. Genes were filtered for low abundance-artefacts to only include genes with at least 10 counts in more than three samples. Significant differential expression was defined as |log2FC| > 1 and p adj < 0.05. Gene set enrichment analysis (GSEA) was performed using a manually curated gene set of BCL6 TCIP targets which were reported for BCL6-targeting TCIPs (**Supplementary Table 3**)^38,42,49^.

Epigenetic Landscape In Silico deletion Analysis (LISA)^59^ analysis was performed using all differentially expressed genes after 16h 0.5 µM TRIP1 treatment compared to DMSO. Ranked transcription factors predicted to regulate the gene set were plotted against their p-value. The large language model (LLM) Claude Opus (versions 4.1/4.6, Anthropic) was used for AI (artificial intelligence)-aided writing of RNA-seq data visualization code. All AI-generated code was manually proofread before usage.

---

**Extended Data Figure 1.**
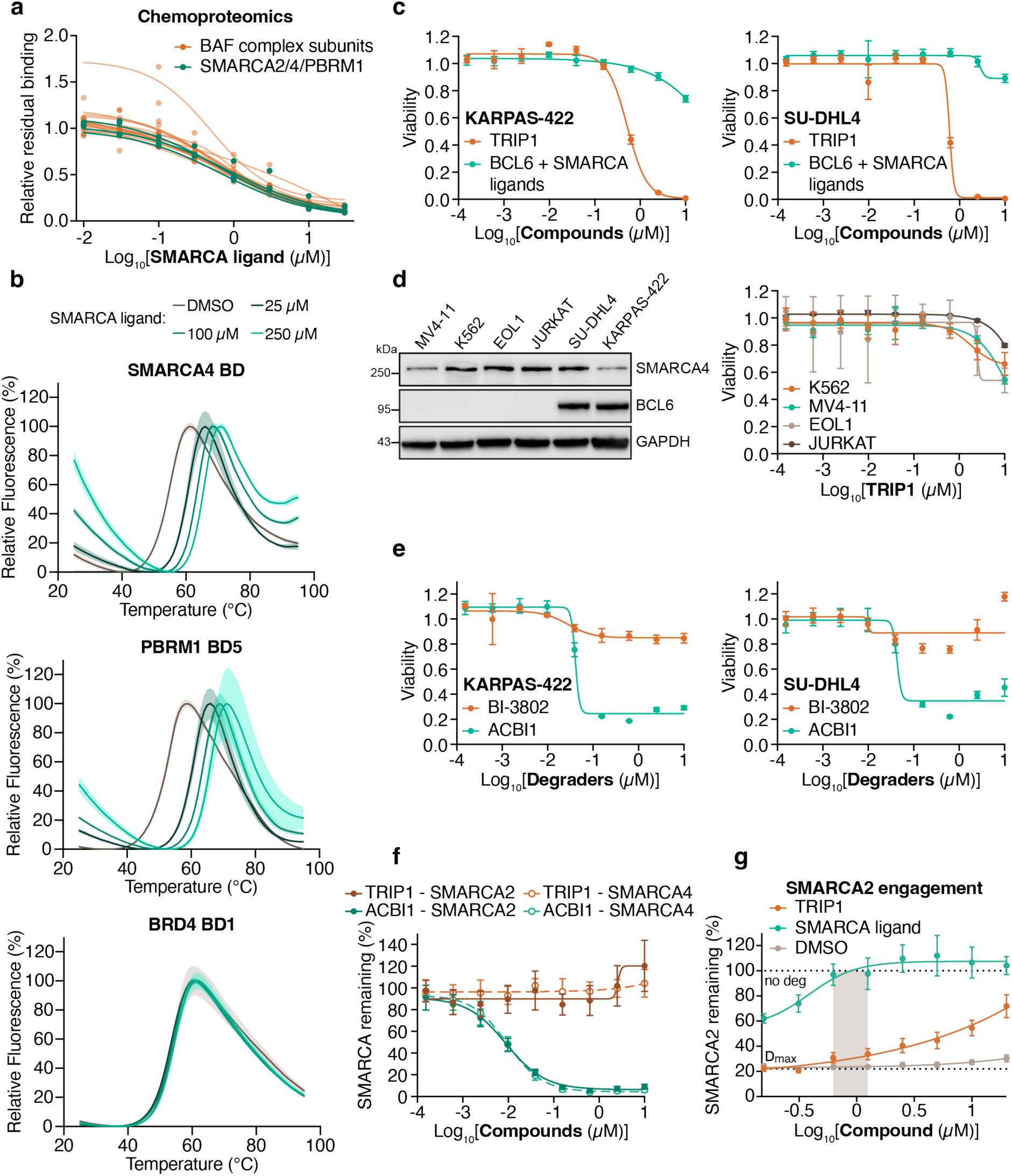
TRIP1 activity differs from degraders and inhibitors and is confined to BCL6-expressing cells. **a**, Dose response curves of BAF complex subunits from the chemoproteomic competition assay highlighted in Figure 1b. **b**, Assessment of ligand binding and selectivity using differential scanning fluorimetry. Thermal melting curves for SMARCA4 BD (top), PBRM1 BD5 (middle), and BRD4 BD1 (bottom) were generated by monitoring relative fluorescence intensity as a function of temperature (25-95°C). Proteins (5 μM) were incubated with DMSO or increasing concentrations (25, 100, and 250 μM) of the SMARCA ligand. Data represent mean ± SD (shaded regions), n = 3 independent replicates. **c**, CellTiter-Glo cell viability assays in DLBCL cell lines. KARPAS-422 (left) and SU-DHL4 (right) cells were treated with a dilution series of TRIP1 or the combination of BCL6 and SMARCA ligands for 72 hours. Viability with compound treatment is normalized to DMSO vehicle control; data represent mean ± SD, n = 3 independent replicates. **d**, Cell viability assays in non-BCL6 expressing cell lines. CellTiter-Glo assays were performed in cell lines not expressing BCL6, as assessed by Western blot (left). K562, MV4-11, JURKAT, and EOL1 cells were treated with a dilution series of TRIP1 for 72 hours (right). Viability is normalized to DMSO vehicle control; data represent mean ± SD, n = 3 independent replicates. **e**, Cell viability assays with degraders. KARPAS-422 (left) and SU-DHL4 (right) cells were treated with a dilution series of BCL6 (BI-3802) or SMARCA degrader (ACBI1) for 72 hours. Viability with compound treatment is normalized to DMSO vehicle control; data represent mean ± SD, n = 3 independent replicates. **f**, SMARCA degradation assay. HEK293T cells expressing endogenously tagged SMARCA2- or SMARCA4-HiBiT were treated with a dilution series of TRIP1 or SMARCA degrader for 24 hours. SMARCA2/4-HiBiT levels are shown as a remaining fraction relative to DMSO vehicle control; data represent mean ± SD, n = 6 independent replicates. **g**, SMARCA2 target engagement. HEK293T cells expressing endogenously tagged SMARCA2-HiBiT were pre-treated with a dilution series of TRIP1 or the SMARCA ligand for 2 hours, followed by co-treatment with 0.1 µM ACBI1 for 8 hours. SMARCA2-HiBiT levels are shown as a remaining fraction relative to DMSO vehicle control (no ACBI1 treatment). D_max_ represents the fraction of SMARCA2 remaining when ACBI1 is co-treated with DMSO. Gray bar indicates early cytotoxic/pro-apoptotic concentrations of TRIP1. Data represent mean ± SD, n = 6independent replicates.

**Extended Data Figure 2.**
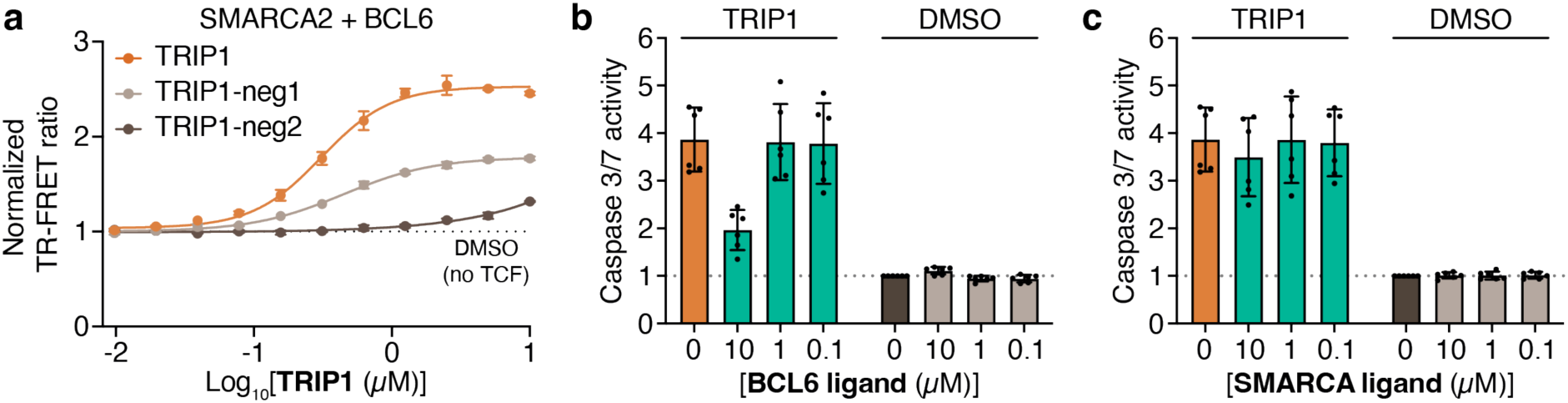
SMARCA2/4 and BCL6 binding are functionally required for TRIP1 efficacy. **a**, *In vitro* TR-FRET ternary complex formation assay. The interaction between biotinylated SMARCA2 bromodomain (100 nM) and FITC-labeled BCL6 BTB domain (500 nM) was measured with increasing concentrations of TRIP1. The biotinylated SMARCA2 bromodomain was captured with streptavidin-terbium cryptate donor (SA-Tb, 2 nM), and TR-FRET signal was monitored upon complex formation. The resulting TR-FRET ratio was normalized to the DMSO vehicle control; data represent mean ± SD, n = 2 independent replicates. **b**, **c**, CaspaseGlo 3/7 apoptosis competition. KARPAS-422 cells were pre-treated with DMSO, the BCL6 ligand (**b**), or SMARCA ligand (**c**) for 8 hours, followed by co-treatment with 1 µM of TRIP1 for 16 hours. Caspase 3/7 activity is normalized to DMSO vehicle control without TRIP1 co-treatment; data represent mean ± SD, n = 6 independent replicates.

**Extended Data Figure 3.**
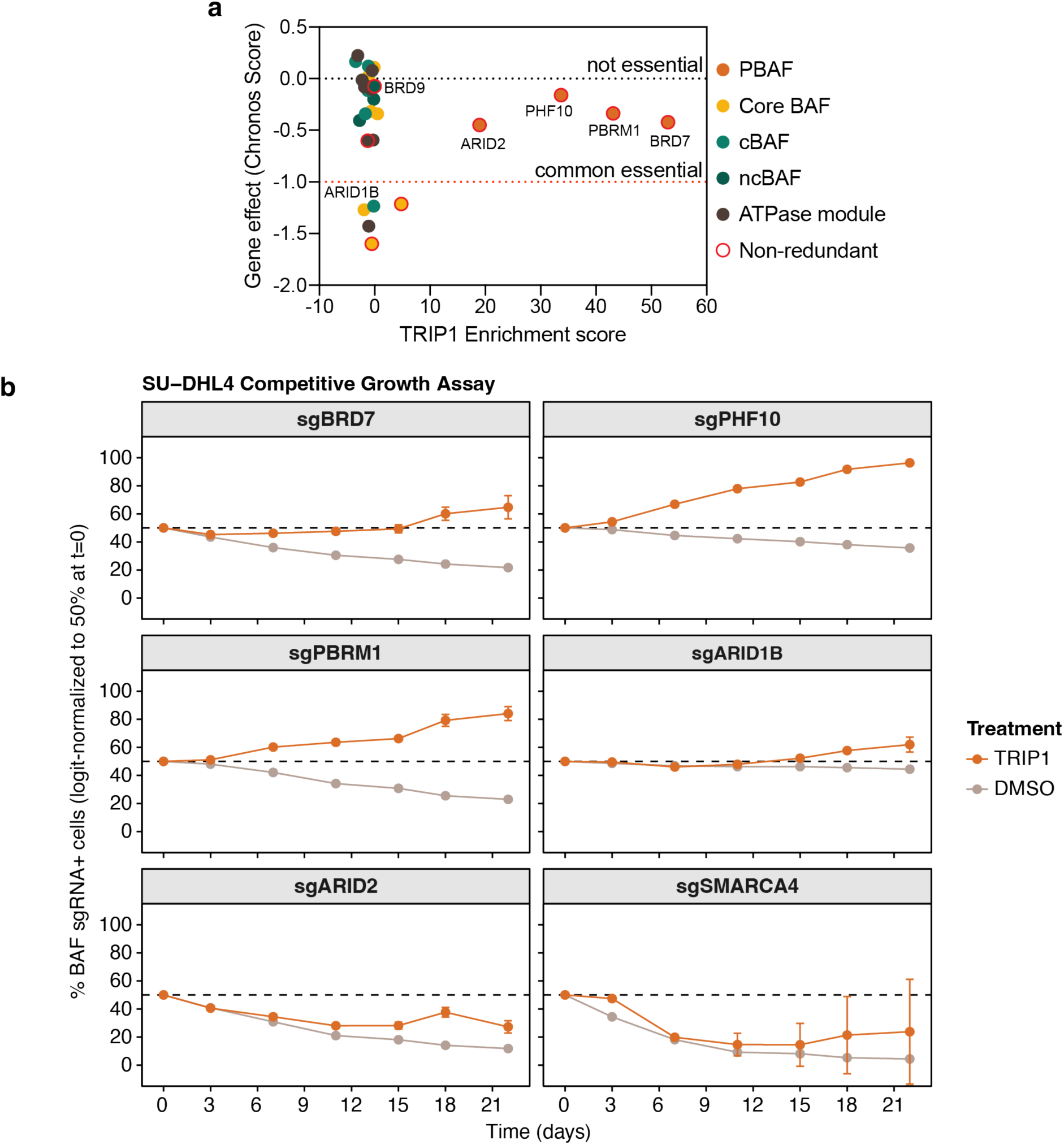
TRIP1 induced dependencies of PBAF components in SU-DHL4 cells. **a**, Essentiality of BAF complex members in KARPAS-422 cells. DepMap CRISPR (Chronos) dependency scores of BAF complex subunits in KARPAS-422 cells plotted against the TRIP1 CRISPR screen enrichment scores. Genes highlighted with a red border indicate non-redundant subunits. **b**, Competitive growth assay in SU-DHL4 cells. Control SU-DHL4 iCas9 cells expressing AAVS1-targeting sgRNA and BFP were mixed with SU-DHL4 iCas9 cells expressing BAF subunit-targeting sgRNAs and mCherry. Cell mixtures were treated with DMSO or TRIP1 (400 nM) and evaluated in regular intervals via flow cytometry. Data were logit-transformed and normalized to 50% at day 0 to enable comparison across conditions. Data represent mean ± SD, n = 3 independent replicates.

**Extended Data Figure 4.**
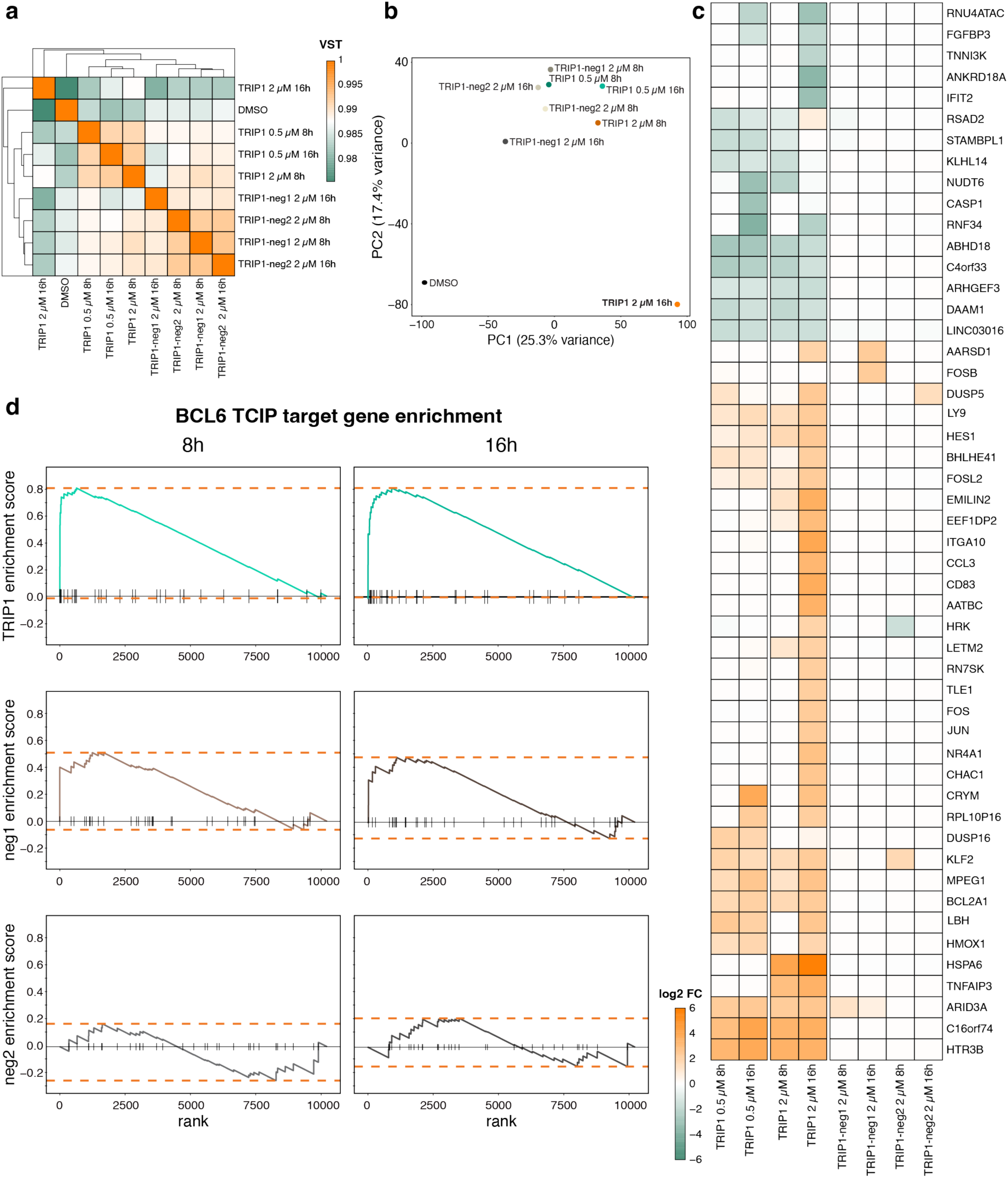
Transcriptional response to TRIP1 treatment. **a**, Gene expression correlation between experimental conditions. Variance stabilizing transformation (VST) was calculated, and conditions were hierarchically clustered. **b**, Principal component analysis (PCA) of experimental conditions. TRIP1 dissimilarity to DMSO and negative controls increases with treatment duration and/or concentration. **c**, Gene expression changes (3’ RNA fingerprinting) in KARPAS-422 cells of the top 50 most variably expressed genes with significant changes (P adj < 0.05) across conditions, concentrations, and timepoints. **d**, Related to Figure 4c. Gene set enrichment analysis (GSEA) of BCL6 target genes and genes shown to be regulated by BCL6 targeting TCIPs following treatment with TRIP1, TRIP1-neg1 and TRIP1-neg2 (2 µM) for 8h and 16h.

**Extended Data Figure 5.**
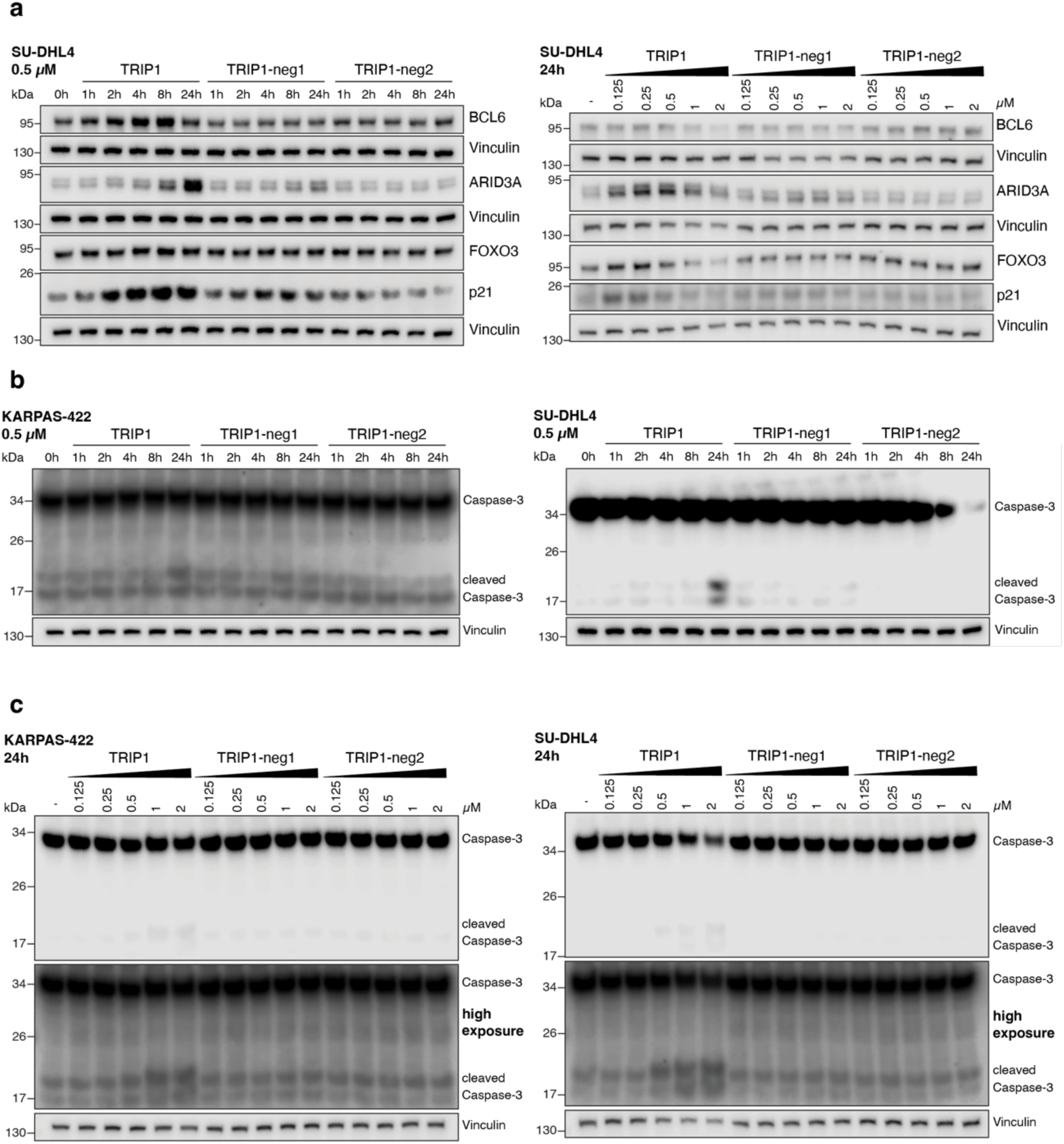
TRIP1 upregulates BCL6 target genes in SU-DHL4 cells and promotes caspase cleavage. **a**, Time-resolved (left) and dose-resolved (right) expression changes of BCL6, ARID3A, FOXO3 and p21 (CDKN1A) in SU-DHL4 cells. **b**, Caspase-3 cleavage following time-resolved TRIP1, TRIP1-neg1, and TRIP1-neg2 treatment in KARPAS-422 (left) and SU-DHL4 (right) cells. **c**, Caspase-3 cleavage following 24h treatment with a dose titration of TRIP1, TRIP1-neg1 and TRIP1-neg2 in KARPAS-422 (left) and SU-DHL4 (right) cells. For **a-c**, loading controls were performed per gel. Western blots are representative of two independent replicates.

**Extended Data Figure 6.**
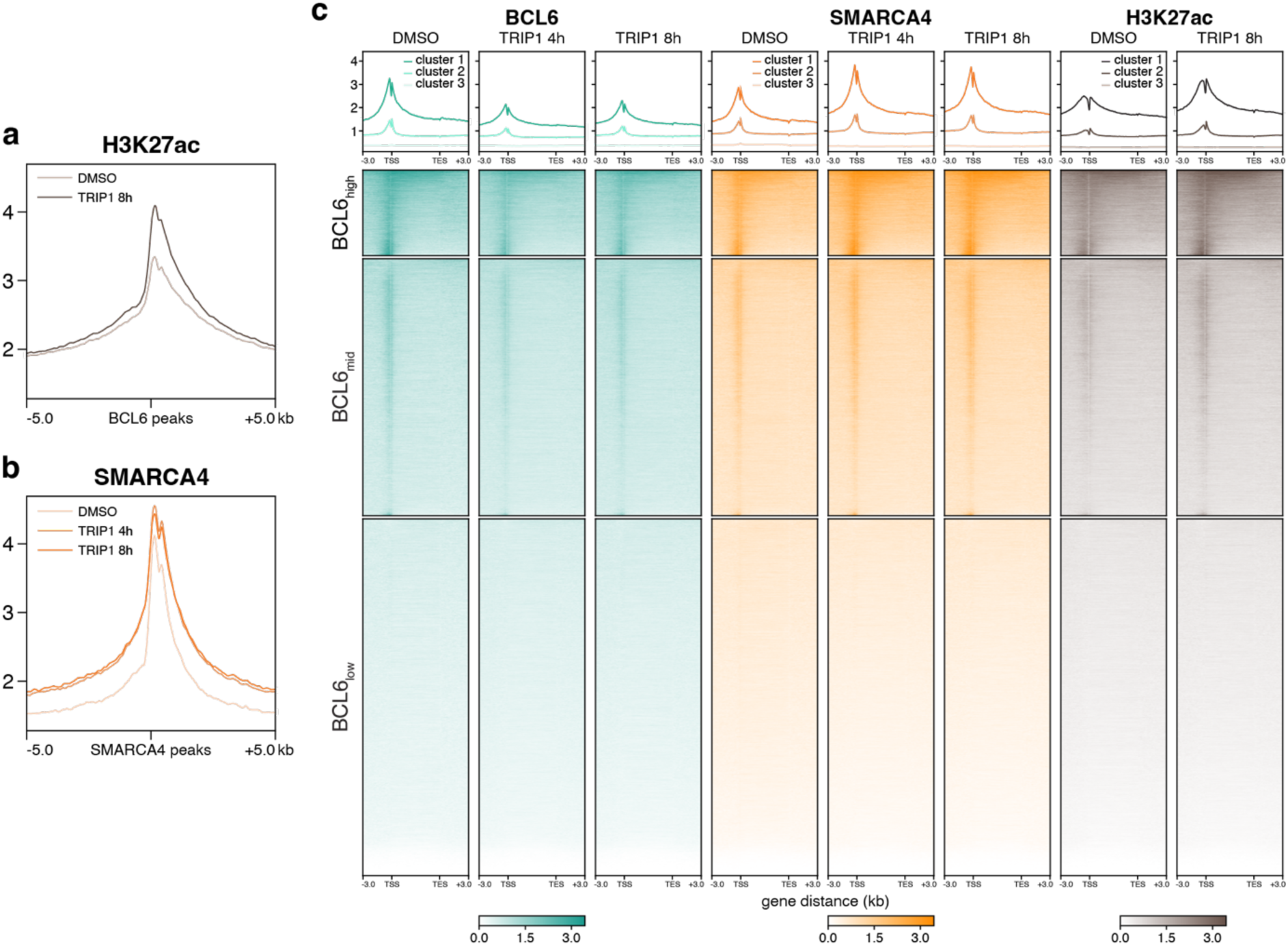
TRIP1-induced chromatin changes at BCL6 binding sites. **a**, **b**, CUT&RUN binding profiles at BCL6 (**a**) or SMARCA4 (**b**) peaks in KARPAS422 cells. **a**, Binding profile of H3K27ac at BCL6 peaks. **b**, Binding profile of SMARCA4 at SMARCA4 peaks. Peaks were called from DMSO treated KARPAS-422 samples. Profiles were centered on peaks and extended 5 kb upstream and downstream of the peak location. **c**, BCL6, SMARCA4 and H3K27ac signal on all annotated genes across TRIP1 conditions. Binding intensities were computed along annotated genes with gene size-normalized signal between transcription start site (TSS) and transcription end site (TES) plus 3 kb up- and downstream of gene borders. Chromatin profiles were k-means clustered into three clusters correlating with BCL6 intensity and termed BCL6 high, mid, or low, respectively (bottom). Binding profiles were calculated to indicate relative signals per gene cluster in DMSO or TRIP1 (2 µM) treated cells after 4 and 8 hours (Top). All CUT&RUN data is from two merged independent replicates.

**Extended Data Figure 7.**
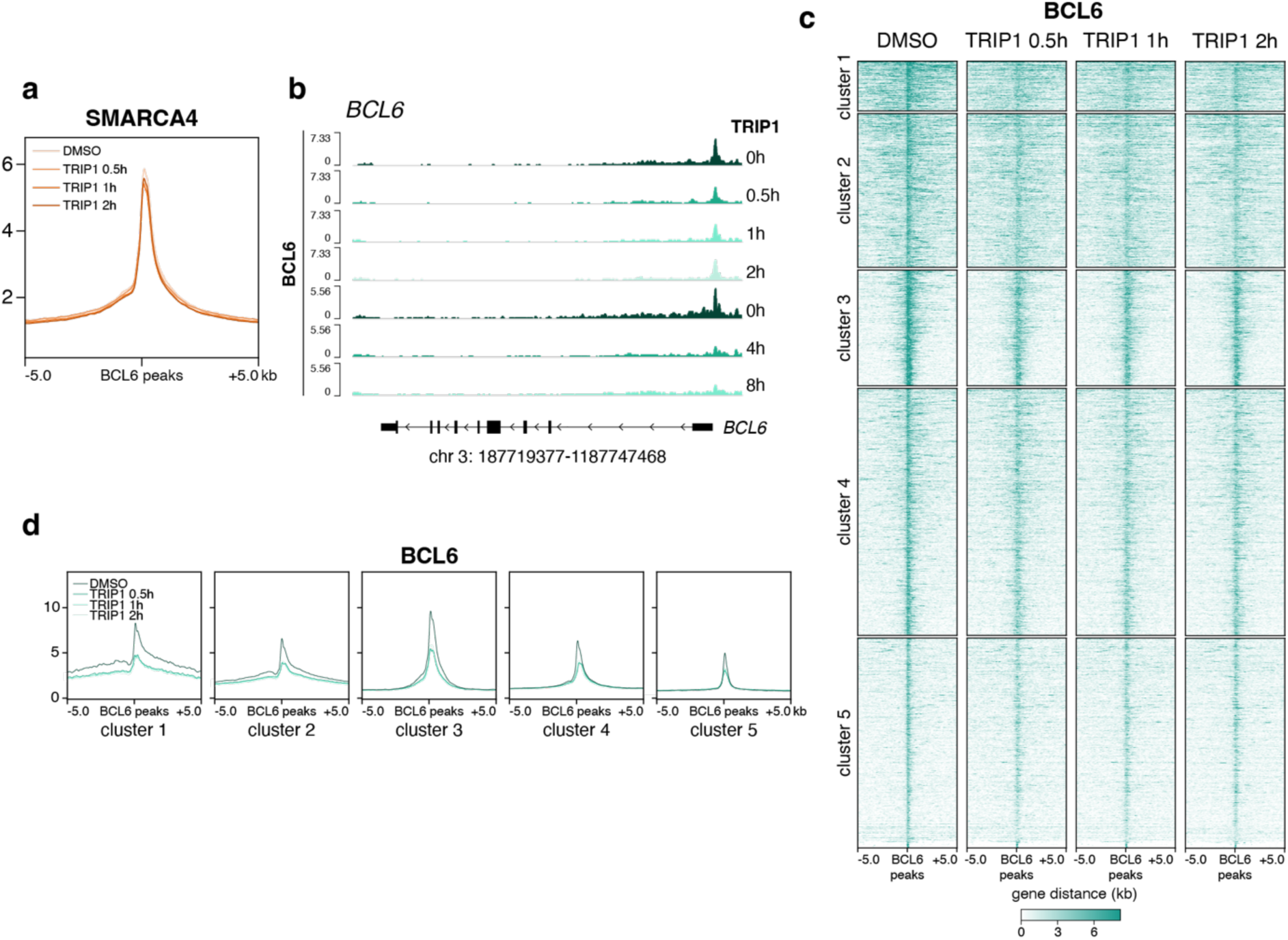
TRIP1 rapidly evicts BCL6 from chromatin. **a**, CUT&RUN binding profiles at BCL6 peaks. SMARCA4 signal at BCL6 peaks was calculated for DMSO and TRIP1-treated cells following 30 min, 1h and 2h of TRIP1 treatment (1 µM). BCL6 peaks were called from DMSO treated KARPAS-422 samples. Profiles were centered on peaks and extended 5 kb upstream and downstream of the peak location. **b**, Genome tracks of the BCL6 gene locus. Time-resolved BCL6 signal is computed along the gene locus. Exon position and genome location are indicated below the genome tracks. **c**, **d**, Changes in BCL6 signal at BCL6 peaks following short TRIP1 (30 minutes, 1 hour, 2 hours) treatment at 1 µM. BCL6 peaks were k-means clustered into five clusters to capture heterogenous BCL6 changes. **c**, Heatmaps of BCL6 signal across time points. **d**, Binding profiles of BCL6 at 5 BCL6 peak clusters from **c**. All CUT&RUN data is from 2 merged independent replicates

**Extended Data Figure 8.**
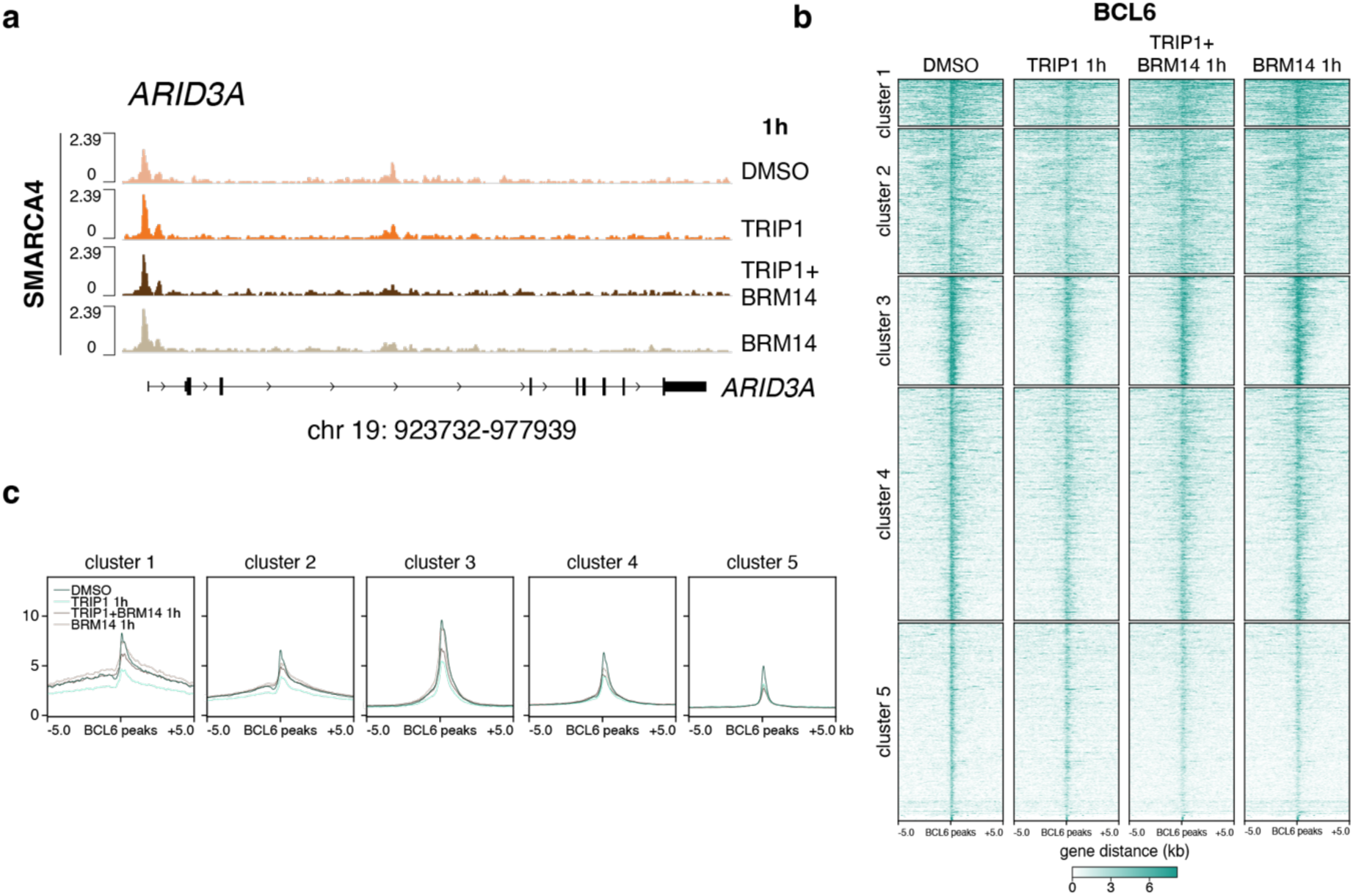
BAF ATPase activity is required for TRIP1-induced targeted chromatin remodeling. **a**, Genome tracks of the ARID3A gene locus following TRIP1 and BRM-014 co-treatment for 1 hour. SMARCA4 signal is computed along the ARID3A gene locus. Exon position and genome location are indicated below the genome tracks. All CUT&RUN data is from two merged independent replicates. **b**, **c**, Changes in BCL6 signal at BCL6 peaks following co-treatment with DMSO or TRIP1 (1 µM) and BRM-014 (1 µM) for 1 hour. BCL6 peaks were k-means clustered into 5 clusters to capture heterogenous BCL6 changes. Signal was centered around peaks and extended ± 5 kb. **b**, Heatmaps of BCL6 signal across conditions. **c**, Binding profiles of BCL6 at 5 BCL6 peak clusters from **b**.

