## Supplementary material for "Leveraging the BAF chromatin remodeling complex for targeted transcriptional rewiring in cancer": Flow Cytometry Gating

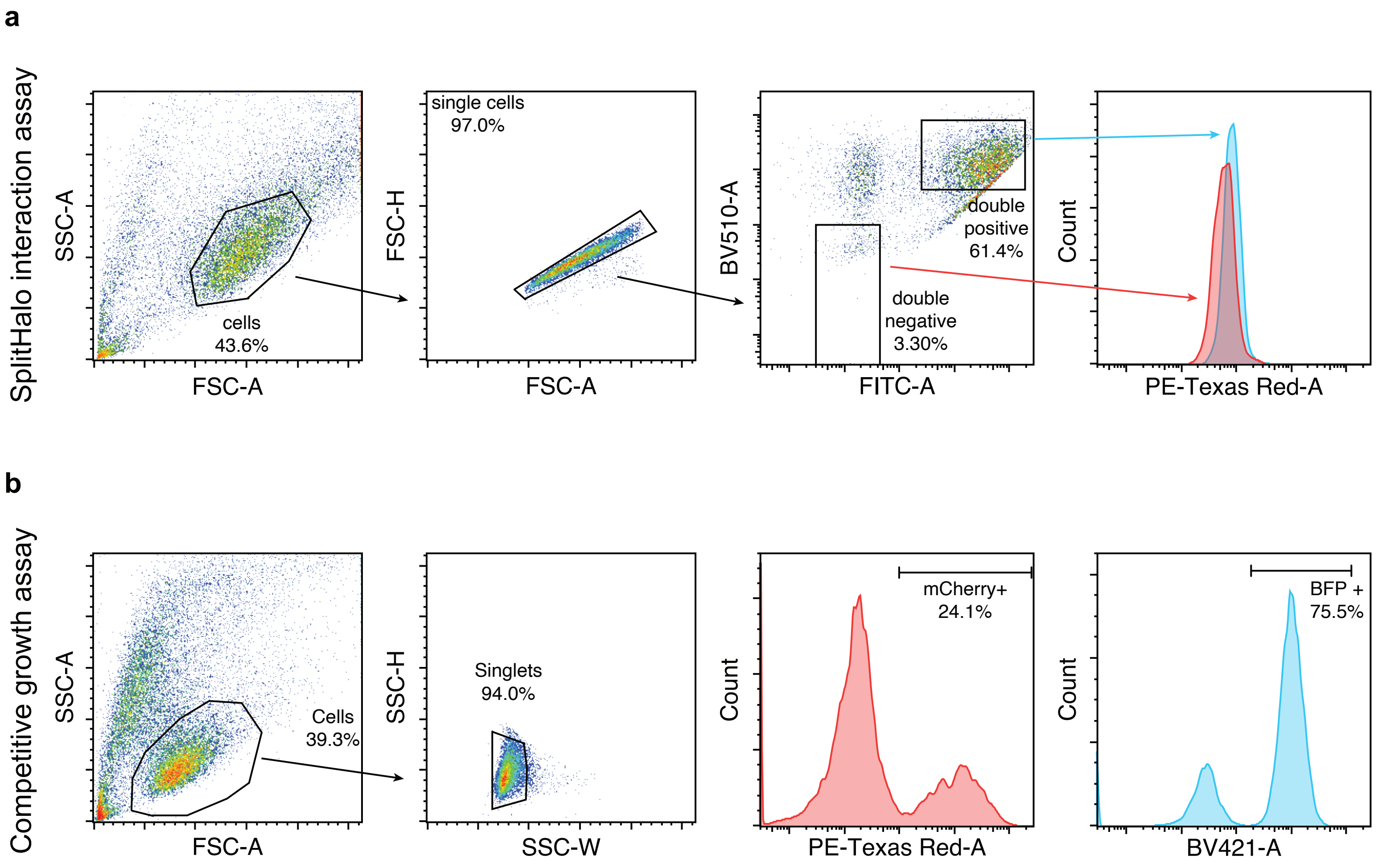


**Supplementary Figure 1**| Gating strategy for flow cytometry-based assays
**a**, SplitHalo interaction assay. **b**, Competative growth assay.
