## Supplementary material for "Leveraging the BAF chromatin remodeling complex for targeted transcriptional rewiring in cancer": Chemical Synthesis

### Chemical synthesis methods

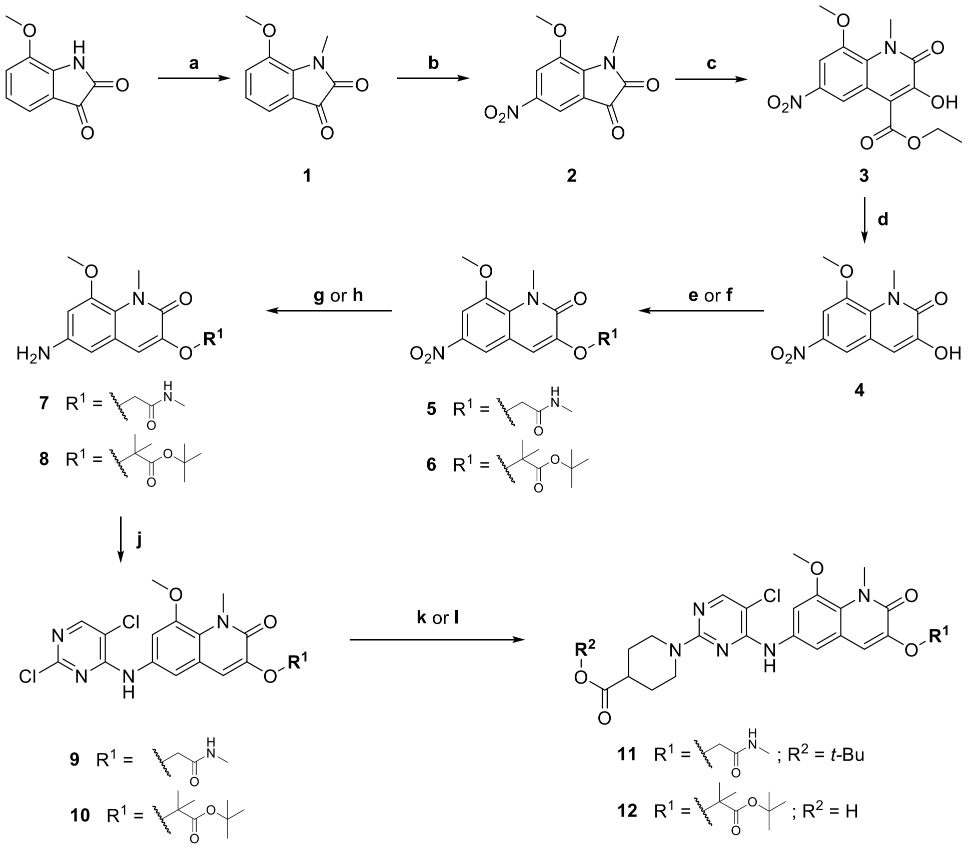

**Scheme 1** Synthesis of the BCL6 inhibitor building block.

*Reagents and conditions:* (a) MeI, $K_2CO_3$, DMF, RT, 2 h, 68%; (b) **1**, $KNO_3$, $H_2SO_4$, 0 °C to RT, 19 h, 64%; (c) **2**, (i) ethyl diazoacetate, DBU, MeOH, RT, 3 h; (ii) dirhodium(II) tetraacetate, MeOH, RT, 16 h, 47%; (d) **3**, NaOH, $H_2O$, 100 °C, 16 h, 71%, (e) **4**, 2-bromo-*N*-methylacetamide, $Cs_2CO_3$, DMF, RT, 90 min, 88%; (f) **4**, *tert*-butyl 2-hydroxy-2-methylpropanoate, TPP, DTAD, THF, 0 °C to RT, 72 h, 25%; (g) **5**, Fe, $H_2O$/EtOH/AcOH, 120 °C, 1 h, 53%; (h) **6**, $H_2$, Pd/C, EtOH, RT, 16 h, 72%; (j) **7** or **8**, 2,4,5-trichloropyrimidine, DIPEA, DMF, 70 °C, 16 h, 86–88%; (k) **9**, *tert*-butyl piperidine-4-carboxylate, DIPEA, DMSO, 100 °C, 16 h, 88%; (l) **10**, piperidine-4-carboxylic acid, DIPEA, DMSO, 100 °C, 16 h, 98%.

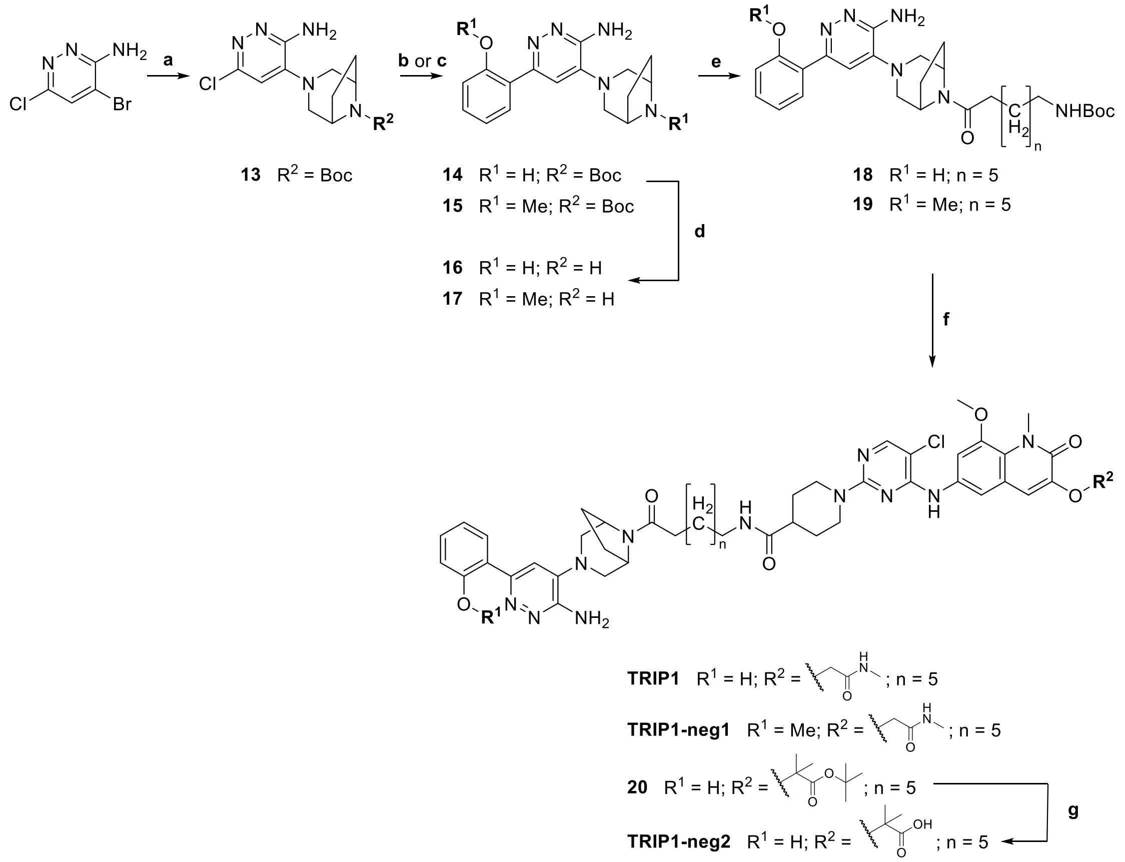

**Scheme 2** Synthesis of TRIP1 and respective negative controls.

*Reagents and conditions:* (a) *Tert*-Butyl 3,8-diazabicyclo[3.2.1]octane-8-carboxylate, DMSO, 90 °C, 16 h, 76%; (b) **13**, 2-(2-methoxyphenyl)-4,4,5,5-tetramethyl-1,3,2-dioxaborolane, PdCl$_2$(dppf) × CH$_2$Cl$_2$, K$_2$CO$_3$, H$_2$O/dioxane, 100 °C, 3 h, 67%; (c) **13**, 2-(4,4,5,5-tetramethyl-1,3,2-dioxaborolan-2-yl)phenol, PdCl$_2$(dppf) × CH$_2$Cl$_2$, K$_2$CO$_3$, H$_2$O/dioxane, 100 °C, 3 h, 89%; (d) **14** or **15**, 0.5M HCl in EtOAc, RT, 16 h, 93%; (e) **16** or **17**, 8-((*tert*-butoxycarbonyl)amino)octanoic acid, HATU, DIPEA, DMF, RT, 16 h, 82–85%; (f) (i) **23**, **24**, or **25**, TFA, CH$_2$Cl$_2$, RT, 2 h; (ii) **11** (deprotected) or **12**, HATU, DIPEA, DMF, RT 16 h, 22–89%; (g) **29**, 1M HCl in EtOAc, RT, 16 h, 83%.

*General Procedure A*

Boc deprotection with TFA. The Boc-protected amine was dissolved in a mixture of dry CH$_2$Cl$_2$ and TFA (1:1, 5 mL). The resulting mixture was stirred at RT for 2 h after which the solvents were evaporated. Next the remaining TFA traces were coevaporated with CH$_2$Cl$_2$ (3 × 10 mL) and the residue was dried under high vacuum for at least one hour.

*General Procedure B*

*Tert*-butyl ester deprotection with TFA. The *tert*-butyl ester was dissolved in a mixture of dry $CH_2Cl_2$ and TFA (1:1, 5 mL). The resulting mixture was stirred at 40 °C for 2 h and after cooling to RT the solvents were evaporated. Next, the remaining TFA traces were coevaporated with $CH_2Cl_2$ (3 × 10 mL), and the residue was dried under high vacuum for at least one hour.

*General Procedure C*

HATU mediated amide coupling. To the (deprotected) carboxylic acid was added first dry DMF (2.5 mL) then DIPEA (4 eq) and lastly HATU (1.2 eq). This mixture was stirred at RT for 15 min before adding a solution of the (deprotected) amine in dry DMF (2.5 mL) and DIPEA (4 eq). The resulting solution was stirred at RT for 16 h.

*7-Methoxy-1-methyl-1H-indole-2,3-dione* (**1**)

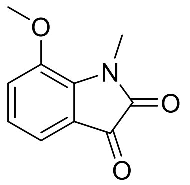

7-Methoxy-1*H*-indole-2,3-dione (10 g, 56.5 mmol) was dissolved in DMF (50 mL) followed by the addition of $K_2CO_3$ (11.7 g, 84.8 mmol). After stirring for 5 min at RT, MeI (4.57 mL, 73.5 mmol) was added, and stirring was continued for 2 h. Water (200 mL) was then added, and the *p*H was adjusted to 1 with concd HCl. After stirring for 30 min, the precipitate was filtered off, washed with $H_2O$ (3 × 20 mL), and the residue was further dried *in vacuo* to yield compound **1** as a dark red solid. Yield: 68% (7.34 g); mp: 167 – 169 °C; $R_f$ = 0.48 (50% EtOAc/petrol ether); **$^1$H NMR** (600 MHz, DMSO-$d_6$) δ 3.33 (d, $J$ = 1.6 Hz, 4H), 3.86 (s, 2H), 7.06 (dd, $J$ = 7.5, 8.2 Hz, 1H), 7.13 (d, $J$ = 7.3 Hz, 1H), 7.38 (d, $J$ = 8.2 Hz, 1H); **$^{13}$C NMR** (151 MHz, DMSO-$d_6$) δ 29.51, 56.78, 116.76, 118.98, 122.93, 124.33, 139.15, 146.02, 158.70, 183.77; **LC-MS** (ESI) (90% $H_2O$ to 100% MeCN in 10 min, then 100% MeCN to 20 min, DAD 220-600 nm), $t_R$ = 4.25 min, 94% purity, $m/z$ [M + H]$^+$ calcd for $C_{10}H_{10}NO_3$, 192.1; found, 192.1.

*7-Methoxy-1-methyl-5-nitro-1H-indole-2,3-dione* (**2**)

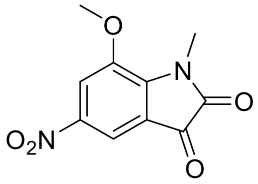

Compound **1** (7.1 g, 37.1 mmol, 1 eq) was dissolved in concentrated $H_2SO_4$ (80 mL). The solution was cooled to 0 °C and $KNO_3$ (3.75 g, 37.1 mmol, 1 eq) was added portion wise. The solution was stirred at 0 °C for 30 min after which it was allowed to warm up to RT and stirring was continued for 18 h. Another portion of $KNO_3$ (748 mg, 7.4 mmol, 0.2 eq) was added and the mixture was stirred for one more hour. The solution was then poured on ice water (300 mL) and stirred for 30 min after which it was extracted with EtOAc (4 × 300 mL). The combined organic phase was washed with sat. NaCl solution (2 × 300 mL), dried over $Na_2SO_4$, filtered and evaporated to yield the crude product which was further purified by column chromatography ($CH_2Cl_2$) to give compound **2** as an orange solid. Yield: 64% (5.6 g); mp: 151 – 153 °C; $R_f$ = 0.45 (50% EtOAc/petrol ether); **$^1$H NMR** (600 MHz, DMSO-$d_6$) δ 3.39 (s, 3H), 4.00 (s, 3H), 7.88 (d, $J$ = 2.1 Hz, 1H), 8.08 (d, $J$ = 2.1 Hz, 1H); **$^{13}$C NMR** (151 MHz, DMSO-*d*6) δ 29.62, 57.38, 111.78, 116.06, 118.52, 143.36, 144.95, 145.96, 159.39, 181.48; **LC-MS** (ESI) (90% $H_2O$ to 100% MeCN in 10 min, then 100% MeCN to 20 min, DAD 220-600 nm), $t_R$ = 4.32 min, 99% purity, *m/z* $[M + H]^+$ calcd for $C_{10}H_9N_2O_5$, 237.1; found, 237.0.

*Ethyl 1,2-dihydro-3-hydroxy-8-methoxy-1-methyl-6-nitro-2-oxo-4-quinolinecarboxylate* (**3**)

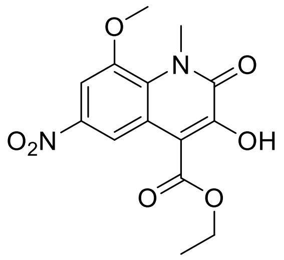

Compound **2** (5.6 g, 23.6 mmol, 1 eq) was dissolved in MeOH (200 mL), DBU (0.7 mL, 4.7 mmol, 0.2 eq) and ethyl diazoacetate (81% solution in $CH_2Cl_2$, 4.99 g, 35,4 mmol, 1.5 eq) were then added and the solution was stirred for 3 h at RT. Rhodium (II) acetate (106 mg, 0.24 mmol, 0.01 eq) was added and the mixture was stirred over night at RT. The precipitation was filtered off, washed with MeOH (3 × 20 mL), and the residue was dried *in vacuo* to yield compound **3** as an off-white solid. Yield: 47% (3.6 g); mp: 185 – 186 °C; $R_f$ = 0.41 (5% MeOH/$CH_2Cl_2$); **$^1$H NMR** (500 MHz, DMSO-$d_6$) δ 1.33 (t, $J$ = 7.1 Hz, 3H), 3.88 (s, 3H), 4.00 (s, 3H), 4.42 (q, $J$ = 7.1 Hz, 2H), 7.76 (d, $J$ = 2.5 Hz, 1H), 7.86 (d, $J$ = 2.4 Hz, 1H), 10.85 (s, 1H); **$^{13}$C NMR** (126 MHz, DMSO-$d_6$) δ 14.21, 36.53, 57.31, 61.79, 104.60, 111.79, 119.55, 129.70, 142.70, 149.15, 159.76, 164.94; **LC-MS** (ESI) (90% $H_2O$ to 100% MeCN in 10 min, then 100% MeCN to 20 min, DAD 220-600 nm), $t_R$ = 5.19 min, 97% purity, *m/z* $[M + H]^+$ calcd for $C_{14}H_{15}N_2O_7$, 323.1; found, 323.1.

*3-Hydroxy-8-methoxy-1-methyl-6-nitro-1,2-dihydroquinolin-2-one* (**4**)

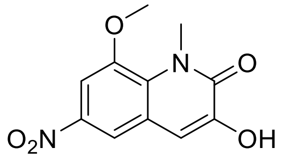

Compound **3** (3.6 g, 11 mmol, 1 eq) and NaOH (880 mg, 22 mmol, 2 eq) were dissolved in water (100 mL) and stirred under reflux for 16 h. After cooling to RT, the *p*H was adjusted to 1 with concd HCl, leading to the formation of a precipitate which was filtered off, washed with water (3 × 20 mL), and the residue was further dried *in vacuo* yielding the crude product which was then purified by flash chromatography (70% EtOAc/cyclohexane) to give compound **4** as a yellow solid. Yield: 71% (1.95 g); thermal decomposition at 290 °C; $R_f$ = 0.54 (2.5% MeOH/$CH_2Cl_2$); **$^1$H NMR** (500 MHz, DMSO-$d_6$) δ 3.89 (s, 3H), 3.97 (s, 3H), 7.24 (s, 1H), 7.67 (d, $J$ = 2.6 Hz, 1H), 8.13 (d, $J$ = 2.5 Hz, 1H), 9.98 (s, 1H); **$^{13}$C NMR** (126 MHz, DMSO-$d_6$) δ 35.66, 57.16, 103.99, 112.04, 115.45, 123.05, 130.00, 142.31, 146.89, 148.55, 159.94; **LC-MS** (ESI) (90% $H_2O$ to 100% MeCN in 10 min, then 100% MeCN to 20 min, DAD 220-600 nm), $t_R$ = 5.43 min, 93% purity, $m/z$ [M + H]$^+$ calcd for $C_{11}H_{11}N_2O_5$, 251.1; found, 251.0.

*2-[(1,2-Dihydro-8-methoxy-1-methyl-6-nitro-2-oxo-3-quinolinyl)oxy]-N-methylacetamide* (**5**)

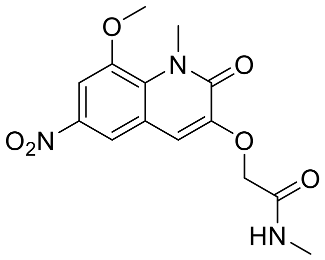


Compound **4** (1.5 g, 6 mmol, 1 eq) and $Cs_2CO_3$ (3.91 g, 12 mmol, 2 eq) were suspended in DMF (30 mL). 2-Bromo-*N*-methylacetamide (1.1 g, 7.2 mmol, 1.2 eq.) was then added and the mixture was stirred for 90 min at RT. Water (60 mL) was added, and the precipitate was filtered off, washed with water (20 mL) and diethyl ether (20 mL), and the residue was further dried *in vacuo* to give compound **5** as a colorless solid. Yield: 88% (1.7 g); mp: 227 – 229 °C; $R_f$ = 0.36 (5% MeOH/$CH_2Cl_2$); **$^1$H NMR** (600 MHz, DMSO-$d_6$) δ 2.66 (d, $J$ = 4.4 Hz, 3H), 3.87 (s, 3H), 3.99 (s, 3H), 4.56 (s, 2H), 7.44 (s, 1H), 7.76 (s, 1H), 7.88 (s, 1H), 8.21 (s, 1H); **$^{13}$C NMR** (151 MHz, DMSO-$d_6$) δ 25.60, 35.61, 57.27, 67.70, 105.17, 112.93, 116.10, 121.61, 131.04, 142.27, 147.47, 148.46, 158.22, 167.04; **LC-MS** (ESI) (90% $H_2O$ to 100% MeCN in 10 min, then 100% MeCN to 20 min, DAD 220-600 nm), $t_R$ = 4.81 min, 99% purity, $m/z$ [M + H]$^+$ calcd for $C_{14}H_{16}N_3O_6$, 322.1; found, 322.1.

*2-[(6-Amino-1,2-dihydro-8-methoxy-1-methyl-2-oxo-3-quinolinyl)oxy]-N-methylacetamide* (**7**)

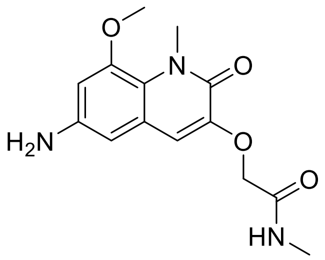


Iron (2.37 g, 42.4 mmol, 8 eq) was suspended in a 1:1 mixture of water and AcOH (100 mL). Compound **5** (1.7 g, 5.3 mmol, 1 eq) was suspended in a mixture of five parts EtOH to two parts AcOH (175 mL). Both suspensions were combined and stirred under reflux for 1 h. After cooling down to RT the solvents were evaporated, and the residue was taken up in sat. $NaHCO_3$ solution (100 mL). The aqueous phase was extracted with 10% MeOH in $CH_2Cl_2$ (3 × 100 mL). The combined organic phase was washed with sat. NaCl solution (100 mL), dried with $Na_2SO_4$, filtered and evaporated to yield the crude product which was further purified by flash chromatography (0–10% MeOH/$CH_2Cl_2$) to give compound **7** as a colorless solid. Yield: 53% (816 mg); mp: 192 – 193 °C; $R_f$ = 0.28 (5% MeOH/$CH_2Cl_2$); **$^1$H NMR** (500 MHz, DMSO-*d*$_6$) δ 2.65 (d, *J* = 4.7 Hz, 3H), 3.76 (s, 3H), 3.78 (s, 3H), 4.48 (s, 2H), 5.05 (s, 2H), 6.27 (d, *J* = 2.4 Hz, 1H), 6.46 (d, *J* = 2.3 Hz, 1H), 6.94 (s, 1H), 7.91 (q, *J* = 4.4 Hz, 1H); **$^{13}$C NMR** (126 MHz, DMSO-*d*$_6$) δ 25.52, 35.04, 56.50, 68.22, 101.15, 103.02, 114.31, 118.01, 122.90, 144.88, 146.45, 148.81, 157.48, 167.81; **LC-MS** (ESI) (90% $H_2O$ to 100% MeCN in 10 min, then 100% MeCN to 20 min, DAD 220-600 nm), $t_R$ = 3.21 min, 96% purity, *m*/*z* $[M + H]^+$ calcd for $C_{14}H_{18}N_3O_4$, 292.1; found, 292.1.

*2-[[6-[(2,5-Dichloro-4-pyrimidinyl)amino]-1,2-dihydro-8-methoxy-1-methyl-2-oxo-3-quinolinyl]oxy]-N-methylacetamide* (**9**)

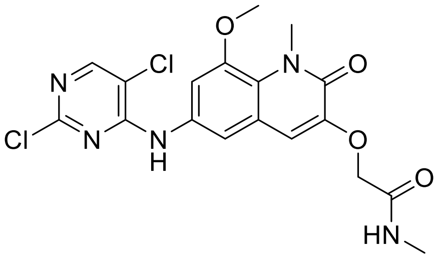


Compound **7** (816 mg, 2.8 mmol, 1 eq), 2,4,5-trichloropyrimidine (770 mg, 4.2 mmol, 1.5 eq) and DIPEA (1.46 mL, 8.4 mmol, 3 eq) were dissolved in dry DMF (10 mL). The solution was purged with argon for 5 min and stirred at 70 °C overnight. Ater cooling down to RT water (40 mL) was added and stirring was continued for 10 min. The formed precipitate was filtered off, washed with water (3 × 10 mL) and dried *in vacuo* to yield compound **9** as an off colorless solid. Yield: 86% (1.05 g); thermal decomposition at 290 °C; $R_f$ = 0.49 (5% MeOH/$CH_2Cl_2$); Due to insufficient solubility no NMR spectrum could be generated of this intermediate; **LC-MS** (ESI) (90% $H_2O$ to

100% MeCN in 10 min, then 100% MeCN to 20 min, DAD 220-600 nm), $t_R$ = 5.44 min, 96% purity, *m/z* $[M + H]^+$ calcd for $C_{18}H_{18}Cl_2N_5O_4$, 438.1; found, 438.1.

*Tert-Butyl 1-[5-chloro-4-({1-methyl-3-[(methylcarbamoyl)methoxy]-2-oxo-1,2-dihydroquinolin-6-yl}amino)pyrimidin-2- yl]piperidine-4-carboxylate* (**11**)

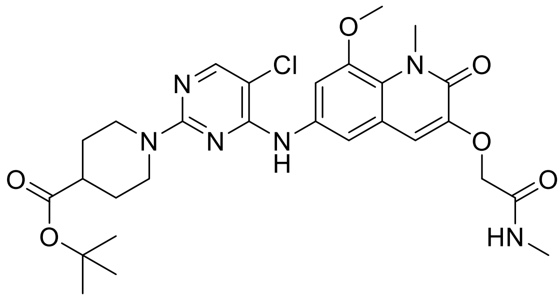

Compound **9** (219 mg, 0.5 mmol, 1 eq) was suspended in dry DMSO (5 mL). *Tert*-Butyl piperidine-4-carboxylate (185 mg, 1 mmol, 2 eq) and DIPEA (261 µL, 1.5 mmol, 3 eq) were added and the mixture was stirred at 100 °C over night. After cooling to RT sat. $NH_4Cl$ (50 mL) was added, and the aqueous phase was extracted with EtOAc (3 × 50 mL). The combined organic phase was washed with 5% LiCl solution (50 mL), sat. NaCl solution (50 mL), dried over $Na_2SO_4$, filtered and evaporated to yield the crude product which was further purified by flash chromatography (0–10% MeOH/$CH_2Cl_2$) to give compound **11** as a pale brown solid. Yield: 88% (258 mg); mp: 216 - 218 °C; $R_f$ = 0.59 (10% MeOH/$CH_2Cl_2$); **$^1$H NMR** (600 MHz, DMSO-$d_6$) δ 1.38 (s, 9H), 1.44 (ddd, *J* = 4.1, 11.2, 13.1 Hz, 2H), 1.82 (dd, *J* = 3.7, 13.0 Hz, 2H), 2.44 – 2.48 (m, 1H), 2.65 (d, *J* = 4.5 Hz, 3H), 3.00 (ddd, *J* = 2.8, 11.4, 13.6 Hz, 2H), 3.85 (s, 3H), 3.86 (s, 3H), 4.38 (ddd, *J* = 3.8, 3.8, 13.3 Hz, 2H), 4.54 (s, 2H), 6.99 (s, 1H), 7.51 (d, *J* = 2.2 Hz, 1H), 7.54 (d, *J* = 2.3 Hz, 1H), 7.92 (q, *J* = 4.7 Hz, 1H), 8.04 (s, 1H), 8.75 (s, 1H); **$^{13}$C NMR** (151 MHz, DMSO-$d_6$) δ 25.54, 27.65, 27.84, 35.17, 41.46, 43.26, 56.77, 68.09, 79.80, 102.23, 107.17, 112.36, 113.87, 121.56, 122.57, 134.48, 146.75, 147.78, 154.82, 155.39, 157.92, 159.24, 167.62, 173.66; **LC-MS** (ESI) (90% $H_2O$ to 100% MeCN in 10 min, then 100% MeCN to 20 min, DAD 220-600 nm), $t_R$ = 7.85 min, 99% purity, *m/z* $[M + H]^+$ calcd for $C_{28}H_{36}ClN_6O_6$, 587.2; found, 587.3, **HRMS** (ESI) *m/z* $[M + H]^+$ calcd for $C_{28}H_{36}ClN_6O_6$, 587.2379; found 587.2361.

*Tert-Butyl 3-(3-amino-6-chloropyridazin-4-yl)-3,8-diazabicyclo[3.2.1]octane-8-carboxylate* (**13**)

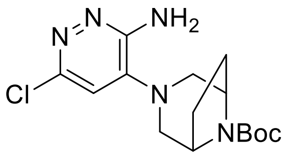

3-Amino-4-bromo-6-chloropyridazine (2.71 g, 13 mmol, 1 eq) was dissolved in dry DMSO (40 mL). After adding tert-butyl 3,8-diazabicyclo[3.2.1]octane-8-carboxylate (8.28 g, 39 mmol, 3 eq) the resulting solution was stirred at 90 °C for 16 h. The solution was cooled to RT and poured into water (200 mL) forming a precipitate which was filtered off to yield the crude product which was purified by column chromatography (EtOAc) to give compound **13** as a colorless solid. Yield: 76% (3.4 g); $R_f$ = 0.48 (EtOAc); **$^1$H NMR** (600 MHz, DMSO-$d_6$) δ 1.41 (s, 9H), 1.80 (d, $J$ = 8.7 Hz, 2H), 1.98 (d, $J$ = 7.7 Hz, 2H), 2.74 (d, $J$ = 11.1 Hz, 2H), 3.25 (d, $J$ = 2.5 Hz, 1H), 3.27 (d, $J$ = 2.8 Hz, 1H), 4.17 (s, 2H), 5.80 (s, 2H), 6.95 (s, 1H); **$^{13}$C NMR** (151 MHz, DMSO-$d_6$) δ 26.88, 27.47, 27.48, 28.23, 40.24, 52.88, 52.98, 53.86, 79.11, 115.19, 141.01, 146.66, 152.96, 155.30, 155.34; **LC-MS** (ESI) (90% $H_2O$ to 100% MeCN in 10 min, then 100% MeCN to 20 min, DAD 220-600 nm), $t_R$ = 7.38 min, 99% purity, $m/z$ [M + H]$^+$ calcd for $C_{15}H_{23}ClN_5O_2$, 340.2; found, 340.3.

*Tert-Butyl 3-(3-amino-6-(2-hydroxyphenyl)pyridazin-4-yl)-3,8-diazabicyclo[3.2.1]octane-8-carboxylate* (**14**)

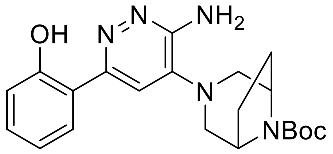


Compound **13** (3.14 g, 9.86 mmol, 1 eq), $K_2CO_3$ (4.15 g, 30 mmol, 3 eq) and 2-(4,4,5,5-tetramethyl-1,3,2-dioxaborolan-2-yl)phenol (4.40 g, 20 mmol, 2 eq) were dissolved in dioxane/water (3:1, 100 mL). The solution was thoroughly purged with argon and 1,1'-bis(diphenylphosphino)ferrocene-palladium(II)dichloride dichloro-methane complex (817 mg, 1 mmol, 0.1 eq) was added. The solution was heated to reflux and stirred under argon atmosphere for 3 h. After cooling to RT, EtOAc (200 mL) was added, the phases were separated and the organic phase was washed with sat. $NH_4Cl$ solution (200 mL), sat. NaCl solution (200 mL), dried with $Na_2SO_4$, filtered and evaporated to yield the crude product, which was purified by flash chromatography (20–60% EtOAc/cyclohexane) to give compound **14** as a yellow solid. Yield: 67% (2.6 g); $R_f$ = 0.51 (60% EtOAc/cyclohexane); **$^1$H NMR** (600 MHz, DMSO-$d_6$) δ 1.42 (s, 9H), 1.82 – 1.86 (m, 2H), 2.01 – 2.05 (m, 2H), 2.86 – 2.91 (m, 2H), 3.31 – 3.36 (m, 2H), 4.23 (s, 2H), 5.95 (s, 2H), 6.84 – 6.90 (m, 2H), 7.23 (dd, $J$ = 7.7, 7.7 Hz, 1H), 7.54 (s, 1H), 7.96 (d, $J$ = 7.8 Hz, 1H), 14.10 (s, 1H); **$^{13}$C NMR** (151 MHz, DMSO-$d_6$) δ 20.90, 28.27, 40.24, 59.89, 79.07, 111.37, 117.51, 117.89, 118.64, 126.59, 130.37, 140.62, 152.88, 153.82, 154.60, 158.67; **LC-MS** (ESI) (90% $H_2O$ to 100% MeCN in 10 min, then 100% MeCN to 20 min, DAD 220-600 nm), $t_R$ = 7.59 min, 99% purity, $m/z$ [M + H]$^+$ calcd for $C_{21}H_{28}N_5O_3$, 398.2; found, 398.6.

*2-(6-Amino-5-(3,8-diazabicyclo[3.2.1]octan-3-yl)pyridazin-3-yl)phenol dihydrochloride* (**16**)

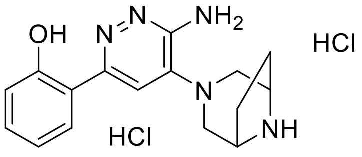

Compound **14** (2.58 g, 6.5 mmol, 1 eq) was suspended in dry EtOAc (25 mL). To this suspension was added 1M HCl in EtOAc (25 mL, 3.85 eq) and the mixture was stirred at RT for 16 h. The precipitate was then filtered off, washed with dry EtOAc (2 × 25 mL), $Et_2O$ (2 × 25 mL) and dried under vacuum to yield compound **16** as a colorless solid. Yield: 93% (2.25 g); **LC-MS** (ESI) (90% $H_2O$ to 100% MeCN in 10 min, then 100% MeCN to 20 min, DAD 220-600 nm), $t_R$ = 3.98 min, 99% purity, *m*/*z* $[M + H]^+$ calcd for $C_{16}H_{20}N_5O$, 298.2; found, 298.3.

*Tert-Butyl (8-(3-(3-amino-6-(2-hydroxyphenyl)pyridazin-4-yl)-3,8-diazabicyclo[3.2.1]octan-8-yl)-8-oxooctyl)carbamate* (**18**)

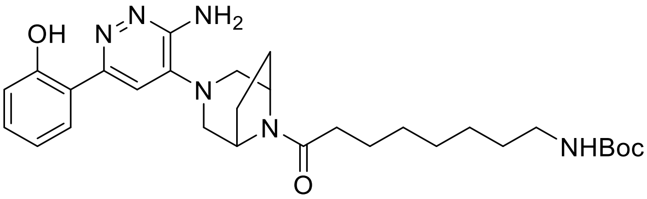

8-((*Tert*-Butoxycarbonyl)amino)octanoic acid (0.13 g, 0.5 mmol, 1 eq) and Compound **16** (0.19 g, 1.0 mmol, 1 eq) were used according to General Procedure C. The reaction was quenched with sat. $NH_4Cl$ (50 mL) and the aqueous phase was extracted with EtOAc (3 × 50 mL). The combined organic layer was washed with 5% LiCl solution (50 mL), sat. NaCl solution (50 mL), dried with $Na_2SO_4$, filtered and evaporated to yield the crude product which was purified by flash chromatography (0–10% MeOH/$CH_2Cl_2$) to give compound **18** as a colorless solid. Yield: 85% (228 mg); mp: 140 – 141 °C $R_f$ = 0.24 (5% MeOH/$CH_2Cl_2$); **$^1$H NMR** (500 MHz, DMSO-*d*$_6$) δ 1.18 – 1.33 (m, 7H), 1.31 – 1.40 (m, 11H), 1.47 – 1.57 (m, 2H), 1.71 – 1.82 (m, 1H), 1.87 – 1.99 (m, 1H), 1.99 – 2.09 (m, 1H), 2.07 – 2.16 (m, 1H), 2.20 – 2.31 (m, 1H), 2.31 – 2.41 (m, 1H), 2.84 – 2.94 (m, 4H), 3.31 (d, *J* = 2.7 Hz, 1H), 3.33 – 3.40 (m, 1H), 4.40 (d, *J* = 6.7 Hz, 1H), 4.61 (d, *J* = 7.0 Hz, 1H), 5.98 (s, 1H), 6.71 (t, *J* = 5.7 Hz, 1H), 6.83 – 6.91 (m, 2H), 7.23 (ddd, *J* = 1.6, 7.1, 8.5 Hz, 1H), 7.56 (s, 1H), 7.92 (dd, *J* = 1.6, 7.9 Hz, 1H), 14.08 (s, 1H); **$^{13}$C NMR** (126 MHz, DMSO-*d*$_6$) δ 24.81, 26.36, 28.09, 28.41, 28.71, 28.94, 29.59, 33.01, 40.30, 50.91, 53.54, 54.05, 54.50, 77.40, 111.78, 117.53, 117.86, 118.62, 126.46, 130.37, 140.55, 153.78, 154.72, 155.72, 158.66, 168.50; **LC-MS** (ESI) (90% $H_2O$ to 100% MeCN in 10 min, then 100% MeCN to 20 min, DAD 220-600 nm), $t_R$ = 7.98 min, 96% purity, *m*/*z* $[M + H]^+$ calcd for $C_{29}H_{43}N_6O_4$, 539.3; found, 539.5; **HRMS** (ESI) *m*/*z* $[M + H]^+$ calcd for $C_{29}H_{43}N_6O_4$, 539.3340; found, 539.3355.

*N-(8-(3-(3-Amino-6-(2-hydroxyphenyl)pyridazin-4-yl)-3,8-diazabicyclo[3.2.1]octan-8-yl)-8-oxooctyl)-1-(5-chloro-4-((8-methoxy-1-methyl-3-(2-(methylamino)-2-oxoethoxy)-2-oxo-1,2-dihydroquinolin-6-yl)amino)pyrimidin-2-yl)piperidine-4-carboxamide* (**TRIP1**)

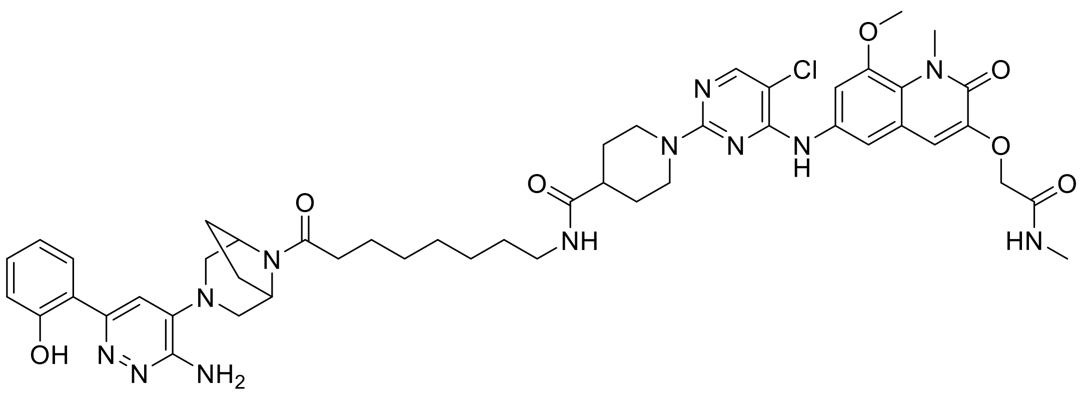

Compound **11** (81 mg, 0.15 mmol, 1 eq) was deprotected according to General Procedure B. Compound **18** (88 mg, 0.11 mmol, 1 eq) was deprotected according to General Procedure A. The deprotected carboxylic acid and deprotected amine were used as described in General Procedure C. The reaction was quenched by adding water (30 mL). The resulting precipitate was filtered off, washed with water (3 × 30 mL), EtOAc (2 × 30 mL) and dried under vacuum to give the crude product which was purified by flash chromatography (0–10% MeOH/$CH_2Cl_2$) yielding **TRIP1** as a colorless solid. Yield: 51% (73 mg); mp: 173 – 174 °C; $R_f$ = 0.52 (10% MeOH/$CH_2Cl_2$); **$^1$H NMR** (600 MHz, DMSO-$d_6$) δ 1.21 – 1.31 (m, 6H), 1.32 – 1.40 (m, 2H), 1.42 – 1.55 (m, 4H), 1.68 (dd, *J* = 3.7, 13.4 Hz, 2H), 1.75 (ddd, *J* = 5.7, 5.7, 16.5 Hz, 1H), 1.91 (ddd, *J* = 4.5, 10.2, 11.1 Hz, 1H), 2.00 (ddd, *J* = 4.2, 9.4, 10.9 Hz, 1H), 2.04 – 2.12 (m, 1H), 2.22 – 2.30 (m, 1H), 2.31 – 2.38 (m, 2H), 2.64 (d, *J* = 4.6 Hz, 3H), 2.83 – 2.95 (m, 3H), 2.95 – 3.03 (m, 3H), 3.09 – 3.16 (m, 1H), 3.57 – 3.65 (m, 1H), 3.84 (s, 3H), 3.86 (s, 3H), 4.39 (d, *J* = 6.8 Hz, 1H), 4.48 – 4.55 (m, 4H), 4.59 (d, *J* = 7.1 Hz, 1H), 6.25 (s, 2H), 6.86 – 6.94 (m, 2H), 6.98 (s, 1H), 7.24 – 7.29 (m, 1H), 7.51 – 7.55 (m, 3H), 7.74 (t, *J* = 5.7 Hz, 1H), 7.80 (d, *J* = 7.8 Hz, 1H), 7.94 (q, *J* = 4.7 Hz, 1H), 8.04 (s, 1H), 8.75 (s, 1H), 13.82 (s, 1H); **$^{13}$C NMR** (151 MHz, DMSO-$d_6$) δ 24.77, 25.53, 26.39, 28.18, 28.67, 28.89, 29.21, 32.97, 35.17, 38.45, 42.26, 43.69, 50.90, 53.51, 54.01, 54.50, 56.76, 68.07, 102.13, 107.16, 112.32, 113.88, 116.42, 117.29, 117.79, 118.41, 118.89, 121.56, 122.56, 127.44, 130.98, 134.52, 146.74, 147.77, 154.16, 154.75, 155.39, 157.83, 157.92, 158.03, 159.19, 167.63, 168.56, 173.95; **LC-MS** (ESI) (90% $H_2O$ to 100% MeCN in 10 min, then 100% MeCN to 20 min, DAD 220-600 nm), $t_R$ = 6.61 min, 98% purity, *m*/*z* [M + H]$^+$ calcd for $C_{48}H_{60}ClN_{12}O_7$, 951.4; found, 951.7, **HRMS** (ESI) *m*/*z* [M + H]$^+$ calcd for $C_{48}H_{60}ClN_{12}O_7$, 951.4391; found, 951.4410.

*Tert-Butyl 3-(3-amino-6-(2-methoxyphenyl)pyridazin-4-yl)-3,8-diazabicyclo[3.2.1]octane-8-carboxylate* (**15**)

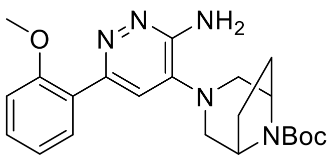

Compound **13** (1.7 g, 5 mmol, 1 eq), $K_2CO_3$ (2.07 g, 15 mmol, 3 eq) and 2-(2-methoxyphenyl)-4,4,5,5-tetramethyl-1,3,2-dioxaborolane (1.52 g, 10 mmol, 2 eq) were dissolved in dioxane/water (3:1, 100 mL). The solution was thoroughly purged with argon and 1,1'-bis(diphenylphosphino)ferrocene-palladium(II)dichloride dichloromethane complex (408 mg, 0.5 mmol, 0.1 eq) was added. The solution was heated to reflux and stirred under argon atmosphere for 3 h. After cooling to RT, EtOAc (200 mL) was added, the phases were separated and the organic phase was washed with sat. $NH_4Cl$ solution (200 mL), sat. NaCl solution (200 mL), dried with $Na_2SO_4$, filtered and evaporated to yield the crude product, which was purified by flash chromatography (0–10% MeOH/$CH_2Cl_2$) to give compound **15** as a beige solid. Yield: 89% (1.8 g); $R_f$ = 0.48 (10% MeOH/$CH_2Cl_2$); **$^1$H NMR** (500 MHz, DMSO-$d_6$) δ 1.41 (s, 9H), 1.80 – 1.86 (m, 2H), 2.03 (d, $J$ = 7.5 Hz, 2H), 2.71 (d, $J$ = 11.2 Hz, 2H), 3.23 (dd, $J$ = 2.6, 11.8 Hz, 2H), 3.78 (s, 3H), 4.20 (s, 2H), 5.64 (s, 2H), 7.02 (ddd, $J$ = 1.0, 7.4, 7.4 Hz, 1H), 7.11 (dd, $J$ = 1.0, 8.4 Hz, 1H), 7.17 (s, 1H), 7.37 (ddd, $J$ = 1.8, 7.4, 8.3 Hz, 1H), 7.59 (dd, $J$ = 1.8, 7.5 Hz, 1H); **$^{13}$C NMR** (126 MHz, DMSO-$d_6$) δ 14.02, 21.85, 28.21, 53.32, 55.71, 79.06, 111.98, 116.37, 120.68, 127.03, 129.86, 130.14, 137.12, 151.51, 153.17, 154.58, 156.62; **LC-MS** (ESI) (90% $H_2O$ to 100% MeCN in 10 min, then 100% MeCN to 20 min, DAD 220-600 nm), $t_R$ = 6.69 min, 99% purity, $m/z$ $[M + H]^+$ calcd for $C_{22}H_{30}N_5O_3$, 412.2; found, 412.4.

*4-(3,8-Diazabicyclo[3.2.1]octan-3-yl)-6-(2-methoxyphenyl)pyridazin-3-amine dihydrochloride* (**17**)

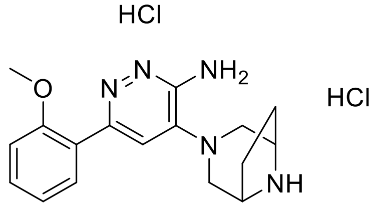

Compound **15** (1.65 g, 4.0 mmol, 1 eq) was suspended in 1M HCl in EtOAc (25 mL, 6.25 eq) and the mixture was stirred at RT for 16 h. The precipitate was then filtered off, washed with dry EtOAc (2 × 25 mL), $Et_2O$ (2 × 25 mL) and dried under vacuum to yield compound **17** as a colorless solid. Yield: 93% (1.42 g); **$^1$H NMR** (600 MHz, DMSO-$d_6$) δ 1.95 – 2.02 (m, 2H), 2.13 – 2.19 (m, 2H), 3.51 – 3.56 (m, 2H), 3.68 – 3.73 (m, 1H), 3.84 (s, 3H), 4.12 (d, $J$ = 4.8 Hz, 2H), 7.13 (dd, $J$ = 7.5, 7.5 Hz, 1H), 7.23 (d, $J$ = 8.3 Hz, 1H), 7.52 (s, 1H), 7.53 – 7.59 (m, 2H), 9.82 (s, 1H), 10.04 (d, $J$ = 9.8 Hz, 1H); **$^{13}$C NMR** (151 MHz, DMSO-$d_6$) δ 24.84, 50.74, 53.84, 56.07, 112.34, 120.99, 130.58, 132.79, 152.32, 156.85; **LC-MS** (ESI) (90% $H_2O$ to 100% MeCN in 10 min, then 100%

MeCN to 20 min, DAD 220-600 nm), $t_R$ = 3.51 min, 99% purity, *m/z* $[M + H]^+$ calcd for $C_{17}H_{22}N$, 312.2; found, 312.1.

*Tert-Butyl (8-(3-(3-amino-6-(2-methoxyphenyl)pyridazin-4-yl)-3,8-diazabicyclo[3.2.1]octan-8-yl)-8-oxooctyl)carbamate* (**19**)

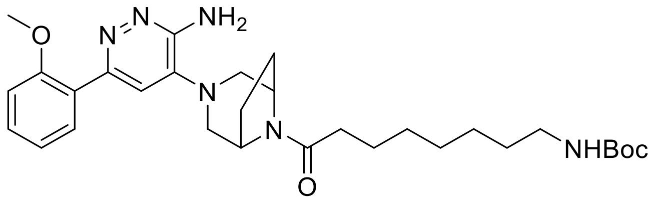


8-((*Tert*-Butoxycarbonyl)amino)octanoic acid (78 m g, 0.3 mmol, 1 eq) and Compound **17** (115 mg, 0.3 mmol, 1 eq) were used according to General Procedure C. The reaction was quenched with sat. $NH_4Cl$ (50 mL) and the aqueous phase was extracted with EtOAc (3 × 50 mL). The combined organic layer was washed with 5% LiCl solution (50 mL), sat. NaCl solution (50 mL), dried with $Na_2SO_4$, filtered and evaporated to yield the crude product which was purified by flash chromatography (0–10% MeOH/$CH_2Cl_2$) to give compound **19** as a colorless solid. Yield: 82% (133 mg); $R_f$ = 0.48 (10% MeOH/$CH_2Cl_2$); **$^1$H NMR** (600 MHz, DMSO-$d_6$) δ 1.17 – 1.27 (m, 6H), 1.29 – 1.39 (m, 11H), 1.43 – 1.54 (m, 2H), 1.70 – 1.79 (m, 1H), 1.86 – 1.95 (m, 1H), 1.97 – 2.04 (m, 1H), 2.05 – 2.13 (m, 1H), 2.21 – 2.29 (m, 1H), 2.29 – 2.37 (m, 1H), 2.70 (d, *J* = 11.2 Hz, 1H), 2.78 (d, *J* = 11.3 Hz, 1H), 2.83 – 2.89 (m, 2H), 3.10 – 3.30 (m, 2H), 3.79 (s, 3H), 4.38 (d, *J* = 6.7 Hz, 1H), 4.59 (d, *J* = 7.0 Hz, 1H), 5.77 (s, 2H), 6.70 (t, *J* = 5.7 Hz, 1H), 7.03 (dd, *J* = 7.4, 7.4 Hz, 1H), 7.12 (d, *J* = 8.3 Hz, 1H), 7.18 (s, 1H), 7.39 (dd, *J* = 7.9, 7.9 Hz, 1H), 7.57 (d, *J* = 7.5 Hz, 1H); **$^{13}$C NMR** (151 MHz, DMSO-$d_6$) δ 24.86, 26.37, 26.42, 28.12, 28.44, 28.73, 28.93, 29.61, 32.95, 40.24, 51.00, 53.42, 54.18, 54.49, 55.82, 77.44, 112.03, 116.43, 120.74, 126.30, 130.21, 130.24, 137.89, 151.15, 154.43, 155.75, 156.67, 168.59; **LC-MS** (ESI) (90% $H_2O$ to 100% MeCN in 10 min, then 100% MeCN to 20 min, DAD 220-600 nm), $t_R$ = 6.69 min, 96% purity, *m/z* $[M + H]^+$ calcd for $C_{30}H_{45}N_6O_4$, 553.4; found, 553.3.

*N-(8-(3-(3-Amino-6-(2-methoxyphenyl)pyridazin-4-yl)-3,8-diazabicyclo[3.2.1]octan-8-yl)-8-oxooctyl)-1-(5-chloro-4-((8-methoxy-1-methyl-3-(2-(methylamino)-2-oxoethoxy)-2-oxo-1,2-dihydroquinolin-6-yl)amino)pyrimidin-2-yl)piperidine-4-carboxamide* (**TRIP1-neg1**)

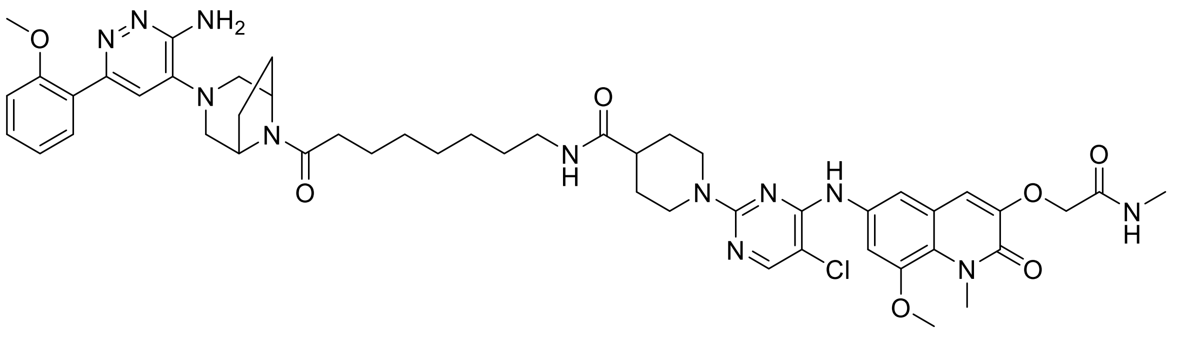

Compound **11** (59 mg, 0.1 mmol, 1 eq) was deprotected according to General Procedure B. Compound **25** (55 mg, 0.1 mmol, 1 eq) was deprotected according to General Procedure A. The deprotected carboxylic acid and deprotected amine were used as described in General Procedure C. The reaction was quenched by adding water (30 mL). The resulting precipitate was filtered off, washed with water (3 × 30 mL), EtOAc (2 × 30 mL) and dried under vacuum to give the crude product which was purified by flash chromatography (0 - 10% MeOH/$CH_2Cl_2$) yielding **TRIP1-neg1** as a colorless solid. Yield: 22% (21 mg); $R_f$ = 0.45 (10% MeOH/$CH_2Cl_2$); **$^1$H NMR** (600 MHz, DMSO-$d_6$) δ 1.23 – 1.31 (m, 6H), 1.32 – 1.39 (m, 2H), 1.44 – 1.51 (m, 4H), 1.67 (d, $J$ = 12.6 Hz, 2H), 1.71 – 1.80 (m, 1H), 1.85 – 1.94 (m, 1H), 1.96 – 2.03 (m, 1H), 2.04 – 2.11 (m, 1H), 2.20 – 2.28 (m, 1H), 2.28 – 2.37 (m, 1H), 2.64 (d, $J$ = 4.6 Hz, 3H), 2.70 (d, $J$ = 11.5 Hz, 1H), 2.79 (d, $J$ = 11.5 Hz, 1H), 2.86 (t, $J$ = 12.7 Hz, 2H), 2.96 – 3.02 (m, 1H), 3.07 – 3.15 (m, 2H), 3.55 – 3.63 (m, 2H), 3.78 (s, 3H), 3.84 (s, 3H), 3.86 (s, 3H), 4.37 (d, $J$ = 6.7 Hz, 1H), 4.52 (d, $J$ = 22.7 Hz, 4H), 4.58 (d, $J$ = 6.9 Hz, 1H), 5.82 (s, 2H), 6.97 – 7.05 (m, 2H), 7.11 (d, $J$ = 8.4 Hz, 1H), 7.18 (s, 1H), 7.38 (t, $J$ = 7.9 Hz, 1H), 7.51 – 7.59 (m, 3H), 7.73 (t, $J$ = 5.4 Hz, 1H), 7.93 – 7.97 (m, 1H), 8.04 (s, 1H), 8.75 (s, 1H); **$^{13}$C NMR** (151 MHz, DMSO-$d_6$) δ 24.80, 25.53, 26.38, 28.08, 28.18, 28.68, 28.86, 29.21, 32.90, 35.17, 38.45, 41.81, 42.26, 43.69, 50.96, 53.39, 54.13, 54.45, 55.80, 56.77, 68.07, 102.12, 107.16, 112.00, 112.32, 113.87, 116.41, 120.71, 121.57, 122.56, 130.23, 134.54, 146.75, 147.77, 151.02, 154.33, 154.82, 155.39, 156.65, 157.92, 159.23, 167.63, 168.55, 173.94; **LC-MS** (ESI) (90% $H_2O$ to 100% MeCN in 10 min, then 100% MeCN to 20 min, DAD 220-600 nm), $t_R$ = 7.50 min, 99% purity, $m/z$ $[M + H]^+$ calcd for $C_{49}H_{62}ClN_{12}O_7$, 965.5; found, 965.5.

*Tert-Butyl 2-((8-methoxy-1-methyl-6-nitro-2-oxo-1,2-dihydroquinolin-3-yl)oxy)-2-methylpropanoate* (**7**)

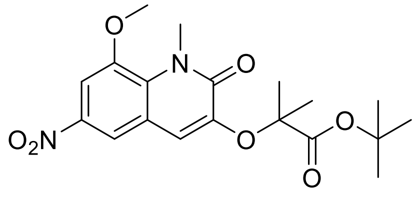

Compound **5** (250 mg, 1 mmol, 1 eq), *tert*-butyl 2-hydroxy-2-methylpropanoate (160 mg, 1 mmol, 1 eq), triphenylphosphine (262 mg, 1 mmol, 1 eq) was weighed into a Schlenk flask and put under argon atmosphere. Dry THF (3 mL) was added via syringe. The resulting suspension was cooled to 0 °C before a solution of di-*tert*-butyl azodicarboxylate in dry THF (3 mL) was added dropwise. The mixture was allowed to warm up to RT and stirred for 72 h. The reaction was quenched by adding 1M NaOH (50 mL). The aqueous phase was extracted with EtOAc (3 × 50 mL). The combined organic phase was washed with sat. NaCl solution (50 mL), dried with $Na_2SO_4$, filtered and evaporated to yield the crude product which was purified by flash chromatography to give compound **7** as a yellow semi-solid. Yield: 25% (100 mg); $R_f$ = 0.35 (50% EtOAc/cyclohexane); **$^1$H NMR** (500 MHz, DMSO-$d_6$) δ 1.38 (s, 9H), 1.59 (s, 6H), 3.85 (s, 3H), 3.99 (s, 3H), 7.14 (s, 1H), 7.77 (d, *J* = 2.6 Hz, 1H), 8.29 (d, *J* = 2.5 Hz, 1H); **$^{13}$C NMR** (126 MHz, DMSO-$d_6$) δ 25.02, 27.53, 35.73, 57.28, 80.68, 81.64, 105.31, 116.25, 116.63, 121.27, 131.19, 142.27, 144.94, 148.37, 158.98, 171.06; **LC-MS** (ESI) (90% $H_2O$ to 100% MeCN in 10 min, then 100% MeCN to 20 min, DAD 220-600 nm), $t_R$ = 5.51 min, 98% purity, *m/z* $[M + H]^+$ calcd for $C_{19}H_{25}N_2O_7$, 393.4; found, 393.2.

*Tert-Butyl 2-((6-amino-8-methoxy-1-methyl-2-oxo-1,2-dihydroquinolin-3-yl)oxy)-2-methylpropanoate* (**9**)

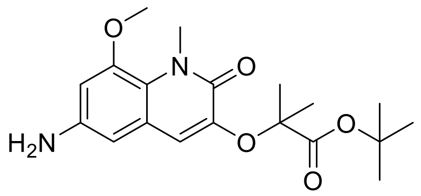

Pd/C (10 mg, 10% by weight) was weighed into a Schlenk flask and put under argon atmosphere. Compound **7** (100 mg, 0.25 mmol, 1 eq) was dissolved in dry EtOH (10 mL) and added via syringe. The Schlenk flask was then put under hydrogen atmosphere (1 atm), and the suspension was stirred for 16 h. Next the suspension was filtered through celite, and the solvent was evaporated to yield the crude product which was purified by flash chromatography (0–5% MeOH/$CH_2Cl_2$) to give compound **9** as a pale brown solid. Yield: 72% (65 mg); $R_f$ = 0.42 (5% MeOH/$CH_2Cl_2$); **$^1$H NMR** (600 MHz, DMSO-$d_6$) δ 1.39 (s, 9H), 1.50 (s, 6H), 3.73 (s, 3H), 3.78 (s, 3H), 5.02 (s, 2H), 6.19 (d, *J* = 2.3 Hz, 1H), 6.45 (d, *J* = 2.3 Hz, 1H), 6.59 (s, 1H); **$^{13}$C NMR** (151 MHz, DMSO-$d_6$) δ 24.79, 27.56, 35.10, 56.49, 80.05, 81.39, 101.20, 102.70, 117.58, 118.14, 122.47, 143.80, 144.83, 148.74, 158.20, 171.88; **LC-MS** (ESI) (90% $H_2O$ to 100% MeCN in 10 min, then 100% MeCN to 20 min, DAD 220-600 nm), $t_R$ = 5.51 min, 98% purity, *m/z* $[M + H]^+$ calcd for $C_{19}H_{27}N_2O_5$, 363.2; found, 363.4.

*Tert-Butyl 2-((6-((2,5-dichloropyrimidin-4-yl)amino)-8-methoxy-1-methyl-2-oxo-1,2-dihydroquinolin-3-yl)oxy)-2-methylpropanoate* (**11**)

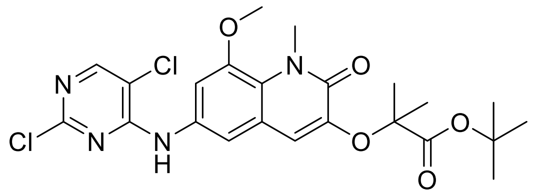

Compound **9** (59 mg, 0.16 mmol, 1 eq), 2,4,5-trichloropyrimidine (35 mg, 0.19 mmol, 1.2 eq) and DIPEA (83 µL, 0.48 mmol, 3 eq) were dissolved in dry DMF (5 mL). The solution was purged with argon for 5 min and stirred at 70 °C overnight. Ater cooling down to RT sat. $NH_4Cl$ (50 mL) was added, and the aqueous phase was extracted with EtOAc (3 × 50 mL). The combined organic layer was washed with 5% LiCl solution (50 mL), sat. NaCl solution (50 mL), dried with $Na_2SO_4$, filtered and evaporated to yield the crude product which was purified by flash chromatography (0–5% MeOH/$CH_2Cl_2$) to give compound **11** as an off-white solid. Yield: 88% (71 mg); $R_f$ = 0.55 (5% MeOH/$CH_2Cl_2$); **$^1$H NMR** (600 MHz, DMSO-$d_6$) δ 1.41 (d, *J* = 1.8 Hz, 9H), 1.54 (d, *J* = 1.9 Hz, 6H), 3.83 (d, *J* = 1.8 Hz, 3H), 3.87 (d, *J* = 1.7 Hz, 3H), 6.75 (s, 1H), 7.44 (d, *J* = 2.4 Hz, 1H), 7.48 (d, *J* = 2.3 Hz, 1H), 8.38 (s, 1H), 9.53 (s, 1H); **$^{13}$C NMR** (151 MHz, DMSO-$d_6$) δ 25.09, 27.94, 35.62, 57.27, 80.62, 82.01, 108.52, 113.86, 114.12, 117.55, 121.53, 124.07, 133.32, 144.42, 148.18, 155.91, 157.28, 157.49, 159.02, 172.01; **LC-MS** (ESI) (90% $H_2O$ to 100% MeCN in 10 min, then 100% MeCN to 20 min, DAD 220-600 nm), $t_R$ = 7.59 min, 97% purity, *m/z* $[M + H]^+$ calcd for $C_{23}H_{27}Cl_2N_4O_5$, 509.1; found, 508.9; **HRMS** (ESI) *m/z* $[M + H]^+$ calcd for $C_{23}H_{27}Cl_2N_4O_5$, 509.1353; found, 509.1352.

*1-(4-((3-((1-(Tert-Butoxy)-2-methyl-1-oxopropan-2-yl)oxy)-8-methoxy-1-methyl-2-oxo-1,2-dihydroquinolin-6-yl)amino)-5-chloropyrimidin-2-yl)piperidine-4-carboxylic acid* (**12**)

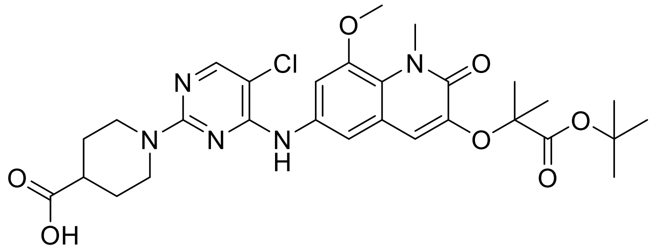

Compound **11** (71 mg, 0.14 mmol, 1 eq) was dissolved in dry DMSO (5 mL). Piperidine-4-carboxylic acid (36 mg, 0.28 mmol, 2 eq) and DIPEA (73 µl, 0.42 mmol, 3 eq) were added and the mixture was stirred at 100 °C overnight. After cooling to RT 10% $KHSO_4$ solution (50 mL) was added, and the aqueous phase was extracted with EtOAc (3 × 50 mL). The combined organic phase was washed with 5% LiCl solution (50 mL), sat. NaCl solution (50 mL), dried over $Na_2SO_4$, filtered and evaporated to yield the crude product which was further purified by flash chromatography (0–10% MeOH/$CH_2Cl_2$) to give compound **12** as a pale brown solid. Yield: 98% (83 mg); $R_f$ = 0.38 (10% MeOH/$CH_2Cl_2$); **$^1$H NMR** (500 MHz, DMSO-$d_6$) δ 1.37 (s, 9H), 1.48 (ddd, *J* = 3.8, 11.3, 12.0 Hz, 2H), 1.54 (s, 6H), 1.84 – 1.91 (m, 2H), 2.50 – 2.57 (m, 1H), 3.03 (ddd, *J* = 2.8,

11.2, 13.7 Hz, 2H), 3.82 (s, 3H), 3.87 (s, 3H), 4.39 (ddd, $J$ = 3.6, 3.6, 14.2 Hz, 2H), 6.72 (s, 1H), 7.53 (d, $J$ = 2.2 Hz, 1H), 7.56 (d, $J$ = 2.3 Hz, 1H), 8.05 (s, 1H), 8.70 (s, 1H), 12.19 (s, 1H); $^{13}$**C NMR** (126 MHz, DMSO-$d_6$) δ 24.70, 27.57, 31.07, 35.23, 40.52, 43.37, 56.77, 80.30, 81.50, 102.22, 107.03, 111.66, 117.24, 121.13, 122.61, 134.57, 144.13, 147.76, 154.79, 155.32, 158.62, 159.23, 171.77, 175.86; **LC-MS** (ESI) (90% $H_2O$ to 100% MeCN in 10 min, then 100% MeCN to 20 min, DAD 220-600 nm), $t_R$ = 5.66 min, 99% purity, $m/z$ [M + H]$^+$ calcd for $C_{29}H_{37}ClN_5O_7$, 602.2; found, 602.5.

*Tert-Butyl 2-((6-((2-(4-((8-(3-(3-amino-6-(2-hydroxyphenyl)pyridazin-4-yl)-3,8-diazabicyclo[3.2.1]octan-8-yl)-8-oxooctyl)carbamoyl)piperidin-1-yl)-5-chloropyrimidin-4-yl)amino)-8-methoxy-1-methyl-2-oxo-1,2-dihydroquinolin-3-yl)oxy)-2-methylpropanoate* (**29**)

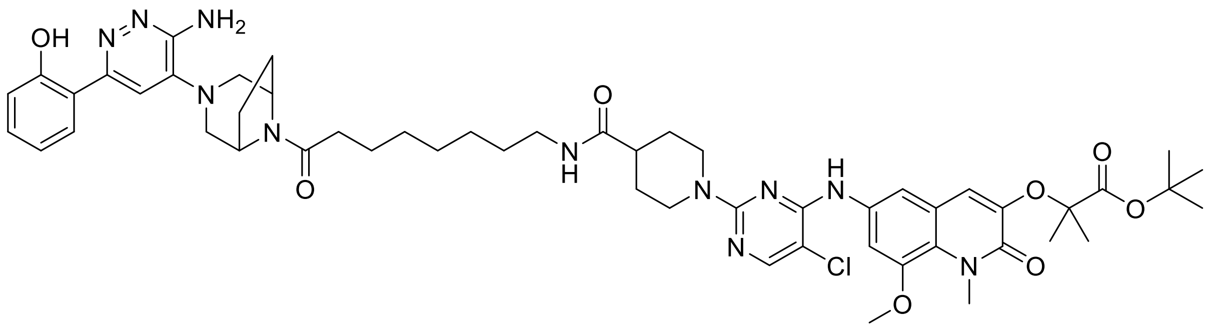

Compound **18** (54 mg, 0.1 mmol, 1 eq) was deprotected according to General Procedure A. Compound **12** (60 mg, 0.1 mmol, 1 eq) and the deprotected amine were used as described in General Procedure C. The reaction was quenched with sat. $NH_4Cl$ (50 mL) and the aqueous phase was extracted with EtOAc (3 × 50 mL). The combined organic layer was washed with 5% LiCl solution (50 mL), sat. NaCl solution (50 mL), dried with $Na_2SO_4$, filtered and evaporated to yield the crude product which was purified by flash chromatography (0–10% MeOH/$CH_2Cl_2$) to give compound **20** as an off-white solid. Yield: 89% (91 mg); $R_f$ = 0.45 (10% MeOH/$CH_2Cl_2$); $^1$**H NMR** (600 MHz, DMSO-$d_6$) δ 1.20 – 1.30 (m, 6H), 1.36 (s, 11H), 1.45 – 1.52 (m, 3H), 1.53 (s, 7H), 1.67 – 1.73 (m, 2H), 1.73 – 1.80 (m, 1H), 1.87 – 1.95 (m, 1H), 2.02 (ddd, $J$ = 4.0, 9.4, 13.5 Hz, 1H), 2.11 (ddd, $J$ = 4.2, 10.9, 12.2 Hz, 1H), 2.26 (dt, $J$ = 7.4, 15.2 Hz, 1H), 2.31 – 2.41 (m, 2H), 2.84 – 2.93 (m, 4H), 3.00 (ddd, $J$ = 6.7, 6.7, 6.7 Hz, 2H), 3.33 – 3.38 (m, 2H), 3.81 (s, 3H), 3.86 (s, 3H), 4.39 (d, $J$ = 6.7 Hz, 1H), 4.52 (d, $J$ = 13.1 Hz, 2H), 4.60 (d, $J$ = 7.0 Hz, 1H), 5.98 (s, 2H), 6.70 (s, 1H), 6.83 – 6.89 (m, 2H), 7.22 (ddd, $J$ = 1.6, 7.3, 8.4 Hz, 1H), 7.53 (d, $J$ = 2.3 Hz, 1H), 7.55 – 7.58 (m, 2H), 7.73 (t, $J$ = 5.6 Hz, 1H), 7.91 (dd, $J$ = 1.7, 8.1 Hz, 1H), 8.04 (s, 1H), 8.70 (s, 1H), 14.08 (s, 1H); $^{13}$**C NMR** (151 MHz, DMSO-$d_6$) δ 24.72, 24.79, 26.40, 27.57, 28.08, 28.17, 28.71, 28.91, 29.23, 31.09, 32.99, 35.25, 38.45, 42.32, 43.72, 50.92, 53.52, 54.05, 54.49, 56.76, 80.27, 81.51, 102.16, 107.00, 111.63, 111.78, 117.17, 117.53, 117.85, 118.61, 121.14, 122.58, 126.44, 130.37, 134.60, 140.55, 144.13, 147.76, 153.79, 154.73, 154.81, 155.33, 158.61, 158.66, 159.23, 168.48, 171.78, 173.92; **LC-MS** (ESI) (90% $H_2O$ to 100% MeCN in 10 min, then 100% MeCN to 20 min, DAD 220-600 nm), $t_R$ = 8.17 min,

99% purity, *m/z* [M + H]$^+$ calcd for $C_{53}H_{69}ClN_{11}O_8$, 1022.5; found, 1023.2; **HRMS** (ESI) *m/z* [M + H]$^+$ calcd for $C_{53}H_{69}ClN_{11}O_8$, 1022.5014; found, 1022.5027.

*2-((6-((2-(4-((8-(3-(3-Amino-6-(2-hydroxyphenyl)pyridazin-4-yl)-3,8-diazabicyclo[3.2.1]octan-8-yl)-8-oxooctyl)carbamoyl)piperidin-1-yl)-5-chloropyrimidin-4-yl)amino)-8-methoxy-1-methyl-2-oxo-1,2-dihydroquinolin-3-yl)oxy)-2-methylpropanoic acid dihydrochloride* (**TRIP1-neg2**)

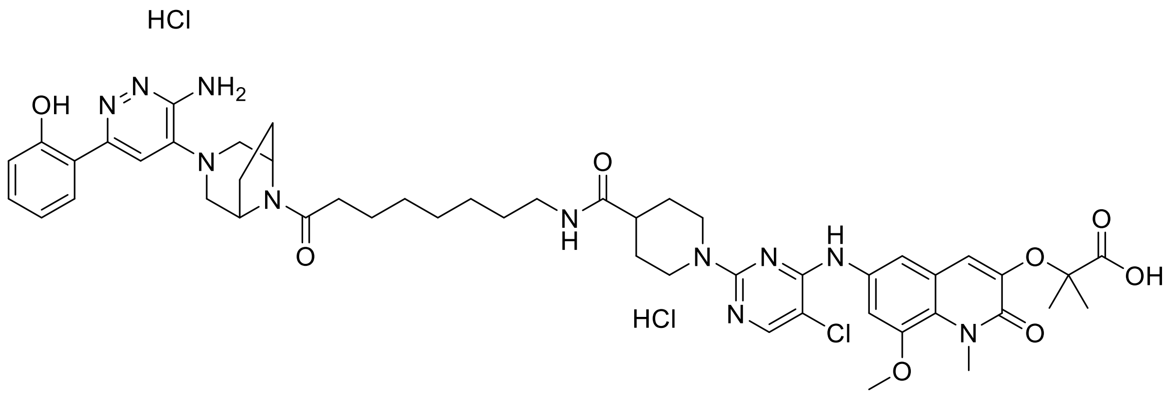

Compound **20** (80 mg, 0.08 mmol, 1 eq) was suspended in 1M HCl in EtOAc (5 mL) and the mixture was stirred at RT for 16 h. The precipitate was then filtered off, washed with dry EtOAc (3 × 30 mL) and dried under vacuum to yield **TRIP1-neg2** as an off-white solid. Yield: 83% (69 mg); **$^1$H NMR** (600 MHz, DMSO-$d_6$) δ 1.19 – 1.30 (m, 6H), 1.32 – 1.40 (m, 2H), 1.43 – 1.63 (m, 10H), 1.72 – 1.79 (m, 3H), 1.86 – 1.98 (m, 2H), 1.98 – 2.04 (m, 1H), 2.22 – 2.30 (m, 1H), 2.31 – 2.37 (m, 1H), 2.39 – 2.47 (m, 2H), 2.96 – 3.10 (m, 5H), 3.16 (d, *J* = 12.5 Hz, 1H), 3.65 (s, 1H), 3.84 (s, 3H), 3.86 (s, 3H), 4.38 – 4.44 (m, 3H), 4.59 (d, *J* = 6.9 Hz, 1H), 6.92 (s, 1H), 6.93 – 6.99 (m, 2H), 7.09 – 7.13 (m, 1H), 7.38 (t, *J* = 7.8 Hz, 1H), 7.47 – 7.55 (m, 4H), 7.84 (t, *J* = 5.6 Hz, 1H), 8.13 (s, 1H), 9.63 (s, 1H); **$^{13}$C NMR** (151 MHz, DMSO-$d_6$) δ 24.79 (d, *J* = 9.4 Hz), 25.82, 26.41, 27.58, 28.01, 28.71, 28.87, 29.20, 32.94, 35.40, 38.50, 41.43, 44.50, 50.86, 53.53, 53.92, 54.56, 56.93, 80.31, 103.22, 108.25, 113.83, 116.73, 118.62, 119.60, 121.27, 123.83, 130.27, 132.58, 133.01, 144.04, 145.11, 147.88, 152.52, 155.61, 156.19, 159.28, 168.73, 172.10, 173.61, 174.24; **LC-MS** (ESI) (90% $H_2O$ to 100% MeCN in 10 min, then 100% MeCN to 20 min, DAD 220-600 nm), $t_R$ = 5.21 min, 94% purity, *m/z* [M + H]$^+$ calcd for $C_{17}H_{22}N$, 966.4; found, 966.6; **HRMS** (ESI) *m/z* [M + H]$^+$ calcd for $C_{17}H_{22}N$, 966.4388; found, 966.4395.

*1-(3-(3-Amino-6-(2-hydroxyphenyl)pyridazin-4-yl)-3,8-diazabicyclo[3.2.1]octan-8-yl)ethan-1-one* (**SMARCA ligand**)

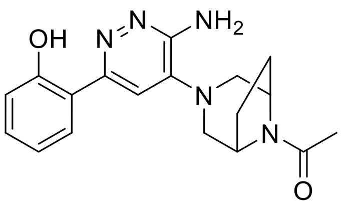

Acetic acid (7 µL, 0.12 mmol, 1 eq) and **SMARCA ligand** (41 mg, 0.12 mmol, 1 eq) were used according to General Procedure C. The reaction was quenched with sat. $NH_4Cl$ (50 mL) and the aqueous phase was extracted with EtOAc (3 × 50 mL). The combined organic layer was washed with 5% LiCl solution (50 mL), sat. NaCl solution (50 mL), dried with $Na_2SO_4$, filtered and evaporated to yield the crude product which was purified by flash chromatography (0–10% MeOH/$CH_2Cl_2$) to give compound **31** as a colorless solid. Yield: 76% (31 mg); $R_f$ = 0.19 (5% MeOH/$CH_2Cl_2$); **$^1$H NMR** (500 MHz, DMSO-$d_6$) δ 1.73 – 1.84 (m, 1H), 1.90 – 2.01 (m, 1H), 2.01 – 2.05 (m, 4H), 2.05 – 2.16 (m, 1H), 2.88 (d, $J$ = 11.3 Hz, 1H), 2.97 (d, $J$ = 11.5 Hz, 1H), 3.23 – 3.29 (m, 1H), 3.36 (dd, $J$ = 2.6, 11.5 Hz, 1H), 4.34 (d, $J$ = 6.7 Hz, 1H), 4.59 (d, $J$ = 6.9 Hz, 1H), 5.99 (s, 2H), 6.84 – 6.91 (m, 2H), 7.20 – 7.27 (m, 1H), 7.58 (s, 1H), 7.89 – 7.94 (m, 1H), 14.07 (s, 1H); **$^{13}$C NMR** (126 MHz, DMSO-$d_6$) δ 21.56, 26.45, 28.00, 50.96, 53.46, 54.38, 54.77, 55.03, 111.87, 117.53, 117.86, 118.63, 126.42, 130.38, 140.55, 153.77, 154.75, 158.65, 165.94; **LC-MS** (ESI) (90% $H_2O$ to 100% MeCN in 10 min, then 100% MeCN to 20 min, DAD 220-600 nm), $t_R$ = 4.67 min, 97% purity, $m/z$ $[M + H]^+$ calcd for $C_{18}H_{22}N_5O_2$, 340.2; found, 340.4.

*tert-butyl 4-(4-(3-(3-amino-6-(2-hydroxyphenyl)pyridazin-4-yl)-3,8-diazabicyclo[3.2.1]octan-8-yl)-4-oxobutyl)piperidine-1-carboxylate* (**22**)

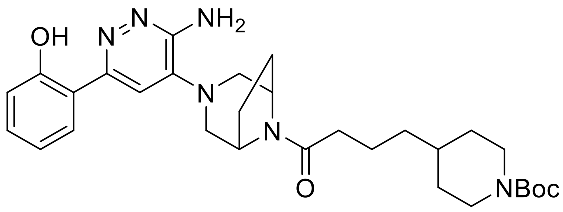

4-Piperidinebutanoic acid (81 m g, 0.3 mmol, 1 eq) and Compound **17** (115 mg, 0.3 mmol, 1 eq) were used according to General Procedure C. The reaction was quenched with sat. $NH_4Cl$ (50 mL) and the aqueous phase was extracted with EtOAc (3 × 50 mL). The combined organic layer was washed with 5% LiCl solution (50 mL), sat. NaCl solution (50 mL), dried with $Na_2SO_4$, filtered and evaporated to yield the crude product which was purified by flash chromatography (0–5% MeOH/$CH_2Cl_2$) to give compound **22** as a pale yellow solid. Yield: 89% (147 mg); $R_f$ = 0.53 (5% MeOH/$CH_2Cl_2$); **$^1$H NMR** (600 MHz, DMSO-$d_6$) δ 0.93 (dt, J = 4.3, 12.5 Hz, 2H), 1.18 – 1.26 (m, 2H), 1.33 – 1.42 (m, 10H), 1.49 – 1.58 (m, 2H), 1.58 – 1.65 (m, 2H), 1.77 (ddd, J = 4.8, 9.7, 12.5 Hz, 1H), 1.91 (ddd, J = 4.7, 8.2, 16.1 Hz, 1H), 2.03 (ddd, J = 4.2, 9.5, 13.4 Hz, 1H), 2.12 (ddd, J = 4.3, 9.3, 11.8 Hz, 1H), 2.23 – 2.31 (m, 1H), 2.31 – 2.39 (m, 1H), 2.55 – 2.75 (m, 2H), 2.85 – 2.93 (m, 2H), 3.26 – 3.29 (m, 1H), 3.34 – 3.40 (m, 1H), 3.89 (d, J =

12.5 Hz, 2H), 4.39 (d, J = 6.8 Hz, 1H), 4.61 (d, J = 7.0 Hz, 1H), 5.99 (s, 2H), 6.84 – 6.90 (m, 2H), 7.23 (ddd, J = 1.6, 7.2, 8.4 Hz, 1H), 7.56 (s, 1H), 7.92 (dd, J = 1.6, 8.0 Hz, 1H), 14.08 (s, 1H); **$^{13}$C NMR** (151 MHz, DMSO-$d_6$) δ 21.93, 26.37, 28.09, 28.25, 31.91, 33.13, 35.20, 35.84, 38.39, 50.94, 53.55, 54.03, 54.48, 78.49, 111.78, 117.54, 117.86, 118.62, 126.44, 130.38, 140.55, 153.78, 154.02, 154.73, 158.66, 168.40; **LC-MS** (ESI) (90% $H_2O$ to 100% MeCN in 10 min, then 100% MeCN to 20 min, DAD 220-600 nm), $t_R$ = 7.49 min, 98% purity, *m*/*z* $[M + H]^+$ calcd for $C_{30}H_{43}N_6O_4$, 551,3; found, 551.7;
